# Triplet-encoded prebiotic RNA aminoacylation

**DOI:** 10.1101/2023.03.13.532408

**Authors:** Meng Su, Christian Schmitt, Ziwei Liu, Samuel J. Roberts, Kim C. Liu, Konstantin Röder, Andres Jäschke, David J. Wales, John D. Sutherland

**Author notes:** Randall Centre for Cell & Molecular Biophysics, King’s College London; London, WC2R 2LS, UK. These authors contributed equally (Meng Su, Christian Schmitt).

## Abstract

The encoding step of translation involves attachment of amino acids to cognate tRNAs by aminoacyl-tRNA synthetases, themselves the product of coded peptide synthesis. So, the question arises — before these enzymes evolved, how were primordial tRNAs selectively aminoacylated? Here we demonstrate enzyme-free, sequence-dependent, chemoselective aminoacylation of RNA. We investigated two potentially prebiotic routes to aminoacyl-tRNA acceptor stem-overhang mimics and analyzed those oligonucleotides undergoing the most efficient aminoacylation. Overhang sequences do not influence the chemoselectivity of aminoacylation by either route. For aminoacyl-transfer from a mixed anhydride donor strand, the chemoselectivity and stereoselectivity of aminoacylation depends on the terminal three base pairs of the stem. The results support early suggestions of a second genetic code in the acceptor stem.

**One-Sentence Summary:** Selectivity of RNA stem-overhang aminoacylation is controlled by the terminal trinucleotide sequence of the stem.

## Introduction

The translation of genetic information contained in mRNA into specific protein sequences according to the genetic code depends on two molecular recognition events: first, attachment of specific amino acids to cognate tRNAs; and second, binding of charged tRNAs to mRNA. The latter depends primarily on anticodon:codon binding mediated by the nucleic acids themselves through Watson-Crick base pairing, with help from the decoding centre of the ribosome. The attachment of amino acids to cognate tRNAs, on the other hand, does not depend on amino acid:RNA interactions, but has an obligate requirement for enzyme control. Specific aminoacyl-tRNA synthetases (aa-tRNA synthetases) recognize both amino acids and cognate tRNAs and catalyze their joining together in an ATP-consuming reaction (*1*). Two questions relating to the origin of the translation thus arise: ‘How could specific aminoacylation of cognate tRNAs have been achieved without enzymes?’ and ‘On what basis were amino acid:codon assignments initially made?’.

Limited experimental work bearing on these questions has left answers mainly in the realm of conjecture (*2*). The role of stereochemical interaction between amino acids and tRNA in setting the genetic code has been considered by several authors with opinions varying as to where the tRNA residues involved in the recognition reside (*3-6*). In principle, such stereochemical interactions could answer both questions at once, if amino acid:anticodon interactions simultaneously controlled both specificity of aminoacylation and amino acid:codon assignment. However, there has been no clear experimental support for this assumption for many decades, so the possibility of stereochemical interactions with RNA residues other than the anticodon remains open, but it is difficult to see how this could have affected codon assignment (*7*). That RNA enzymes (ribozymes) can catalyze uncoded aminoacylations and also peptide couplings in the absence of proteins has been amply demonstrated (*8-12*). However, these systems do not allow conclusions to be drawn about codon-amino acid assignments. The ‘frozen accident theory’ has it that assignments were made at random and then became fixed to maintain genotype:phenotype integrity (*13*).

Alternatively, it has been argued that assignments were made to minimize the phenotypic effect of coding errors or were made sequentially as new amino acids became available through biosynthesis (*14*).

The realization that tRNA identity determinants — used by aminoacyl-tRNA synthetases to determine whether a tRNA is cognate — do not always include the anticodon, resulted in a major conceptual advance (*15, 16*). It was found that many identity determinants are clustered in the tRNA acceptor stem (*17*), indeed in one case identity is determined by a single base pair in the acceptor stem (*18-20*). This led to the suggestion that a ‘second genetic code’ is written in the acceptor stem of tRNA and read by aa-tRNA synthetases (*21*). It was speculated that this ‘still largely undeciphered’ second code might be older and more deterministic than the classical genetic code; possibly even depending on stereochemical interactions between a particular sequence in the acceptor stem and the cognate amino acid or aminoacyl-intermediate (*22-24*). Short RNA molecules (aptamers) selected from random sequence mixtures by amino acid binding have been reported to be enriched with cognate triplets for the respective amino acids (*25*), but the studies have not been extended to chemical reactions of the bound amino acids. In the absence of enzymes, production of selectively aminoacylated tRNAs would require aminoacylation to somehow be coded by RNA sequence.

Several years ago, we uncovered a ‘protometabolic’ network of reactions based on the reductive homologation of hydrogen cyanide and its derivatives by hydrogen sulfide under photochemical conditions (*26*). This ‘cyanosulfidic’ chemistry led to precursors of nucleotides as well as amino acids suggesting that the assembly into higher-order structures occurred in mixtures of these building blocks. The activated nucleotides and oligonucleotides required to assemble RNA by sequential monomer addition and ligation, respectively, are known to undergo competing reactions with amino acids that depend on pH (*27*). In a mildly alkaline solution, reaction of the free amino groups with activated nucleotides results in phosphoramidates, whereas under slightly acidic conditions, where amino groups are substantially protonated, the carboxylate groups attack instead, affording mixed carboxylic-phosphoric anhydrides (*28-30*) (Fig. 1A). We recently discovered efficient chemistries whereby an amino acid is transferred from either sort of RNA:amino acid conjugate at the 5′-phosphate of a tRNA acceptor stem mimic to the 2′,3′-diol terminus of a short 3′-overhang (*31, 32*) (Fig. 1B). In light of the aforementioned suggestions of a second genetic code, we wondered if there might be a relationship between the sequence of the acceptor stem-overhang and which amino acid is most efficiently transferred by one or other chemistry.

**Fig. 1.**
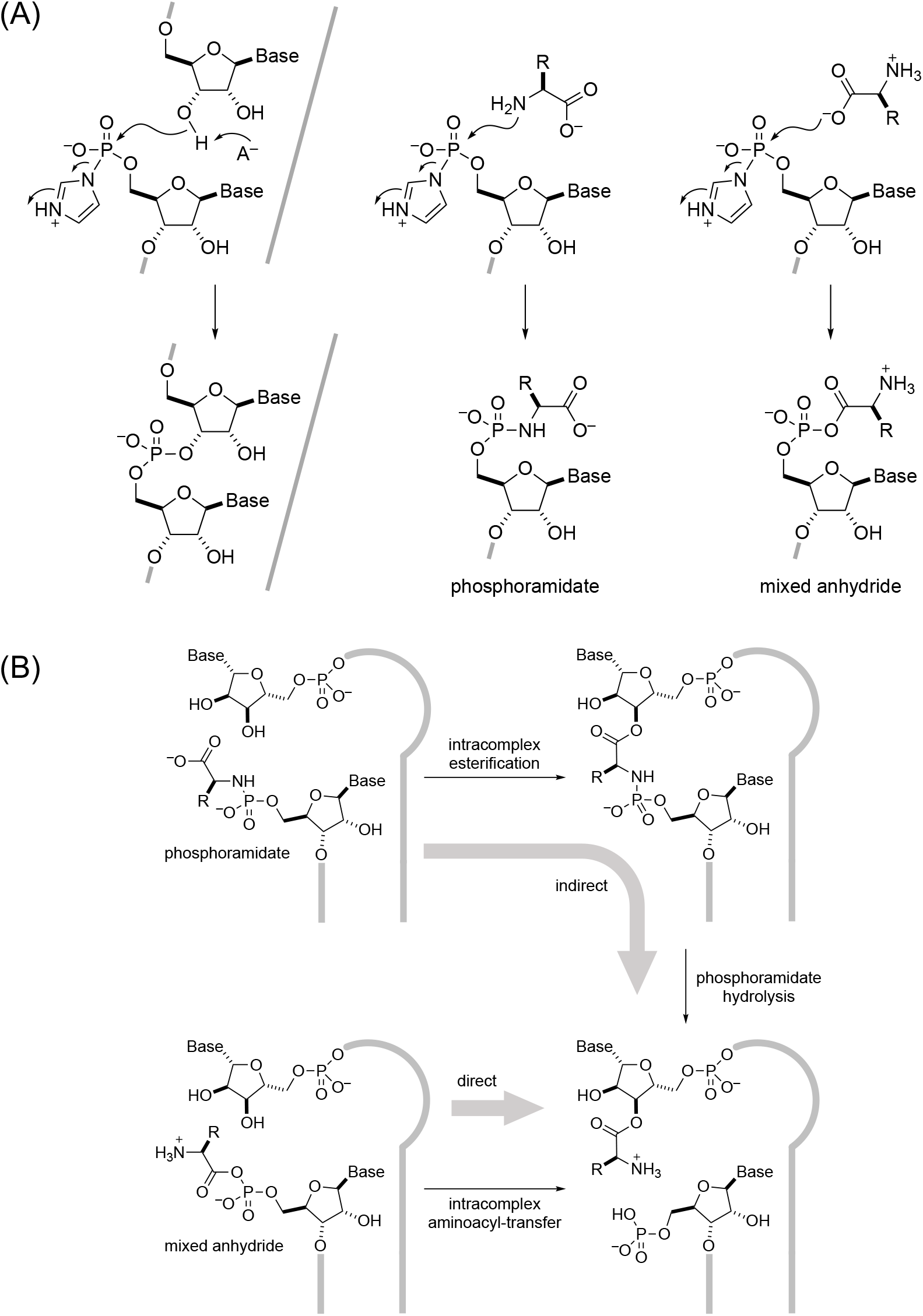
RNA aminoacylation via activated phosphates. (**A**) Activated phosphates required for RNA ligation chemistry also react with amino acids to give phosphoramidates and/or mixed anhydrides depending on pH; (**B**) production of 3′-aminoacyl-RNA by indirect and direct transfer of aminoacyl groups from phosphoramidates and mixed anhydrides respectively.

## Results and discussion

Based on our earlier work on mixed anhydride chemistry, we varied the nature of the 3′-overhang attached to an acceptor stem duplex mimic and used HPLC to monitor the efficiency of direct inter-strand aminoacyl-transfer. Through a very sparse sampling of aminoacyl residues and overhang length and sequence (Table S1 and S2), we found that the five base overhang 5′-UUCCA-3′ (overhang sequences underlined) allowed the most efficient and stereoselective (L-preferred over D-)alanyl-transfer. We then used this same overhang sequence to investigate stereoselectivity and the effect of changing amino acids on indirect aminoacyl-transfer starting from phosphoramidates and proceeding via phosphoramidate-ester intermediates. Stereoselectivity (L-preferred over D-) was maintained for this indirect aminoacyl-transfer across a range of amino acids, but yields were amino acid-dependent (Table S3). To extend our earlier investigations, we fully randomized a five-base overhang (Table S4) and selected those sequences best able to undergo direct or indirect aminoacyl-transfer using either chemistry at the 5′-terminus of a tRNA acceptor stem mimic. Within the subset of amino acids that can be made by cyanosulfidic chemistry (*26*), for ease of synthesis we further restricted ourselves to making glycyl-, L-/D-alanyl, L-prolyl-, L-leucyl-and L-valyl-mixed anhydrides and phosphoramidates of 5′-pAGCGA-3′ and separately subjected them to aminoacyl-transfer to an annealed pool of oligonucleotides with the 3′-terminal decanucleotide sequence: 5′-UCGCU<u>NNNNN</u>-3′. In the case of indirect aminoacyl-transfer, we had previously shown the second step (mild acid hydrolysis, Fig. 1B) proceeds in uniformly high yield with the amino acids we had chosen (*32*). For this reason, as well as for experimental practicality, we did not hydrolyse the phosphoramidate P–N bond to form an aminoacyl-ester in the current work.

To determine which sequences had undergone the most efficient direct aminoacyl-transfer, or phosphoramidate-ester formation with the two chemistries, we developed a high-throughput screen strategy (Fig. S1). Acylation of an RNA 2′,3′-diol terminus protects it against oxidation by periodate whereas free 2′,3′-diol termini are oxidized to non-ligatable dialdehyde termini. Subsequent hydrolysis of aminoacyl-ester termini and bridged phosphoramidate-esters allowed ligation of the newly liberated diol termini to an adapter oligonucleotide complementary to a reverse transcription primer. Successful ligation enabled reverse transcription and PCR amplification using primers adapted to multiplex sequencing (Table S4). Next-generation sequencing of the amplified sequence mixture then allowed us to find those overhang sequence variants that gave the most efficient aminoacyl-transfer or phosphoramidate-ester formation (by analyzing the number of sequencing reads for all 1024 pentanucleotides) and to rank them for each amino acid. As a blank, we subjected the partially randomized oligonucleotide pool only to the ligation, reverse transcription and PCR amplification and sequenced it to reveal those variants favoured by this sample processing procedure. The tabulated data for the aminoacyl-transfer and phosphoramidate-ester formation experiments and the blank were visualised to display the sequence preferences in a graphical format and compared them in scatter plots to discern whether the preferred sequences for one aminoacyl residue were systematically related to those of any other (Fig. S2). For direct aminoacyl-transfer from mixed anhydride RNA:amino acid conjugates the results revealed that different aminoacyl residues are transferred to a similar subset of U, G-rich overhang sequences and that this subset of sequences is different from the U,A-rich subset that is favoured by the blank sample processing procedure. This demonstrates that certain overhang sequences enable more efficient aminoacyl-transfer than others. However, crucially, there appears to be no idiosyncrasy with respect to the transfer of specific aminoacyl residues. We found that the 5′-<u>UUCCA</u>-3′ sequence (previously determined from sparse sampling to undergo efficient L-alanyl-transfer) was among the top 10% of sequences for all amino acids tested, and in each case was the best among sequences ending in the 5′-CCA-3′ trinucleotide sequence (Table S5 and S6). For phosphoramidate-ester formation, A-deficient overhang sequences were preferred across the range of amino acids whilst the blank preference was U, A-rich. Thus neither chemistry was associated with significant selective preference of different overhang sequences for different amino acids — no hints of coding were apparent. Our attention was thus switched to the stem region. We stuck with 5′-<u>UUCCA</u>-3′ because it was favoured in the mixed anhydride chemistry, reasonably efficient in the phosphoramidate-ester chemistry, and close to the canonical tRNA overhang sequence.

We focused on the effect of stem sequence on the mixed anhydride chemistry first. Unsure of how many stem residues to randomize, we synthesized a range of oligonucleotide donor-acceptor pairings with base pair changes at specific positions to delineate the region of the stem, if any, affecting the specificity of aminoacyl-transfer. These oligonucleotide combinations were then analysed in an HPLC-based kinetic assay (Table S1, S7-S12, Ref (*31*)). In this way, we found that the three base pairs of the stem proximal to the overhang influenced aminoacyl-transfer (Fig. 2A, C-F) whereas changing the more distal fourth base pair had no effect (Fig. 2B). We, therefore, randomized the three residues upstream of the 5′-<u>UUCCA</u>-3′ overhang of an acceptor oligonucleotide (3′-terminal tridecanucleotide sequence: 5′-GAUUCNNN<u>UUCCA</u>-3′) to be used with a randomized octanucleotide donor (5′-pNNNGAAUC-3′, Fig. S3). After annealing of separate aminoacyl-phosphate mixed anhydrides of the donor oligonucleotide pool randomized in the three 5′-residues, we submitted the samples to aminoacyl-transfer conditions and sequenced those acceptor strands that were protected from periodate oxidation by aminoacyl-transfer. Again, we performed a blank in which solely the partly randomized acceptor oligonucleotide was subjected to sample processing. Sequencing revealed that for some amino acids, those sequence variants that underwent the most efficient aminoacyl-transfer were very different, whereas for others there were similarities (Fig. 2G, Table S13-S15). Thus, for example, the preferred sequences for L-alanyl-, D-alanyl-and glycyl-transfer were very different from each other and from the sequences preferred for transfer of other aminoacyl residues (as apparent from the comparison of the graphic representation of the top 10% of reads and from scatter plots). On the other hand, there was a clear similarity between those sequences preferred for L-valyl-and L-leucyl-transfer and a lesser similarity between those sequences and the sequences preferred for L-prolyl-transfer. Glycyl-transfer stood out in the sense that the results were dominated by a single sequence (5′-CUC-3′) which appeared in 44% of reads. Ignoring this highly represented sequence, the remaining preferences were still different from those for transfer of other aminoacyl residues (Fig. S4). In no case did the trinucleotide preferences for transfer of any particular aminoacyl residue bear any obvious resemblance to the extant codon or anticodon sequences for the corresponding amino acid. So, it appears unlikely that this stereochemical coding is connected to codon assignment. However, it is consistent with a ‘second genetic code’ in the acceptor stem (*21*). This second code relates to the specificity of tRNA aminoacylation, but not directly to codon assignment.

**Fig. 2.**
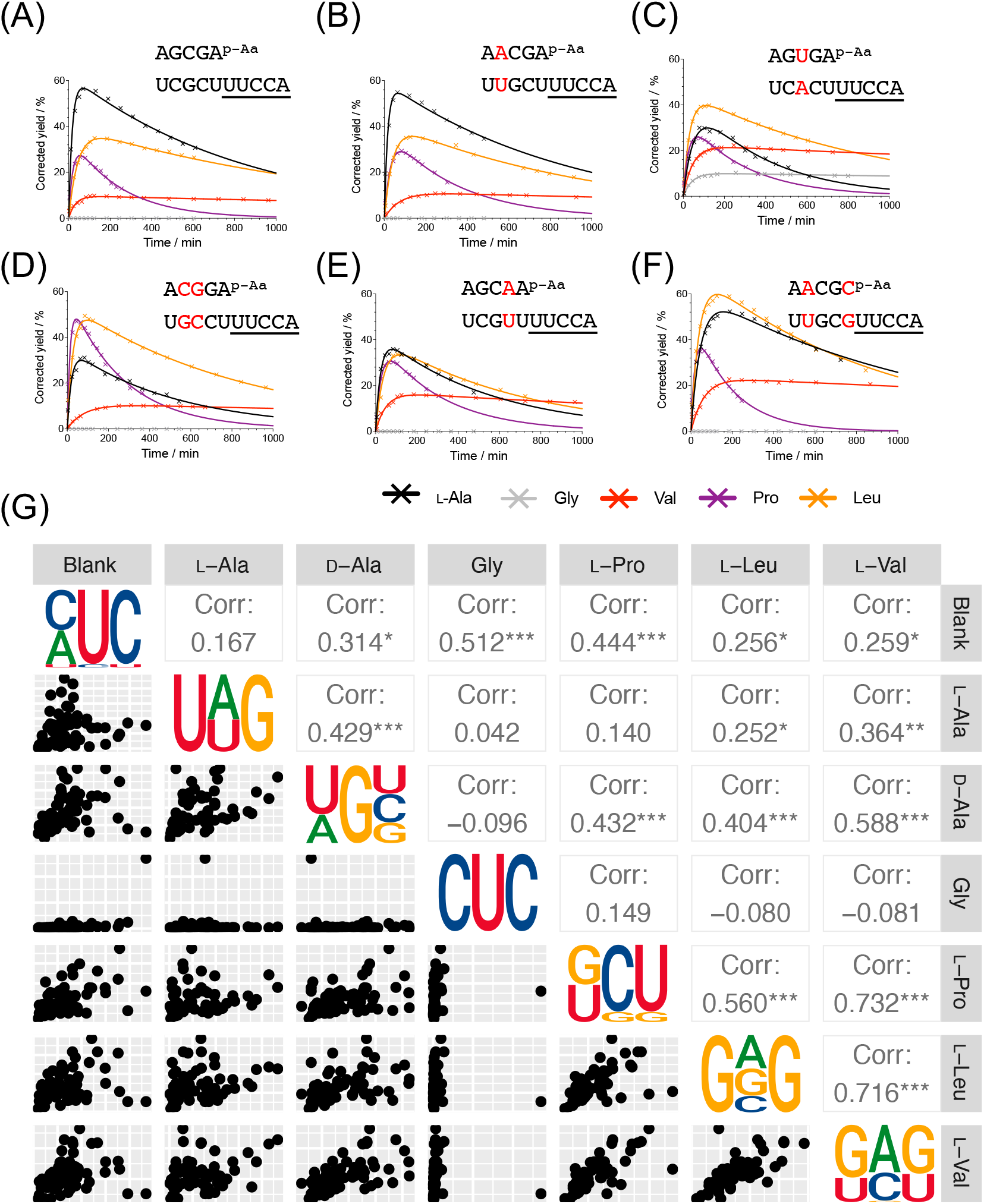
Divergent pattenrs in the first three nucleotides in the stem. (**A**-**F**) Time courses showing the aminoacyl-transfer kinetics with identical overhang ‘UUCCA’, but base pairs changed in the stem; (**G**) Plot of sequencing results after mixed anhydride aminoacylation selection using a partially randomized stem (5′-GAUUCNNN<u>UUCCA</u>) showing correlation between amino acids. Sequence logo represents the top 10% of trinucleotides from the raw reading numbers. Pearson correlation coefficients are shown in the upper right corner. Aminoacyl-transfer conditions: both oligos (100 μM), NaCl (100 mM), MgCl_2_ (5 mM), HEPES (50 mM, pH 6.8).

We then investigated the effect of the stem sequence on the phosphoramidate-ester chemistry. To allow comparison to the results for the mixed anhydride chemistry, we randomized the same trinucleotide region of the stem and separately annealed portions of this library to specific amino acid phosphoramidates of the complementary randomized donor strand (Fig. S5). After subjecting the various acceptor stem-overhang combinations to esterification conditions, we then used the same procedure that we had used before. In contrast to what we had seen with the mixed anhydride chemistry, sequencing revealed that the nature of the amino acid had almost no effect on the preferred sequences for phosphoramidate-ester formation although the consensus sequence preference was distinctly different from the preferences seen for the blank sample (Fig. S6, Table S16 and S17). We felt that the complete lack of amino acid side chain influence on the preferred stem sequences for phosphoramidate-ester formation made the idiosyncrasies we had seen with the mixed anhydride chemistry even more remarkable. Accordingly, we now abandoned the indirect aminoacyl-transfer chemistry and focused exclusively on the direct transfer chemistry.

The selection and sequencing protocol cannot be expected to give quantitative data about the kinetics of aminoacyl-transfer. Accordingly, we used the results to guide our choice of sequences for the determination of the kinetics of individual aminoacyl-transfer reactions by HPLC. As sequencing only determined the acceptor strand, for the kinetic analysis we paired it with its Watson-Crick complement in most cases. The first such pairing was based on a trinucleotide combination that the sequencing data suggested should give efficient and selective aminoacylation with L-Ala (acceptor 5′-CAG-3′ and its complement 5′-CUG-3′).

Indeed, L-Ala was more rapidly and completely converted in the HPLC assay than any of the other amino acids tested (Fig. 3A, Table S18).

**Fig. 3.**
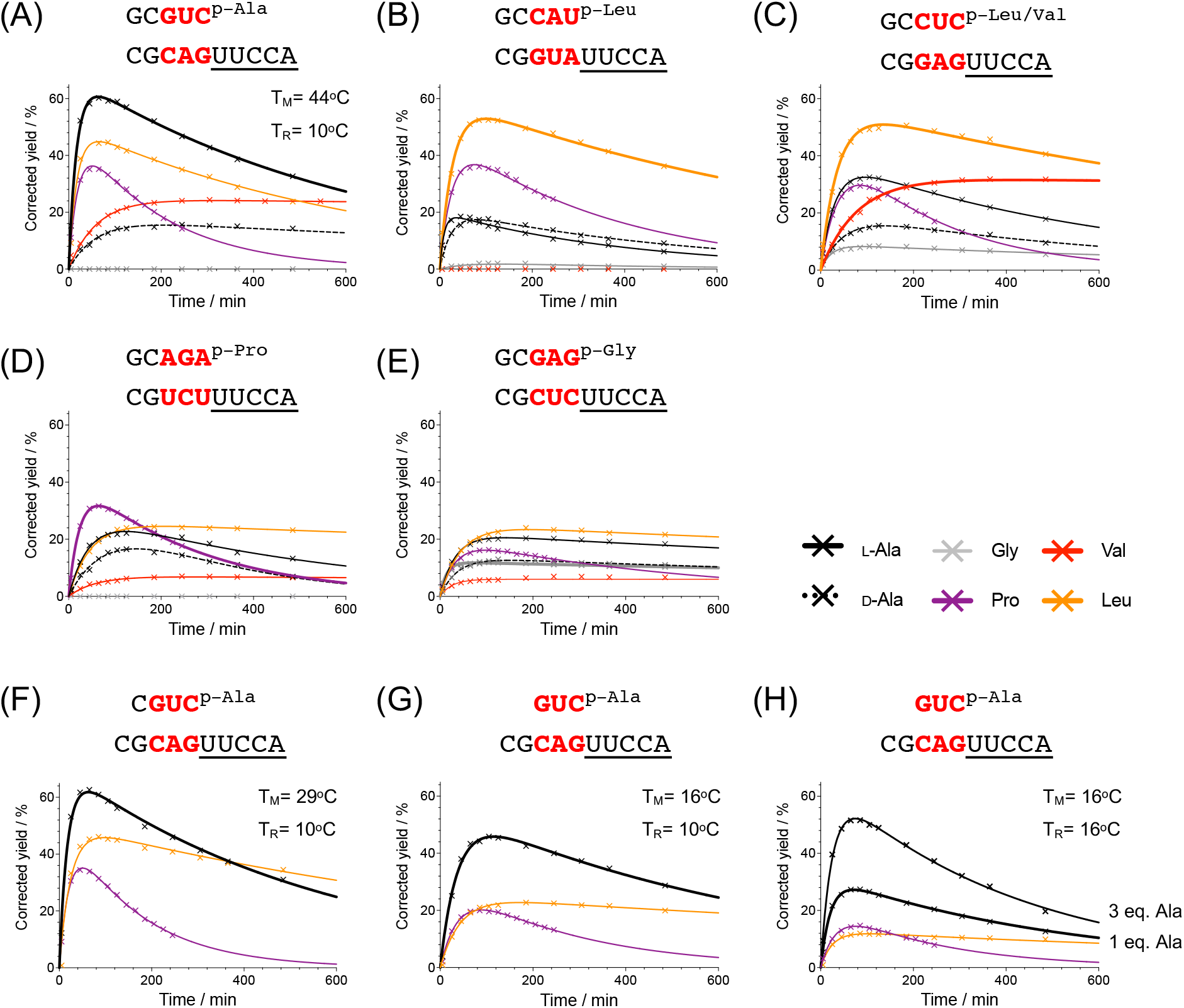
Stereochemical coding dictated by stem-terminal sequence. Time courses showing the mixed anhydride aminoacyl-transfer kinetics with varied trinucleotide stem sequences according to the top selectivity score (**A**) L-Ala; (**B**) L-Leu; (**C**) L-Leu and L-Val; (**D**) L-Pro; (**E**) Gly, (**A**-**E**) all with 1 eq. donor transferred at 10°C; (**F**) 1 eq. tetramer donor for L-Ala at 10°C; (**G**) 1 eq. trimer donor for L-Ala at 10°C; (**H**) 1 or 3 eq. trimer donor for L-Ala at 16°C. T_M_, melting temperature; T_R_, temperature at which the aminoacyl-transfer reaction was conducted. Conditions: both oligos (100 μM), NaCl (100 mM), MgCl_2_ (5 mM), HEPES (50 mM, pH 6.8).

Remarkably, glycyl-transfer between the same oligonucleotides was undetectable. The stereoselectivity of alanyl-transfer was high (L-preferred over D-by a factor of 3.5, Table S18), and although L-prolyl, L-valyl and L-leucyl residues were also transferred fairly well, product yields were not as high as they were with L-alanine. The principal alanyl-tRNA identity determinant in extant biology is the G3:U70 wobble base pair (although a C3:G70 base pair is also functional) (*18, 19, 33*). We investigated the effect of a similar wobble base pair on L-alanyl-transfer in our system by keeping the same donor strand and changing the acceptor to one terminating in the sequence 5′-UAG-3′ (Table S18). The change to a wobble base pair slowed down L-alanyl-transfer and almost halved the maximum yield of the transfer product.

According to the sequencing data, the general trinucleotide preferences for L-leucyl and L-valyl transfer are similar. However, the kinetic analysis of the transfer of these more hydrophobic amino acids revealed that the 5′-GUA-3′ trinucleotide is efficient in the transfer of leucine but inefficient in the transfer of valine (Fig. 3B, Table S19). In contrast, the trinucleotide 5′-GAG-3′, which sequencing had suggested as the consensus sequence for both L-leucyl-and L-valyl-transfer, transfers both amino acids efficiently. In this system, glycyl transfer is now observed, albeit in a lower yield (Fig. 3C, Table S20). The top-ranked sequence for proline transfer, 5′-UCU-3′, was found to transfer L-proline more efficiently than all other amino acids in the HPLC assay (Fig. 3D, Table S21). The sequencing data further suggest that L-valyl-transfer should be more efficient than L-leucyl-transfer with the sequence 5′-GCU-3′ (Table S14). However, we had already investigated that sequence when we were delineating which part of the stem affects aminoacyl-transfer — albeit with a stem differing at the fourth and fifth positions — and found the opposite to be the case (Fig. 2A and B). Thus, as is generally held in the field although the sequencing data can serve as a guide to discern selectivity, it cannot be relied upon to give quantitative data.

For many stem-terminal trinucleotide sequences, glycyl-transfer was barely detectable. However, as mentioned above, low yield transfer was apparent for the trinucleotide 5′-GAG-3′ (Fig. 3C). Using the top-ranked trinucleotide 5′-CUC-3′, glycyl-transfer was somewhat more efficient, but was still less efficient than L-leucyl-and L-prolyl-and alanyl-transfer (Fig. 3E, Table S22). If the abundances of prebiotic amino acids on early Earth were largely equal, then the implication is that selectively glycylated tRNAs could not likely have been produced by stereochemical coding. However, given its simplicity, glycine was the most abundant prebiotic amino acid (*26, 34, 35*). If its concentration was significantly higher than any of the other amino acids, then stem-overhang pairings based on the trinucleotide 5′-CUC-3′ could have been selectively glycylated whilst glycylation of other stem-overhangs might have been less efficient than their aminoacylation with other amino acids. Thus, with a restricted subset of amino acids, stereochemical coding dictated by the stem-terminal trinucleotide sequence is possible. Furthermore, high stereoselectivity (L-preferred over D-) was also imposed on alanyl-transfer using the stem-terminal trinucleotide 5′-CAG-3′ and its complement 5′-CUG-3′ (Fig. 3A). Several other oligonucleotide donor-acceptor pairings were associated with similarly stereoselective alanyl-transfer (e.g. Fig. 3C), but the donor-acceptor pair that underwent efficient L-leucyl-transfer (stem-terminal trinucleotide 5′-GUA-3′ and its complement 5′-UAC-3′, Fig. 3B) displayed almost no stereoselectivity for alanyl-transfer. Stereoselectivity, like amino acid chemoselectivity, is thus controlled by the stem-terminal trinucleotide sequence.

In extant biology, tRNAs are aminoacylated using aminoacyl-adenylates which are much smaller than the donor oligonucleotides used here. Therefore, we next investigated if stereochemical coding is maintained in our system when the length of the oligonucleotide donor is progressively reduced. For the duplex of a nonaminoacylated pentanucleotide donor and a decanucleotide acceptor (5′-CGCAGUUCCA-3′), we determined a melting temperature T_M_ of 44°C. This value is well above the 10°C temperature at which we carried out the aminoacyl-transfer reactions, resulting in the formation of a stable duplex under the assay conditions (Fig. 3A and Fig. S7). Truncating the donor to a tetranucleotide while keeping the acceptor constant lowered the T_M_ to 29°C and the aminoacyl-transfer efficiency remained almost unchanged for the three amino acids studied (compare Fig. 3A and F). Using a trinucleotide donor with the T_M_ of 16°C reduced the efficiency of transfer of all three amino acids, but increased in the selectivity of coding for L-alanyl-transfer (Fig. 3G and Table S23). At a T_M_ so close to the reaction temperature, a significant portion of the strands are present dissociated, and we found that the yield of the aminoacylated acceptor strand was markedly increased when we used three equivalents of the L-alanyl donor strand (Fig. 3H). Ribozyme auxiliaries could then improve the aminoacylation selectivity resulting in better coding and progressively allow the use of aminoacylated di- and mono-nucleotides. Thus, a series of small steps can be envisaged to take the chemistry that we have uncovered here to the point where coded peptides could start to contribute to the efficiency of their own generation through becoming aminoacyl-tRNA synthetases.

While there is clear evidence for sequence and aminoacyl side chain dependence in the transfer selectivity, it is not clear how this selectivity arises. One possibility is that differences in the structures adopted by the different sequences and aminoacyl groups may play a key role in determining the observed trends. To investigate this possibility further, we conducted simulations exploring the underlying energy landscapes to identify whether such structural changes exist. The first set of calculations focused on two distinct sequences each for L-alanyl and L-leucyl transfer, one with a high, and one with a lower, yield and *k*_*transfer*_ (Fig. S8 and Table S24, S25). We found that the structural ensembles indeed exhibit distinct characteristics. In both cases, the lower-yielding sequence (Fig. S8B and D) makes many close contacts between the amino acid and atoms of the oligonucleotide duplex (red spikes in the energy landscapes) that the higher-yielding counterpart (Fig. S8A and C) does not make. These interactions are either between the Hoogsteen edges of the second and third base pair with either the phosphate group linking the aminoacyl group to the nucleic acid (Fig. S8 snapshot B1and B2) or the amino group of the amino acid (Fig. S8 snapshot D1-3). The selectivity in these scenarios arises for the first set (Fig. S8A and B) of interactions from the nature of the Hoogsteen edge (an available electron-rich O or N for the amide or a OH or NH for the phosphate), and in the second case from the ability to rearrange the backbone when there is weaker base pairing at the top of the stem (A:U vs C:G, Fig. S8C and D). These contacts make a close approach of the overhanging 3′-end of the acceptor strand to the aminoacyl group geometrically challenging, suggesting an energy barrier for the transfer, as these contacts need to be lost first. This effect likely lowers the yield, but the contacts are not strong enough to completely suppress the reactions.

Once we had diagnosed a structural basis for the observed difference in yields, a second set of simulations focused on the systems in Fig. 3A and 3B to determine whether the stereoselectivity for alanyl-transfer, the difference in yields for L-valyl- and L-leucyl-transfer, and the low yield for glycyl-transfer can also be linked to structural features (Fig. 4 and Fig. S9). For the sequence from Fig. 3A, the top base pair is G:C and always intact, while for the systems from Fig. 3B, the A:U base pair is absent in the case of valine, alanine and glycine.

**Fig. 4.**
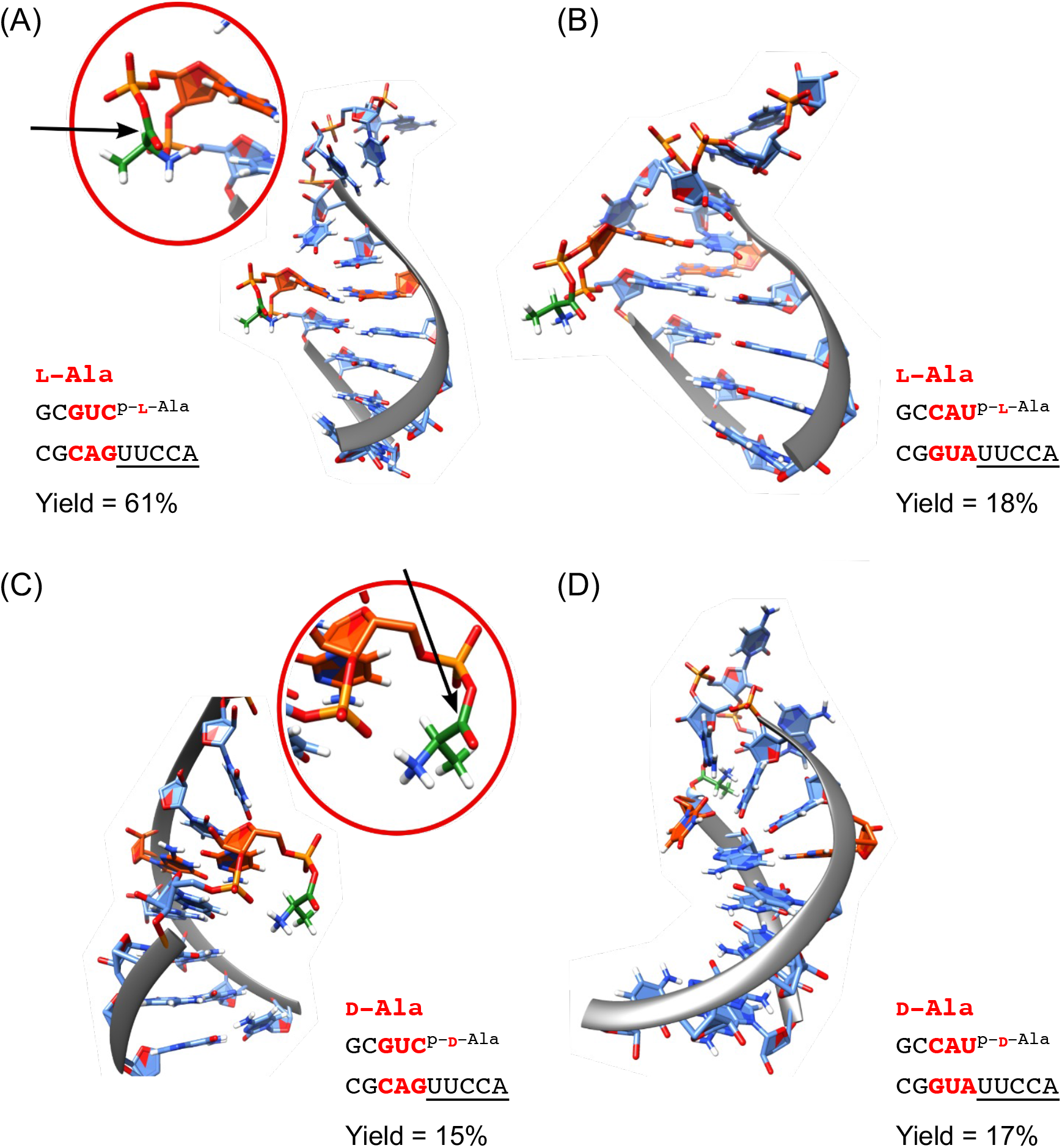
Representative structures for L-Ala (**A** and **B**) and D-Ala systems (**C** and **D**), selected from the energy landscape databases. Comparing **A** and **C** suggests an explanation for the observed stereoselectivity. For L-Ala (**A**) the Burgi-Dunitz trajectory is freely accessible (black arrow). In contrast, the trajectory is blocked in D-Ala (**C**). For **B** and **D**, the top base pair in the stem is frayed in both cases, giving more flexible structures. The aminoacyl moiety is highlighted in green and the top pair of bases in the stem is highlighted in orange.

For the L-valyl-mixed anhydride, the loss of base pairing leads to strong interactions of the aminoacyl group and the distorted stack, which forms a triplet. As a result, the overhang is not able to achieve the proximity required for the transfer (Fig. S9C, E, and F). In contrast, the L-leucyl-stem-overhang exhibits no such distortions and is accessible for the transfer reaction (Fig. S9A, B, and E). We speculate that the difference in behaviour arises from the side chain length, and the increased hydrophobicity, as the glycyl-, alanyl- and valyl-stem-overhangs all exhibit the change in the stem, but the leucyl-stem-overhang does not.

The stereoselectivity between L- and D-alanyl-transfer is also linked to the stability of the top base pair in the stem. A stable base pair allows interactions of the alanyl residue with the stack, but in such a fashion that the 3′-overhang is able to get in close proximity.

However, while the L-stereoisomer enables access to the carbonyl carbon along the Burgi-Dunitz trajectory, the D-stereoisomer, as the interactions with the nucleobases are the same, is flipped and the approach to the carbonyl is blocked by the stem (Fig. 4A and C). When the top base pair is lost, these configurations are no longer observed, and the selection bias disappears (Fig. 4B and D).

Finally, the glycyl residue in all cases exhibits strong interactions, likely stabilized by the absence of a hydrophobic side chain.

## Conclusions

The attractiveness of tRNA self-aminoacylation as a prelude to coded translation was alluded to over fifty years ago (*12*). Our finding that triplet-encoded chemo- and stereoselective tRNA acceptor stem-overhang mimic aminoacylation is possible and thus provides the firstexperimental support for these earlier suggestions. The function of the first coded peptides is not known, nor is the precision of coding required to enable this function. The degree of coding chemo- and stereoselectivity that we have discovered is not particularly high, but it might have resulted in the loosely coded synthesis of short peptides composed predominantly of L-amino acids. It would also have been something that nascent biology could have built on. Thus, ribozymes could have evolved to enhance this intrinsic chemical coding. These ribozymes could then have assisted the aminoacyl-transfer from progressively shorter oligonucleotide donors, ultimately resulting in the transfer from aminoacyl-adenylates. The chemistry, involving direct transfer to the 2′,3′-diol from a mixed anhydride of amino acid and the 5′-phosphate, resembles the second step of the chemistry catalyzed by aminoacyl-tRNA synthetases in extant biology. Concomitant improvements in chemo- and stereoselectivity along with assignment to codons and development of translation could then lead to the synthesis of peptides sufficiently well encoded that they could augment the function of the ribozymes and, ultimately replace them.

Our original choice of stem-overhang was based on a model for the origin of tRNA by direct gene duplication (*31, 36*). The ligation junction required by this model is situated in the anticodon loop and we postulated that an overhang sequence adopting a folded-back conformation would be most prone to such a ligation. Symmetry then dictated that what would have been destined to become the acceptor overhang would also have adopted a folded-back conformation (*37*). Based on this model, we then used both canonical extant acceptor stem-overhang sequence preferences and the corresponding anticodon loop sequence preferences in the design of the putative ancestral overhang. This resulted in a five-base overhang which is shorter than the extant anticodon loop (*38, 39*), but longer than the extant acceptor overhang. We then showed experimentally that the folded-back overhang indeed allows both aminoacyl-transfer from the 5′-phosphate to the 2′,3′-diol and loop-closing ligation (*40*). The pentanucleotide overhang and often frayed acceptor stem terminus that allow coded aminoacylation by purely chemical means would not be expected to remain unchanged as ribozyme- and then enzyme-catalyzed aminoacylation emerged. Indeed, predominant adoption of a G1:C72 base pair, replacement of the U responsible for the U-turn motif and truncation of the overhang would enable a more rigid acceptor stem-overhang which could project into a catalyst active site. So, extant tRNA acceptor stem overhang structures might only bear passing resemblance to ancestral structures. However, the coding by the terminal trinucleotide of the tRNA mimic provides a plausible explanation as to why modern tRNA identity determinants now cluster at the acceptor stem as well as at the anticodon.

The earliest aminoacyl-tRNA synthetase enzymes likely could not span the long distance (∼75Å in extant tRNAs) from the site of acceptor stem aminoacylation to the anticodon, but could easily recognize RNA in the vicinity of the aminoacylation site. If the enzymes originally evolved to build on the intrinsic chemical coding by the acceptor stem, they would be expected to use the trinucleotide or elements thereof in the process of cognate tRNA recognition. The position of these identity determinants within the tRNA molecule would be expected to be retained even if their sequence identity was changed over time.

## Acknowledgments

The authors thank JDS group members for fruitful discussions.

## Funding

Medical Research Council MC_UP_A024_1009 (JDS) Simons Foundation 290362 (JDS)

German Research Council CRC 235 – P11 (AJ)

## Author contributions

AJ conceived the selection protocol which was experimentally implemented by CS. DJW oversaw the modelling which was carried out by KR. JDS conceived the overall project and oversaw the mixed anhdride chemistry carried out by MS and the phosphoramidate chemistry carried out by ZL. SJR assisted with chemistry and KCL assisted with the sequencing protocol. The manuscript was 10 written by JDS, DJW and MS with input from the other authors. The SI was assembled by MS, CS, KR and ZL with input from the other authors.

## Competing interests

Authors declare that they have no competing interests.

## Data and materials availability

The data that support the findings of this study are available within the paper and its Supplementary Information. All raw sequencing 15 data and code for data cleaning and analysis associated with the current submission is available in a zenodo repository at 10.5281/zenodo.7515305. All simulation data is available at 10.5281/zenodo.7371596.

## Supplementary Materials

Materials and Methods Figs. S1 to S15

Tables S1 to S25

References (*41-52*)

## Notes

https://zenodo.org/record/7371596#.ZA9B_ex_pdo

https://zenodo.org/record/7515305#.ZA9Bxex_pdo

### Competing Interest Statement

The authors have declared no competing interest.

## References and Notes

1. M. A. Rubio Gomez, M. Ibba, Aminoacyl-tRNA synthetases. RNA 26, 910–936 (2020).

2. F. H. Crick, Codon--anticodon pairing: the wobble hypothesis. J Mol Biol 19, 548–555 (1966).

3. G. Gamow, Possible Relation between Deoxyribonucleic Acid and Protein Structures. Nature 173, 318 5 (1954).

4. S. R. Pelc, Correlation between coding-triplets and amino-acids. Nature 207, 597–599 (1965).

5. S. R. Pelc, M. G. Welton, Stereochemical relationship between coding triplets and amino-acids. Nature 209, 868–870 (1966).

6. M. Di Giulio, Arguments against the stereochemical theory of the origin of the genetic code. Biosystems 10 221, 104750 (2022).

7. J. J. Hopfield, Origin of the genetic code: a testable hypothesis based on tRNA structure, sequence, and kinetic proofreading. Proceedings of the National Academy of Sciences 75, 4334–4338 (1978).

8. M. Illangasekare, M. Yarus, Specific, rapid synthesis of Phe-RNA by RNA. Proc Natl Acad Sci U S A 96, 5470–5475 (1999).

9. N. V. Chumachenko, Y. Novikov, M. Yarus, Rapid and simple ribozymic aminoacylation using three conserved nucleotides. J Am Chem Soc 131, 5257–5263 (2009).

10. D. B. Johnson, L. Wang, Imprints of the genetic code in the ribosome. Proc Natl Acad Sci U S A 107, 8298–8303 (2010).

11. R. M. Turk, M. Illangasekare, M. Yarus, Catalyzed and spontaneous reactions on ribozyme ribose. J Am 20 Chem Soc 133, 6044–6050 (2011).

12. B. Jash, P. Tremmel, D. Jovanovic, C. Richert, Single nucleotide translation without ribosomes. Nat Chem 13, 751–757 (2021).

13. F. H. Crick, The origin of the genetic code. J Mol Biol 38, 367–379 (1968).

14. R. D. Knight, L. F. Landweber, The early evolution of the genetic code. Cell 101, 569–572 (2000).

15. M. E. Saks, J. R. Sampson, J. N. Abelson, The transfer RNA identity problem: a search for rules. Science 263, 191–197 (1994).

16. R. Giege, M. Sissler, C. Florentz, Universal rules and idiosyncratic features in tRNA identity. Nucleic Acids Res 26, 5017–5035 (1998).

17. R. Giegé, G. Eriani, The tRNA identity landscape for aminoacylation and beyond. Nucleic Acids Res 51, 30 1528–1570 (2023).

18. Y. M. Hou, P. Schimmel, A simple structural feature is a major determinant of the identity of a transfer RNA. Nature 333, 140–145 (1988).

19. W. H. McClain, K. Foss, Changing the identity of a tRNA by introducing a G-U wobble pair near the 3’ acceptor end. Science 240, 793–796 (1988).

20. C. Francklyn, P. Schimmel, Aminoacylation of RNA minihelices with alanine. Nature 337, 478–481 (1989).

21. C. de Duve, Transfer RNAs: the second genetic code. Nature 333, 117–118 (1988).

22. N. Maizels, A. M. Weiner, Phylogeny from function: evidence from the molecular fossil record that tRNA originated in replication, not translation. Proc Natl Acad Sci U S A 91, 6729–6734 (1994).

23. P. Schimmel, L. Ribas de Pouplana, Transfer RNA: from minihelix to genetic code. Cell 81, 983–986 (1995).

24. A. S. Petrov et al., History of the ribosome and the origin of translation. Proc Natl Acad Sci U S A 112, 15396–15401 (2015).

25. M. Yarus, The Genetic Code and RNA-Amino Acid Affinities. Life 7, (2017).

26. B. H. Patel, C. Percivalle, D. J. Ritson, C. D. Duffy, J. D. Sutherland, Common origins of RNA, protein and lipid precursors in a cyanosulfidic protometabolism. Nat Chem 7, 301–307 (2015).

27. Z. Liu et al., Harnessing chemical energy for the activation and joining of prebiotic building blocks. Nat Chem 12, 1023–1028 (2020).

28. B. J. Weimann, R. Lohrmann, L. E. Orgel, H. Schneider-Bernloehr, J. E. Sulston, Template-directed 50 synthesis with adenosine-5’-phosphorimidazolide. Science 161, 387 (1968).

29. A. L. Weber, L. E. Orgel, Amino acid activation with adenosine 5’-phosphorimidazolide. J Mol Evol 11, 9–16 (1978).

30. J. L. Shim, R. Lohrmann, L. E. Orgel, Poly(U)-directed transamidation between adenosine 5’-phosphorimidazolide and 5’-phosphoadenosine 2’(3’)-glycine ester. J Am Chem Soc 96, 5283–5284 (1974).

31. L. F. Wu, M. Su, Z. Liu, S. J. Bjork, J. D. Sutherland, Interstrand Aminoacyl Transfer in a tRNA Acceptor Stem-Overhang Mimic. J Am Chem Soc 143, 11836–11842 (2021).

32. S. J. Roberts, Z. Liu, J. D. Sutherland, Potentially Prebiotic Synthesis of Aminoacyl-RNA via a Bridging Phosphoramidate-Ester Intermediate. J Am Chem Soc 144, 4254–4259 (2022).

33. M. Arutaki et al., G:U-Independent RNA Minihelix Aminoacylation by Nanoarchaeum equitans Alanyl-tRNA Synthetase: An Insight into the Evolution of Aminoacyl-tRNA Synthetases. J Mol Evol 88, 501–509 (2020).

34. S. L. Miller, Production of Some Organic Compounds under Possible Primitive Earth Conditions1. J Am Chem Soc 77, 2351–2361 (1955).

35. A. P. Johnson et al., The Miller volcanic spark discharge experiment. Science 322, 404 (2008).

36. M. Di Giulio, On the origin of the transfer RNA molecule. J Theor Biol 159, 199–214 (1992).

37. E. V. Puglisi, J. D. Puglisi, J. R. Williamson, U. L. RajBhandary, NMR analysis of tRNA acceptor stem microhelices: discriminator base change affects tRNA conformation at the 3’ end. Proc Natl Acad Sci U S A 91, 11467–11471 (1994).

38. S. Limmer, H. P. Hofmann, G. Ott, M. Sprinzl, The 3’-terminal end (NCCA) of tRNA determines the structure and stability of the aminoacyl acceptor stem. Proc Natl Acad Sci U S A 90, 6199–6202 (1993).

39. A. Czech, Deep sequencing of tRNA’s 3’-termini sheds light on CCA-tail integrity and maturation. RNA 26, 199–208 (2020).

40. L. F. Wu et al., Template-Free Assembly of Functional RNAs by Loop-Closing Ligation. J Am Chem Soc 144, 13920–13927 (2022).

41. J. A. Joseph, K. Roder, D. Chakraborty, R. G. Mantell, D. J. Wales, Exploring biomolecular energy landscapes. Chem Commun (Camb) 53, 6974–6988 (2017).

42. K. Röder, J. A. Joseph, B. E. Husic, D. J. Wales, Energy Landscapes for Proteins: From Single Funnels to Multifunctional Systems. Adv Theory Simul 2, (2019).

43. A. Perez et al., Refinement of the AMBER force field for nucleic acids: improving the description of alpha/gamma conformers. Biophys J 92, 3817–3829 (2007).

44. M. Zgarbova et al., Refinement of the Cornell et al. Nucleic Acids Force Field Based on ReferencQuantum Chemical Calculations of Glycosidic Torsion Profiles. J Chem Theory Comput 7, 2886–290 (2011).

45. C. Tian et al., ff19SB: Amino-Acid-Specific Protein Backbone Parameters Trained against Quantum Mechanics Energy Surfaces in Solution. J Chem Theory Comput 16, 528–552 (2020).

46. G. W. T. M. J. Frisch, H. B. Schlegel, G. E. Scuseria, M. A. Robb, J. R. Cheeseman, J. A. Montgomery, Jr., T. Vreven, K. N. Kudin, J. C. Burant, J. M. Millam, S. S. Iyengar, J. Tomasi, V. Barone, B. Mennucci, M. Cossi, G. Scalmani, N. Rega, G. A. Petersson, H. Nakatsuji, M. Hada, M. Ehara, K. Toyota, R. Fukuda, J. Hasegawa, M. Ishida, T. Nakajima, Y. Honda, O. Kitao, H. Nakai, M. Klene, X. Li, J. E. Knox, H. P. Hratchian, J. B. Cross, V. Bakken, C. Adamo, J. Jaramillo, R. Gomperts, R. E. Stratmann, O. Yazyev, A. J. Austin, R. Cammi, C. Pomelli, J. W. Ochterski, P. Y. Ayala, K. Morokuma, G. A. Voth, P. Salvador, J. J. Dannenberg, V. G. Zakrzewski, S. Dapprich, A. D. Daniels, M. C. Strain, O. Farkas, D. K. Malick, A. D. Rabuck, K. Raghavachari, J. B. Foresman, J. V. Ortiz, Q. Cui, A. G. Baboul, S. Clifford, J. Cioslowski, B. B. Stefanov, G. Liu, A. Liashenko, P. Piskorz, I. Komaromi, R. L. Martin, D. J. Fox, T. Keith, M. A. Al-Laham, C. Y. Peng, A. Nanayakkara, M. Challacombe, P. M. W. Gill, B. Johnson, W. Chen, M. W. Wong, C. Gonzalez, and J. A. Pople,. (Gaussian, Inc., 2004).

47. C. I. Bayly, P. Cieplak, W. Cornell, P. A. Kollman, A well-behaved electrostatic potential based method using charge restraints for deriving atomic charges: the RESP model. J Phys Chem A 97, 10269–10280 (1993).

48. M. Popenda et al., Automated 3D structure composition for large RNAs. Nucleic Acids Res 40, e11 (2012).

49. D. J. Wales, J. P. K. Doye, Global Optimization by Basin-Hopping and the Lowest Energy Structures of Lennard-Jones Clusters Containing up to 110 Atoms. J Phys Chem A 101, 5111–5116 (1997).

50. O. M. Becker, M. Karplus, The topology of multidimensional potential energy surfaces: Theory and application to peptide structure and kinetics. J Chem Phys 106, 1495–1517 (1997).

51. D. J. Wales, M. A. Miller, T. R. Walsh, Archetypal energy landscapes. Nature 394, 758–760 (1998).

52. E. F. Pettersen et al., UCSF Chimera--a visualization system for exploratory research and analysis. J Comput Chem 25, 1605–1612 (2004).

