## Supplementary Materials for "Triplet-encoded prebiotic RNA aminoacylation"

**The PDF file includes:**

Materials and Methods

Figs. S1 to S15

Tables S1 to S25

References 41-52

**Materials and Methods**

Reagents and solvents were obtained from *Acros Organics*, *Santa Cruz Biotechnology*, *Sigma-Aldrich*, *Synthon Chemicals GmbH & Co. KG* and *VWR International*, and were used without further purification. For solid-phase RNA synthesis, primer Support 5G with a loading of 300 μmol/g was purchased from *GE Healthcare*. Phosphoramidites for RNA synthesis were purchased from *Sigma-Aldrich* or *Link Technologies*. RNA oligomers in Table S1 and **Lib-S3-D** were synthesized using an ÄKTA Oligopilot plus 10 (*GE Healthcare*) on a 20 to 50 μmol scale. All other RNA oligomers in Table S3 were purchased from *Sigma-Aldrich* with HPLC purification. The purity of purchased oligomers were not checked before used.

A *Mettler Toledo* SevenEasy pH Meter S20 combined with a *ThermoFisher Scientific* Orion 8103BN Ross semi-micro pH electrode was used to measure pH during adjustment to the desired value. ^1^H-, ^13^C-, and ^31^P-nuclear magnetic resonance (NMR) spectra were acquired using a *Bruker* Ultrashield 400 Plus or *Bruker* Ascend 400 operating at 400.13, 100.62, and 161.97 MHz, respectively. The notations s, t, and br represent the multiplicities of singlet, triplet, and broad signal, respectively. Chemical shifts (*δ*) are shown in ppm. Chemical shifts for P-NMR are referenced to 85% phosphoric acid, which is assigned the chemical shift of 0. Mass spectra were acquired on an *Agilent* 1200 LC-MS system equipped with an electrospray ionization (ESI) source and a 6130 quadrupole spectrometer. LC solvents: A, 0.2 % formic acid in H_2_O – B, and 0.2 % formic acid in acetonitrile. Analytical High-Pressure Liquid Chromatography (HPLC) was run on Dionex Ultimate 3000 (*Thermo Scientific*) using a *Waters* A*tlantis* T3, 3 μm, 4.6 × 150 mm column. LC solvents: A, 20 mM triethylammonium acetate aq., pH = 7.0, and B, acetonitrile. Preparative HPLC were run on *Varian* PrepStar using a *Waters* Altantis T3 Prep OBD, 5 μm, 15 × 250 mm column. LC solvents: A, 100 mM triethylammonium acetate aq., pH = 7.0, and B, acetonitrile. Oligonucleotide concentrations were determined by UV absorbance at 260 nm using a *NanoDrop* ND-1000 spectrophotometer.

Chemical synthesis of RNA oligomers

After automated synthesis, RNAs were first cleaved from the solid support by treating with 3 mL of a 1:1 mixture of NH_3_ aqueous solution (28% wt) and CH_3_NH_2_ ethanol solution (33% wt) at 55°C for 80 minutes in a tube with a sealed cap. The solid was removed by filtration and washed with 50 % EtOH/H_2_O. The solutions were combined and evaporated to dryness under reduced pressure. Silyl protecting groups were removed by treating the residues with 2 mL of a 1:1 mixture of triethylamine trihydrofluoride and DMSO at 65°C for 150 minutes in a tube with a sealed cap. After brief cooling at -32°C, 40 mL of cold 50 mM NaClO_4_ in acetone was added to the solution to precipitate the oligoribonucleotides. The resulting mixture was centrifuged and the recovered oligoribonucleotides purified by preparative HPLC followed by lyophilization. The purified RNA was redissolved in 2 mL of water and passed through a *Waters* Sep-Pak C18 Cartridge with 10 g sorbent.The cartridge was pre-washed with 50 mL of acetonitrile then 50 mL of water before sample loading, washed with 100 mL of H_2_O and 50 mL of 20 % aqueous acetonitrile. Eluates were checked for RNA content using a NanoDrop spectrophotometer, checked for purity using analytical HPLC, combined and lyophilized. The resulting white powder was stored at -32°C for future use.

Mixed anhydride chemistry

**Chemical synthesis of protected amino acids**

**Synthesis of *N*-4-monomethoxytritylvaline (MMTr-Val-OH), *N*-tritylleucine (Trt-Leu-OH), and *N*-tritylproline (Trt-Pro-OH)**

Trimethylsilyl chloride (1.27 mL, 10 mmol, 1 eq.) was added to a magnetically stirred suspension of the amino acid (Val, Leu, or Pro, 10 mmol, 1 eq.) in chloroform/acetonitrile (18 mL, v/v 5/1) at room temperature. The reaction mixture was heated under reflux for 2 h and then allowed to cool to room temperature. A chloroform solution (10 mL) of triethylamine (2.79 mL, 20 mmol, 2 eq.) and 4-monomethoxytrityl chloride or trityl chloride (10 mmol, 1 eq.) was added to the mixture. The reaction was further stirred at room temperature overnight before the addition of methanol (2 mL, 50 mmol). The resulting mixture was evaporated to dryness and the residue was treated with a mixture of diethyl ether (50 mL) and a pre-cooled aqueous solution of citric acid (5%, 50 mL). The organic phase was separated and washed with 1 M NaOH solution (2 × 20 mL) and water (2 × 20 mL). The aqueous layers were combined and neutralized to pH = 7 with concentrated HCl on ice, then extracted using diethyl ether. The organic phase was dried over Na_2_SO_4_ and concentrated to give the desired product as a yellowish or white foam.

*N*-4-Monomethoxytritylvaline (**MMTr-Val-OH**), 45% yield.

**^1^H NMR** (400 MHz, *d_6_*-DMSO) *δ* = 7.40 (4H, m, MMTr), 7.25 (6H, m, MMTr), 7.17 (2H, m, MMTr), 6.82 (2H, d, *J* = 8.7 Hz, MMTr), 3.71 (3H, s, OC***H_3_***), 2.94 (1H, d, *J* = 3.9 Hz, NHC***H***COOH), 1.68 (1H, m, C***H***(CH_3_)_2_), 0.82 (6H, t, *J* = 7.5 Hz, CH(C***H_3_***)_2_).

**^13^C NMR** (100 MHz, *d_6_*-DMSO) *δ* = 175.0 (***C***OOH), 157.4 (C4'), 147.0, 146.7 (2×C1), 138.2 (C1'), 129.9 (2×C4), 128.4 (2×C3, 2×C5), 127.6 (2×C2, 2×C5), 126.1 (C2', C6'), 112.9 (C3', C5'), 70.2 (***C***Ph), 60.7 (NH***C***HCOOH), 54.9 (O***C***H_3_), 32.9 (***C***H(CH_3_)_2_), 19.9 (CH(***C***H_3_)_2_), 17.6 (CH(***C***H_3_)_2_).

**Mass**: C_25_H_26_NO_3_^–^ [M-H^+^]: calc. 388.2, found 388.1.

*N*-Tritylleucine (**Trt-Leu-OH**), 47% yield.

**^1^H NMR** (400 MHz, CDCl_3_) *δ* = 7.37 (6H, m, Trt), 7.15 (6H, m, Trt), 7.07 (3H, m, Trt), 3.20 (1H, t, *J* = 7.2 Hz, NHC***H***COOH), 1.61 (1H, m, (***C***H(CH_3_)_2_), 1.40 (2H, t, *J* = 7.1 Hz, CHC***H_2_***)， 0.79 (6H, dd, *J* = 6.5, 21.8 Hz, CH(***C***H_3_)_2_).

**^13^C NMR** (100 MHz, CDCl_3_) *δ* = 179.9 (***C***OOH), 145.9 (3×C1), 128.9 (3×C3, 3×C5), 127.8 (3×C2, 3×C6), 126.5 (3×C4), 71.5 (***C***Ph), 55.4 (NH***C***HCOOH), 45.1 (CH***C***H_2_CH), 24.7 (***C***H(CH_3_)_2_), 23.6 (CH(***C***H_3_)_2_), 22.0 (CH(***C***H_3_)_2_).

**Mass**: C_25_H_26_NO_2_^–^ [M-H^+^]: calc. 372.2, found 372.1.

*N*-Tritylproline (**Trt-Pro-OH**), 52% yield.

**^1^H NMR** (400 MHz, CDCl_3_) *δ* = 7.43 (6H, m, Trt), 7.23 (6H, m, Trt), 7.17 (3H, m, Trt), 4.04 (1H, dd, J = 2.5, 9.3 Hz, NC***H***COOH), 3.41 (1H, m, NC***H***_2_), 3.00 (1H, m, NC***H***_2_), 1.81 (1H, m, C***H***_2_CH), 1.51 (1H, m, C***H***_2_CH), 1.10 (1H, m, NCH_2_C***H***_2_), 0.84 (1H, m, NCH_2_C***H***_2_).

**^13^C NMR** (100 MHz, CDCl_3_) *δ* = 175.9 (***C***OOH), 142.9 (3×C1), 129.1 (3×C3, 3×C5), 128.2 (3×C2, 3×C6), 127.2 (3×C4), 78.8 (***C***Ph), 65.2 (N***C***HCOOH), 50.9 (N***C***H_2_), 30.5 (***C***H_2_CH), 24.2 (N***C***H_2_CH_2_).

**Mass**: C_24_H_22_NO_2_^–^ [M-H^+^]: calc. 356.2, found 356.1.

**Chemical synthesis of oligoribonucleotide:amino acid mixed anhydrides.** A mixture of protected amino acids, i.e. MMTr-Val-OH (50 eq.), Trt-Leu-OH (50 eq.), Trt-Pro-OH (20 eq.), 1-ethyl-3-(3-dimethylaminopropyl)carbodiimide hydrochloride (EDC, 50 eq.) and 5-mer donor oligoribonucleotide (1 eq., 4-8 mg) were dissolved in a mixture of pyridine (600 μL) and formamide (300 μL). The resulting solution was stirred in a 0°C cold room for 2 h (proline), or 24 h (valine, leucine). Then the solution was centrifuged, and the clarified solution was transferred to a pre-cooled acetone solution of NaClO_4_ (50 mM, 4.0 mL), and maintained at ‑20°C for 10 min. The product was collected as a pellet by centrifugation and was briefly dried under vacuum to remove the remaining acetone and pyridine. The white solid (ca. 4-8 mg) was dissolved in formic acid aq. (50%, 200 μL) and placed on ice for 3 min (valine, leucine, proline). Then, a pre-cooled acetone solution of NaClO_4_ (50 mM, 4.0 mL) was added to the solution. The product was again collected as a pellet by centrifugation, briefly dried under vacuum, and immediately stored dry at -70°C for future use. The oligoribo-nucleotide:amino acid mixed anhydride pellet is stable at -70°C for at least 2 months. The purity was checked using analytical HPLC. The yields for mixed anhdrides donor RNA were >85% for Gly and L-/D-Ala, ~75% for L-Pro, ~60% for L-Leu, and ~30% for L-Val. The mass of product was characterized using LCMS by dissolving a tiny amount of the mixed anhydride in 0.2% formic acid aq. The product was characterized using ^31^P NMR by dissolving 4 mg of the mixed anhydride in 0.2% formic acid aq. (D_2_O/H_2_O, 1/9). Once dissolved, the oligoribonucleotide mixed anhydrides should be stored in liquid nitrogen or at -70°C and used within one month. Ala and Gly mixed anhydrides were prepared as previously published.

**^31^P NMR** (162 MHz, D_2_O) *δ* = 0.2 (5'-***P***O_3_H), -1.2 ­– -0.5 (O***P***O_3_), -7.9 (5'-***P***O_3_CO).

**Aminoacyl-transfer from a mixed anhydride donor to the 3'-terminus of an acceptor overhang.** In a typical reaction, 5-mer oligoribonucleotide:amino acid mixed anhydride donor (6 mM in pH = 5.0, 50 mM TEAA buffer, 3 μL) was added to a solution (180 µL, 100 µM) of 10-mer oligoribonucleotide acceptor in HEPES buffer (50 mM, pH = 6.8), NaCl aq. (100 mM), MgCl_2_ aq. (5 mM). The HPLC autosampler was set to 10°C unless otherwise noted, and samples (8 μL) were injected for analysis every 20 min. Each reaction was monitored 14-20 times. HPLC was monitored at 260 nm, with a flow rate of 1 mL/min. LC solvents: A, 10 mM triethylammonium acetate aq. (2.77 mL triethylamine and 1.50 mL acetate acid in 2.0 L ddH_2_O), pH = 5.1, and B, acetonitrile. Column compartment temperature was set to 30°C. The samples were kept in the autosampler and analyzed again after 72 h to give a baseline.

**Calculation of the corrected yields.** Because of the incomplete coupling chemistry and/or background hydrolysis of the donor oligoribonucleotide:amino acid mixed anhydride, the yields of aminoacyl-transfer product directly determined by integration of HPLC peaks did not correspond to the yields that would have been determined had the mixed anhydride donor strand been pure. From the amount of the 5'-phosphorylated donor strand at the initial time point and the composite decay curve of the peak for the mixed anhydride thereof (by hydrolysis and transfer), it was possible to use nonlinear regression analysis to infer the relative amounts of these two materials at time zero. The yield of the aminoacyl-transfer product observed by HPLC could then be scaled to a corrected yield based on the calculated amount of mixed anhydride at time zero.

**HPLC quantification and kinetic regression analysis.** The peak areas of the mixed anhydride donor strand, hydrolyzed mixed anhydride, acceptor strand and diol ester product were determined at different time points. The half-life and the inferred percentage of mixed anhydride at time zero were calculated by nonlinear regression using a one-phase exponential decay, e.g., $ln\left( P_{mixed anhydride} \right)=kt+b$, where *t* refers to the time. Nonlinear regression using a two-phase exponential association decay, e.g., $Yield=A \left( e^{-k_{1}t}-e^{-k_{2}t} \right)$ was used to calculate *k*_transfer_ (*k*_2_) and *k*_hydrolysis_ (*k*_1_). The half-life of the diol ester was calculated by *t*_1/2_ = 0.69 / *k*_1_. The baseline, determined by the HPLC trace after at least three half-lives, was deducted from the peak areas of transfer products before the regression analysis. The time at which the concentration of the diol ester was maximum was calculated as $t_{max}= \frac{1}{(k_{2}-k_{1})ln(k_{2}/k_{1})}$. The corrected yield was the peak yield from the nonlinear regression, e.g., ${Yield}_{calc.}=A\left( e^{-k_{1}t_{peak}}-e^{-k_{2}t_{peak}} \right)$, over the percentage of mixed anhydride at time zero. Data were processed using GraphPad Prism.

**Preparation of RNA aminoacyl-transfer library.** Oligoribonucleotides **Lib-L5**, **Lib-S3**, and oligodeoxyribonucleotide blocker were ordered from *Integrated DNA Technology* and purified by HPLC. **Lib-L5** and **Lib-S3** were labelled with Cyanine 5 at the 5'-terminus. DNA blocker was modified with a C3 spacer, i.e., CH_2_CH_2_CH_2_OH, at the 3'-terminus. Mixed anhydrides of the donor strand **SM-1** or **Lib-S3-D** were freshly prepared as described above and stored in TEAA buffer (50 mM, pH = 5.0) in liquid nitrogen. **Lib-S3-D** mixed anhydrides were used without any purification (See Fig. S2), and were not characterized by LC-MS or ^31^P-NMR due to the limited amount of material. **Lib-L5** or **Lib-S3** (1 mM, 1 µL), blocker (1 mM, 1 µL), HEPES buffer (500 mM, pH = 6.8, 2 µL), NaCl aq. (1000 mM, 2 µL), MgCl_2_ aq. (50 mM, 2 µL) were mixed, heated to 95°C for 4 min then slowly cooled down to 10°C in 1 h. To the mixture, donor strand mixed anhydride **SM-1-Aa** or **Lib-S3-D-Aa** (1 mM, 2 µL) and ddH_2_O were added to the final volume of 20 µL. The transfer reactions were carried out at 10°C. Transfer reactions with L-Ala, D-Ala, Gly, and L-Pro were quenched with 0.4% HCl (1 µL) after 1 h. Transfer reaction with L-Leu was quenched after 2 h; reaction with L-Val was quenched after 3 h.

**Free diol oxidation.** To the resulting reaction mixture (21 µL), sodium acetate aq. (50 mM, pH = 4.5, 8 µL), sodium periodate (200 mM, 5 µL), and nuclease-free water (6 µL) were added. The mixture was incubated for 40 sec at room temperature to oxidize the free 2',3'-diol of the unreacted acceptor strands before quenching by addition of ethylene glycol (2 µL) followed by thorough mixing. After further incubation for 1 min at room temperature, the reaction was adjusted to pH = 12 with NaOH aq. (1 M). Hydrolysis of the 2'-/3'-acyl ester was allowed to proceed for 30 sec and the mixture was then adjusted back to pH = 7 with HCl (1 M). RNA-grade glycogen (20 µg/µL, 2 µL) and sodium acetate (3 M, pH = 5.2, 9 µL) were then added with mixing, followed by isopropanol (93 µL). The mixture was centrifuged at 16,000 g for 20 min at 4°C. The supernatant was removed and ethanol aq. (75%, 500 µL) was added. The mixture was further centrifuged at 16,000 g for 5 min at 4°C. The supernatant was removed and the pellet was lyophilized until dry.

**Ligation.** Oligodeoxyribonucleotides (phosphorylated at the 5'-terminus and modified with a C3 spacer, i.e. CH_2_CH_2_CH_2_OH at the 3'-terminus) were purchased from Integrated DNA Technologies. An ImpA/MgCl_2_ stock solution was prepared by dissolving adenosine-5’-phosphoimidazolide (ImpA, 7.1 mg) in an aqueous MgCl_2_ solution (100 mM, 90 µL). One adaptor at a time (1 mM, 60 µL) was mixed with the ImpA/MgCl_2_ solution (60 µL) and the resultant solution was incubated at 50°C for 1.5 h. ImpA/MgCl_2_ solution (30 µL) and water (30 µL) were added and incubation was continued at 50°C for another 1.5 h. Water (820 µL) was added to reach 1 mL final volume. The oligonucleotides were purified using two NAP-5 columns (2 × 500 µL in parallel) according to the manufacturer’s instructions. The typical yield for this reaction was 50%. In a subsequent reaction, the non-adenylated oligonucleotides were digested with Lambda Exonuclease (*NEB* M0262) according to the manufacturer’s instructions and using 5 µg as input. The adaptors were subsequently purified by phenol/chloroform/ isoamyl alcohol (PCI, 25/24/1) extraction and ethanol precipitation. Finally, the resulting material was dissolved in nuclease-free water (100 ng/µL) and analyzed via PAGE supplemented with N-acryloyl-3-aminophenylboronic acid (APB). 1 µg of each sample was mixed with T4 RNA Ligase Reaction Buffer (10×,2 µL), PEG8000 (50%, 4 µL), digested adaptor (1 µL, 100 ng), T4 RNA Ligase 2 truncated KQ (*NEB* M0373, 1 µL), and nuclease-free water (12 µL) were added and the resultant solution was incubated at room temperature for 2 h. 5’-Deadenylase (*NEB* M0331, 1 µL) was then added and incubation continued for 30 min at 30°C before addition of Lambda Exonuclease (*NEB* M0262, 1 µL) and further incubation for 30 min at 37°C. Finally, the reaction was purified via PCI-extraction and ethanol precipitation.

**Reverse transcription.** The pellet from the purified ligation reaction was dissolved in 11 µL nuclease‑free water and mixed with dNTP-mix (10 mM each dATP, dCTP, dGTP, dTTP, 1 µL) and RT‑primer (2 µM, 1 µL). The resultant solution was incubated at 65°C for 5 min and subsequently cooled down to 4°C. SuperScript IV RT-Buffer (5×, 4 µL), nuclease‑free water (1 µL), DTT (100 mM, 1 µL) and SuperScript IV reverse transcriptase (*ThermoFisher* 18090200, 1 µL) were added and the solution was then incubated at 55°C for 10 min and at 80°C for another 10 min before being cooled down to 4°C.

**PCR.** The whole reverse transcription reaction (20 µL) was used as the template for PCR. It was supplemented with nuclease-free water (13.5 µL), Q5 reaction buffer (5×,10 µL), forward and reverse primer (10 µM, 2.5 µL each), dNTP-mix (10 mM each, 1 µL) and Q5 High-Fidelity DNA Polymerase (*NEB* M0491, 0.5 µL). The reaction was heated to 98°C for 30 sec then subjected to four cycles of 98°C for 10 sec, 67°C for 15 sec, and 72°C for 10 sec. Final extension was carried out at 72°C for 45 sec. The reaction products were subjected to AMPure bead purification, analyzed using Bioanalyzer DNA High Sensitivity Chips and sequenced on an iSeq 100 all according to the manufacturer’s instructions.

Phosphoramidate chemistry

**Chemical synthesis of oligoribonucleotide:amino acid phosphoramidates.**

4-(Dimethylamino)pyridine (DMAP, 6.6 mg, 0.054 mmol) and RNA (**SM-1**, 5'-pAGCGA-3', 3 mg, 1.8 μmol, or **Lib-S3-D**, 5'-pNNNGAAUC-3', 3 mg) were dissolved in H_2_O/D_2_O (9:1 v/v, 0.5 mL) and the pH value was adjusted to 8.0 with hydrochloric acid (5 M). To the resulting solution, 1-ethyl-3-(3-dimethylaminopropyl)carbodiimide hydrochloride (EDC, 50 mg, 0.23 mmol) was added. The reaction mixture was kept at room temperature for 2 hours, and the reaction was followed by ^31^P-NMR spectroscopy. Afterwards, the solution was added dropwise to a cold solution of NaClO_4_ in acetone (50 mM, 10 mL) to precipitate the oligonucleotide-5'-DMAP phosphoramidate intermediate. The precipitate was separated by centrifugation and washed with diethyl ether (10 mL), then dried *in vacuo*. A solution of amino acid (50 mg/mL) was prepared and its pH was adjusted to 8.0 by the addition of NaOH, then 0.5 mL of this solution was added to the dried precipitate and the resultant solution was incubated at room temperature for 18 hours. The reaction was monitored by ^31^P-NMR spectroscopy or HPLC. The **Lib-S3-D** products were used without any purification (See Fig. S4). The **SM-1** phosphoramidate products were purified using analytical HPLC. LC solvents: A, 50 mM triethylammonium acetate, pH = 7.0 in water; B, acetonitrile. Column compartment temperature: 25°C. The product was stored as a solid or dissolved in basic pH solution at -32°C for future usage. Yields for two steps: 12% for Gly, 13% for L-/D-Ala, 40% for L-Pro, 17% for L-Leu, 21% for L-Val.

**^31^P NMR** (162 MHz, D_2_O) *δ* = 7.1 (5'-***P***O_3_N), 0.2 (5'-***P***O_3_H), -1.2 – -0.5 (O***P***O_3_),

**Preparation of RNA phosphoramidate-ester library.** A mixture (10 μL) containing the above synthesized RNA phosphoramidate (**SM-1** phosphoramidate or **Lib-S3-D** phosphoramidate 50 μM), aminoacyl acceptor RNA (**Lib-L5** or **Lib-S3**, 50 μM), DNA blocker (Blocker**,** 50 μM), NaCl (200 mM), MgCl_2_ (50 mM), EDC (50 mM) and imidazole (10 mM) in HEPES buffer (100 mM, pH = 7.0), was incubated at room temperature for 2 hours. Over time, 0.5 μL aliquots were taken and mixed with 2 μL of loading dye (9.5 M urea, 25 mM EDTA, pH = 8.0, blue bromophenol) before being analysed using polyacrylamide gel electrophoresis.

**Ligation.** The above pellet was dissolved in ddH_2_O. Cyanine3 labelled 36-mer ligator was purchased from *Integrated DNA Technology*. A mixture (20 µL) of the oxidized library, or the RNA library **Lib-L5** or **Lib-S3** as blank (final concentration 10 µM), ligator (final concentration 20 µM), ATP (10 mM, 2 µL), PEG 8000 (50%, 6 µL), T4 buffer (10×, 2 µL), and T4 RNA ligase 1 (*NEB* M0204, 10 U/µL, 1 µL) was incubated at 25°C for 2 h followed by inactivation at 65°C for 15 min. The mixture was cleaned up using *NEB* Monarch RNA Cleanup Kit (*NEB* T2030, 10 μg). A 6 µL eluate was obtained. The nucleic acid concentration was checked using a NanoDrop spectrophotometer and the composition was analyzed by 12% denaturing polyacrylamide gel electrophoresis.

**Reverse transcription.** The resulting solution (2 µL), reverse transcription primer (RT primer, 50 µM, 2 µL), dNTP (10 mM, 1µL) and nuclease-free H_2_O were mixed to a final volume of 10 µL. The mixture was heated at 65°C for 5 min and then placed on ice immediately. M-MuLV buffer (10×, 2 µL), M-MuLV reverse transcriptase (*NEB* M0253, 200 U/µL, 1 µL), RNase inhibitor (*NEB* M0314, 40 U/µL, 0.2 µL) and nuclease-free H_2_O were added to the mixture to a final volume of 20 µL. The cDNA synthesis reaction was incubated at 25°C for 5 min, then 37°C for 1 h, and inactivated at 80°C for 10 min. RNase H (*NEB* M0297, 5 U/µL, 0.2 µL) was added and the mixture was further incubated at 37°C for 20 min followed by inactivation at 80°C for 10 min. The mixture was cleaned up using a Monarch PCR & DNA Cleanup Kit (*NEB* T1030, 5 μg) following the oligonucleotides cleanup protocol. A 6 µL eluate was obtained and the nucleic acid concentration therein was checked using a NanoDrop spectrophotometer.

**PCR.** cDNA from above reverse transcription was prepared for Illumina sequencing by PCR with Fwd and Rev primers containing Illumina adapters plus indexes. The reverse transcription product (2 µL), dNTP (10 mM, 0.5 µL), Fwd and Rev primers (10 µM, 0.5 µL each), Taq buffer (10×, 2.5 µL), Taq polymerase (*NEB* M0273, 5 U/µL, 0.125 µL) and nuclease-free water were mixed to a final volume of 25 µL. PCR amplification was carried out as follows: pre-denaturation at 95℃ for 30 s, followed by denaturation at 95°C for 30 s, annealing at 55℃ for 30 s, and extension at 68℃ for 15 s, for a total of 15 cycles. The final extension was at 68℃ for 5 min. Exonuclease I (ExoI, *Thermo Scientific*, EN0581, 20 U/µL, 2 µL) was added directly after the completion of PCR. The resultant mixture was incubated at 37℃ for 20 min, the exonuclease inactivated at 85℃ for 15 min, and the products purified using a Monarch PCR & DNA Cleanup Kit (5 μg) using the PCR cleanup protocol. The volume of eluate obtained was 10 µL, a sample was analyzed by 6% denaturing polyacrylamide gel electrophoresis, and the remainder quantified using an *Invitrogen* Qubit 2.0 Fluorometer. Next Generation Sequencing was performed on an *Illumina* MiSeq system with MiSeq Reagent Kit v3 (150-cycle) according to the methods guide from the manufacturer.

**Data processing.** The raw FASTQ data was first filtered and quality-controlled using custom VBA scripts in Microsoft Excel. Frameshifts were corrected and reads with unidentified bases as well as PCR duplicates were excluded. Afterwards, the results were processed, analyzed and visualized using bowtie2, PEAR, and R. ‘ggpairs’ was used to calculate the Pearson correlation coefficient between the results of two amino acids. The correlation is expressed with a value between -1 to 1, where -1 shows negative correlation while 1 indicates positive correlation. P-value of permutation test for Pearson correlation coefficient is displayed as ‘***’ if the p-value is < 0.001, ‘**’ if the p-value is < 0.01; ‘*’ if the p-value is < 0.05; ‘.’ if the p-value is < 0.10; ‘’otherwise. Scores for the selectivity of L-Ala, Gly, L-Pro, L-Leu, L-Val and L-Leu/Val aminoacyl-ester formation were calculated as the ratios of the amino acid ranking (or sum of the rankings) to the sum of all the rest amino acids rankings for the same triplet sequence.

Melting temperature measurement using MicroCal VP-Capillary DSC

Differential scanning calorimetry (DSC) experiments were performed on a *General Electric* MicroCal Variable Pressure-capillary DSC. The donor strand (without mixed anhydride) and the acceptor strand (100 µM each) were dissolved in the same buffer (400 µL) as used in the transfer reactions, i.e., 50 mM HEPES pH 6.8, 100 mM NaCl, 5 mM MgCl_2_. The sample was initially maintained at 5°C then heated to 80°C at the rate of 2°C/min before being cooled down to 5°C at the same rate. The heating/cooling cycle was repeated three times. A blank buffer was used to calibrate the DSC curve.

Structure simulation

To obtain the characteristics of the structural ensembles adopted by the aminoacyl-tRNA overhang mimics, we used the computational energy landscape framework (*41, 42*). All simulation data is available in a zenodo repository (10.5281/zenodo.7371596).

The simulations used the AMBER OL3 force field for RNA (f99 with Barcelona α/γ backbone modifications and the χ correction for RNA (*43, 44*) in implicit solvent (igb = 2). The parameters for the amino acyl groups were derived in the follows. The amino acid equivalent N-terminal residue from the ff19SB force field (*45*) was used as template. We edited this residue manually to add a phosphate group with a methyl cap. The structure was then optimised with Gaussian (*46*) and charges fitted using RESP (*47*). The library file was then created by modifying the entries for the corresponding N-terminal amino acid form the AMBER library adding the missing information and updating the coordinates and charges. A single missing parameter, the CX-C-OS angle, was added as force modification file from GAFF. The libraries for the L-amino-acyl groups for Ala, Pro, Leu and Val, the D-amino-acyl group for Ala and the amino-acyl group for Gly are available in the online repository.

We used RNAcomposer (*48*) to obtain initial three-dimensional structures. As this tool requires a single chain, we added a UUUUU segment between the strands and enforced basepairing in the stem. The U loop was then removed, and the aaX group added. Basin-hopping global optimisation (*49*) was used to find low energy configurations, and the lowest 50 structures saved. We used short and long BH runs of 5000 and 25000 steps, respectively. The low-lying structures from both runs were used to seed our explorations of the energy landscape.

**Fig. S1.** Schematic representation of the library preparation workflow.

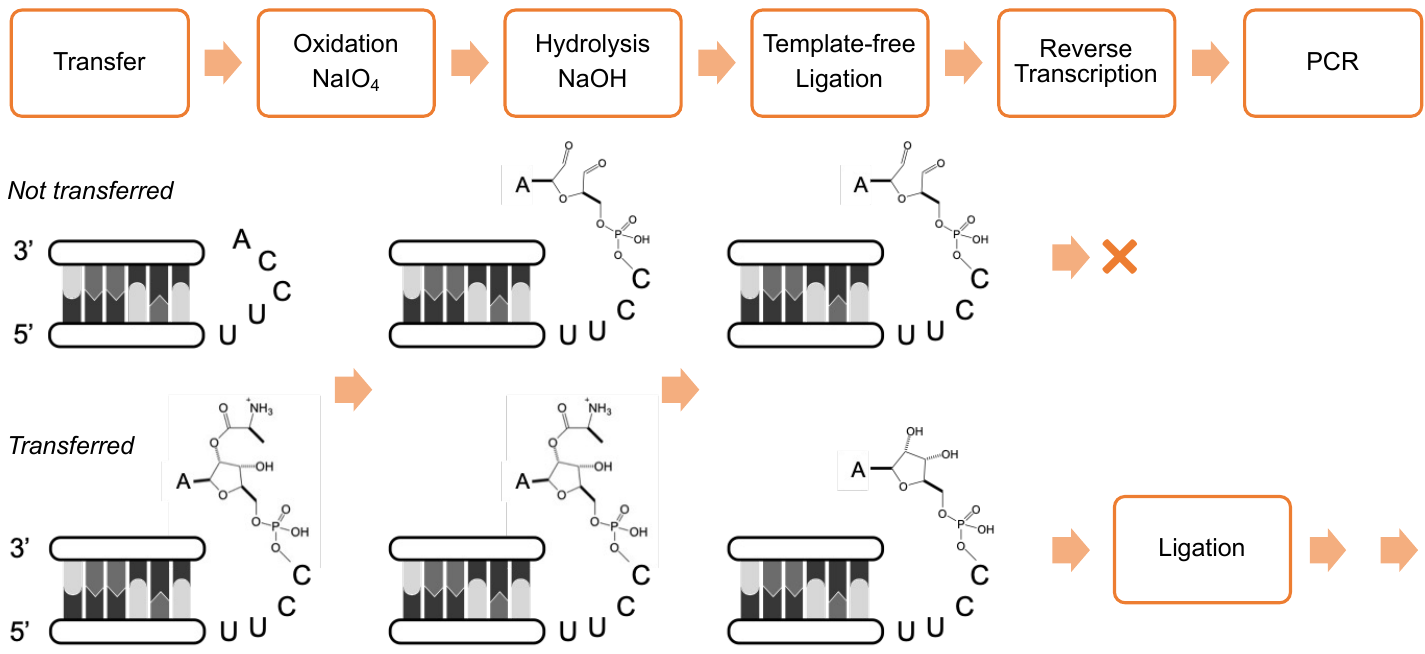

**Fig. S2. Consistent patterns in the overhang.** Plot of sequencing results for different pentanucleotides in the overhang (5′-UCGCUNNNNN) to show the correlation between amino acids: (**A**) mixed anhydride chemistry; (**B**) phosphoramidate chemistry. Sequence logos represent the top 10% of pentanucleotides from the quality-filtered read numbers. Pearson correlation coefficients are shown in the upper right corners.

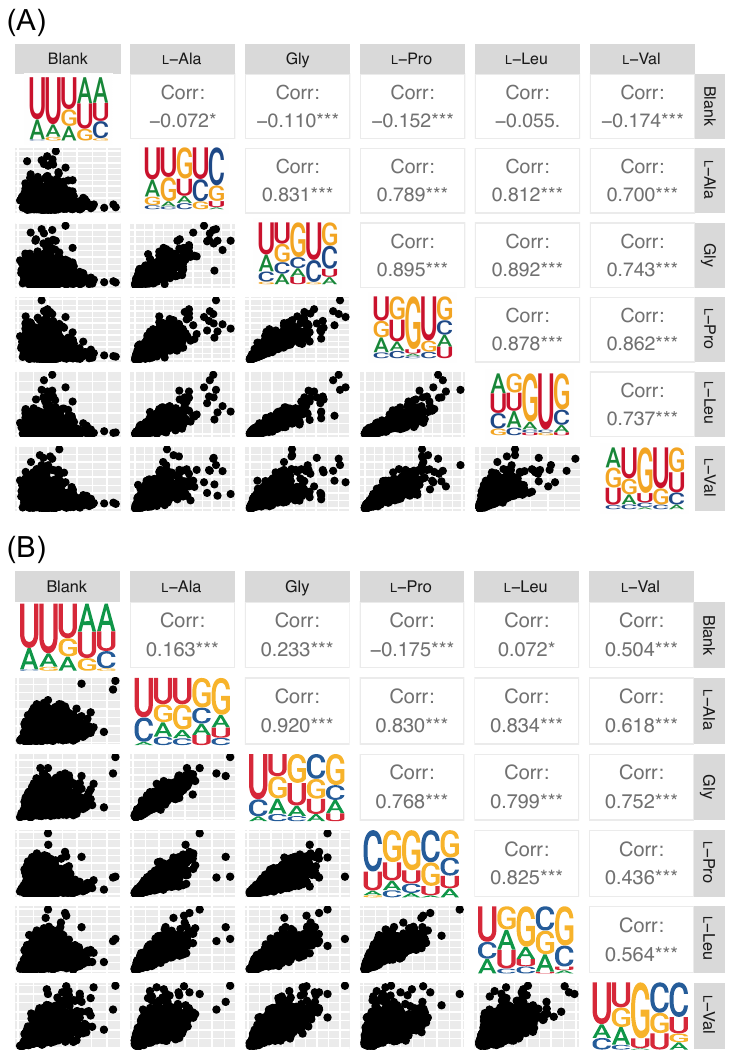

**Fig. S3.** Formation of RNA:amino acid mixed anhydrides with **Lib-S3-D.** HPLC conditions: Atlantis^TM^ T3, 5 μm, 4.6 x 250 mm column; flow rate 1 mL/min; LC solvents: A, 10 mM triethylammonium acetate, pH 5.1 in water and B, acetonitrile. Gradient: 0 min (7% B), to 1 min (7% B), to 15 min (20% B), to 15.5 min (95% B). Column compartment temperature is set at 25°C. Chromatograms, in descending order, show the HPLC profiles for the unmodified library and oligonucleotide mixed anhydrides.

**Fig. S4.** Sequencing results for acceptor strand with random trinucleotide in the stem (**Lib-S3**, and **Lib-S3-D** 5'-pNNNGAAUC) using mixed anhydride chemistry. Reads for the trinucleotide CUC omitted from the glycine data to enable visualization of other sequence preferences. Sequence logo represents top 10% trinucleotides from the raw reading numbers. Pearson correlation coefficients are shown in the upper right corner.

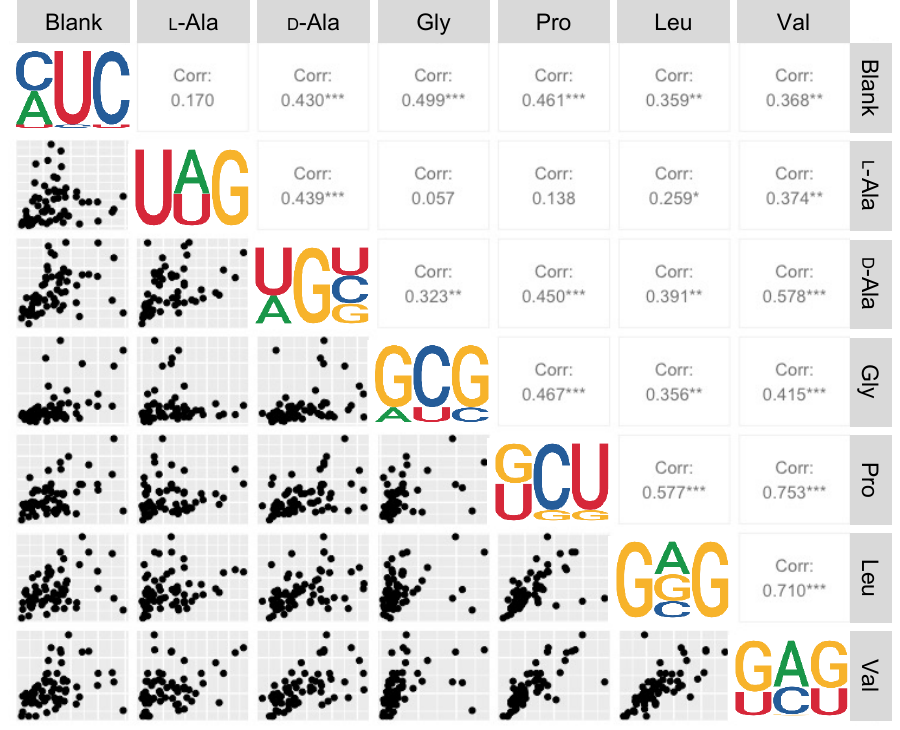

**Fig. S5.** Formation of RNA:amino acid phosphoramidate with **Lib-S3-D.** HPLC conditions: Atlantis^TM^ T3, 5 μm, 4.6 x 250 mm column; flow rate 1 mL/min; LC solvents: A, 20 mM triethylammonium acetate, pH 7 in water and B, acetonitrile. Gradient: 0 min (7% B), to 1 min (7% B), to 20 min (12% B), to 22 min (16% B) and 23 min (95% B). Column compartment temperature is set at 25°C. Chromatograms, in descending order, show the HPLC profiles for the unmodified library; the library after conversion to a DMAP phosphoramidate intermediate; products of reaction of the DMAP phosphoramidate intermediate with the amino acids indicated.

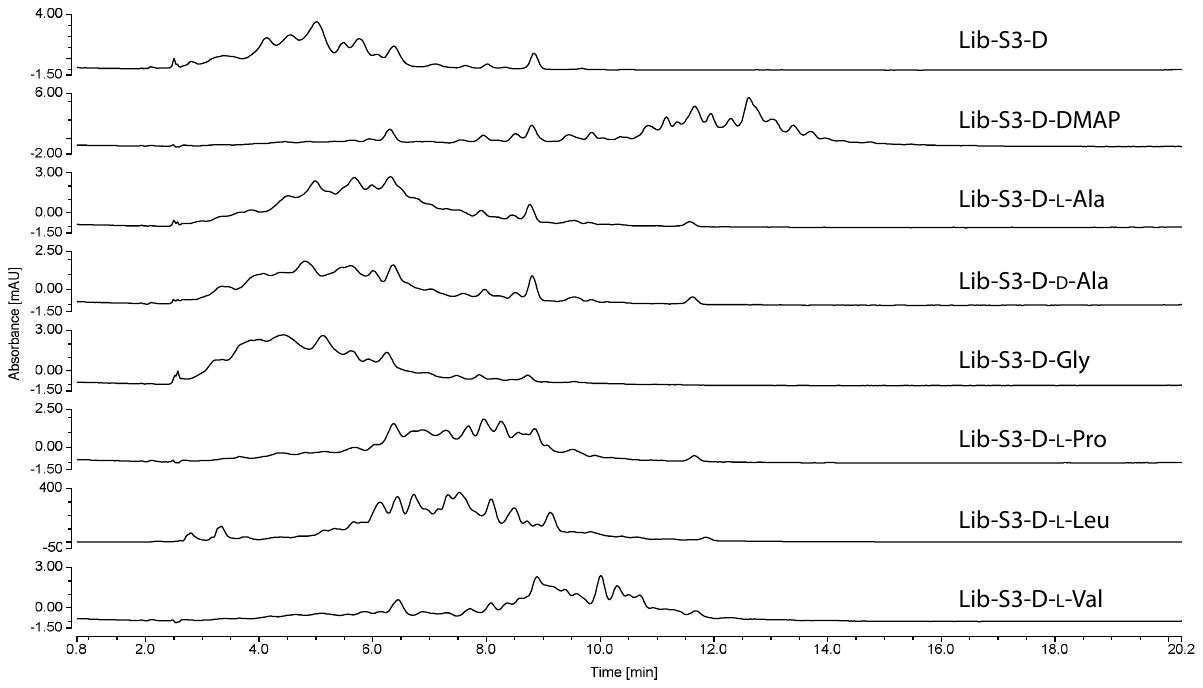

**Fig. S6. Consistent patterns in the first three nucleotides of the stem using phosphor-amidate chemistry.** Plot of sequencing results for different trinucleotides in the stem (5′-GAUUCNNNUUCCA) to show the correlation between amino acids. Sequence logos represent the top 10% of trinucleotides from the quality-filtered read numbers. Pearson correlation coefficients are shown in the upper right corner.

**Fig. S7.** DSC melting curves of 100 µM duplex **SM-11-Acc** with **SM-11** (black), **SM-16** (blue), and **SM-17** (orange). Curve in gray shows the blank using buffer, e.g. 50 mM HEPES, 100 mM NaCl, 5 mM MgCl_2_, pH = 6.8.

3’	GUC^p^

CGUC^p^

GCGUC^p^

5’ CGCAGUUCCA

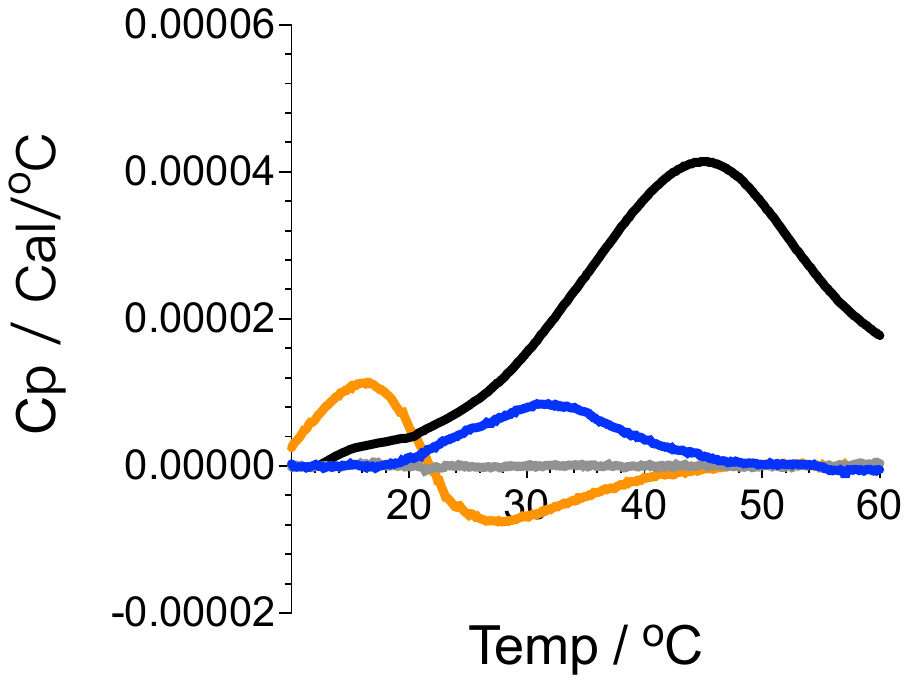

**Fig. S8.** Disconnectivity graphs (*50, 51*) for the free energy landscapes of L-alanyl- and L-leucyl- transfer using the sequences of donor and acceptor strands indicated. (**A**) **SM-18-Ala**; (**B**) **SM-4-Ala**; (**C**) **SM-6-Leu**; (**D**) **SM-19-Leu**. The order parameters are as follows. For the Ala sequences (**A** and **B**) the colour indicates the distances between the NH_3_^+^ group of the amino acid attached to the acceptor and the exocyclic N^6^ in the Hoogsteen edge of adenosine in the third base pair (**SM-18-Ala**, **A**) or the O^6^ in the guanine in the third base pair (**SM-4-Ala**, **B**). For the Leu sequences (**C** and **D**), it is the smallest distance between the bridging anhydride oxygen bnetween P and C in the phosphate linking Leu with the first nucleotide, and the Hoogsteen edge in the second base pair (O^6^ in guanine for **SM-6-Leu**, **C**, and exocyclic N^4^ in cytosine for **SM-19-Leu**, **D**). Green and blue indicate distances beyond 10 Å, while red highlights closer proximity. For both **SM-4-Ala** and **SM-19-Leu** (**B** and **D**), i.e. the sequences with lower observed yields, such contacts are formed in the lowest energy structures, but they are absent in **SM-18-Ala** and **SM-6-Leu** (**A** and **C**). The contacts make a close approach of the overhang to the amino acyl group geometrically challenging, and, as a result, we expect an activation barrier for the transfer, as these contacts need to be lost first. This effect likely lowers the yield, but the contacts are not strong enough to completely suppress the reaction.

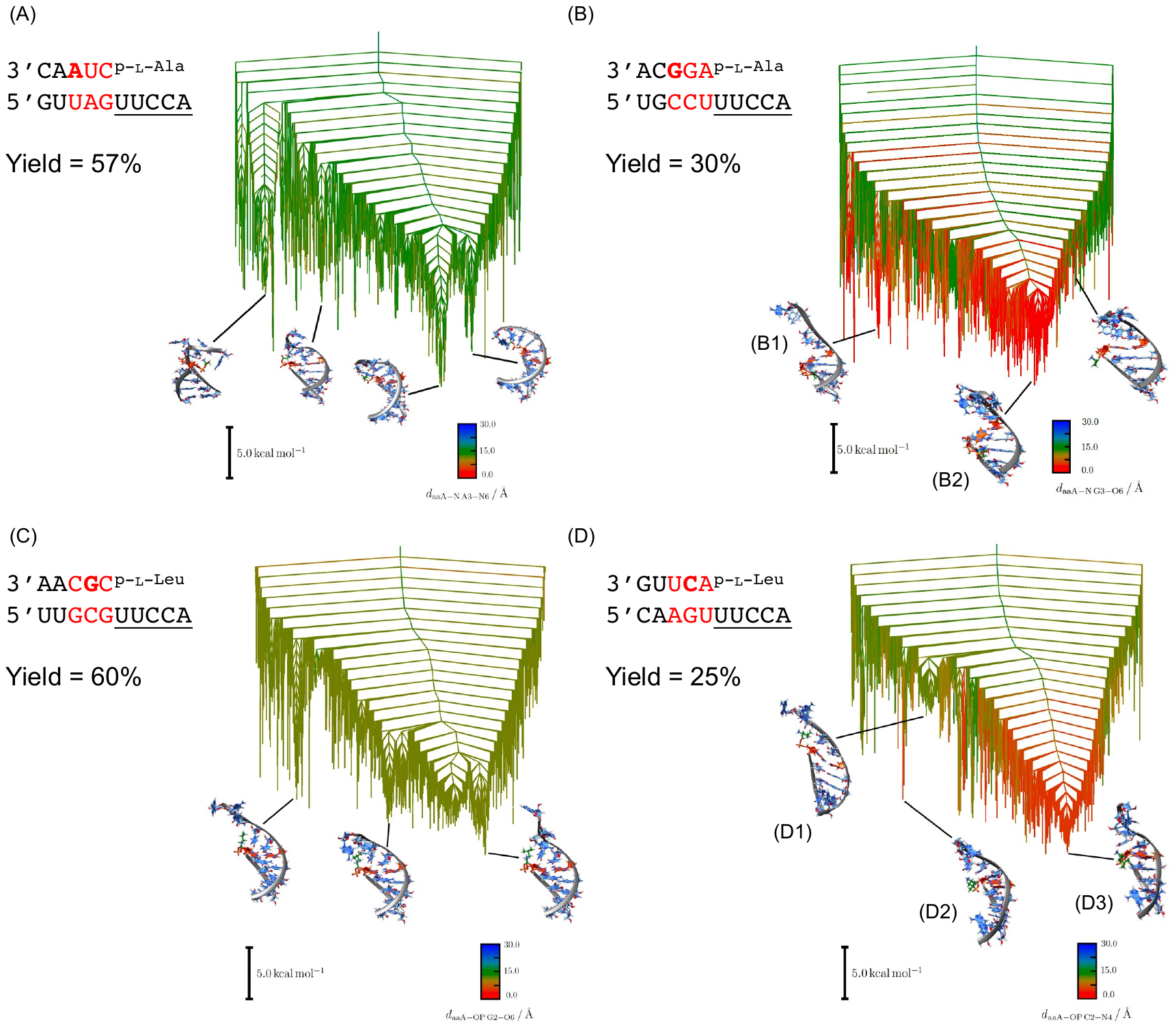

**Fig. S9.** Representative structures for L-Leu and L-Val selected from the energy landscape databases for two sequence combinations with different preferences. Kinetic curves shown in **Fig. 4A** (**A**-**C**) and **4B** (**D**-**F**), kinetic data presented in **Tables S18** (**A**-**C**) and **S19** (**D**-**F**). In **SM-11-Leu** (**A** and **B**) and **SM-11-Val** (**C**), the Leu and Val amino acyl groups form some interactions, but they are effectively free to move, and the transfer reaction should be possible. For **SM-13-Leu** (**D**) this picture is unchanged, and we therefore find no effect that would impede the transfer reaction. However, in **SM-13-Val**, i.e. the same sequence but with Val, we see significant changes. Stronger interactions are formed, which either result in interactions with and stacking of the overhang (**E**) or the AU base pair on top of the stem is lost, and strong interactions are formed (**F**). Both of these changes result in a barrier to the transfer explaining why the Val transfer has a significantly lower yield than the Leu transfer. Given the position of the side chain of Leu in the structures, and the strong charge interactions in RNA, wespeculate that the behaviour is based on the difference in hydrophobicity. The amino moiety is highlighted in green and the top base pair in orange. The images were created with Chimera (*52*).

**Fig. S10.** ^1^H- and ^13^C-NMR for *N*-4-monomethoxytritylvaline (contains triethylamine)

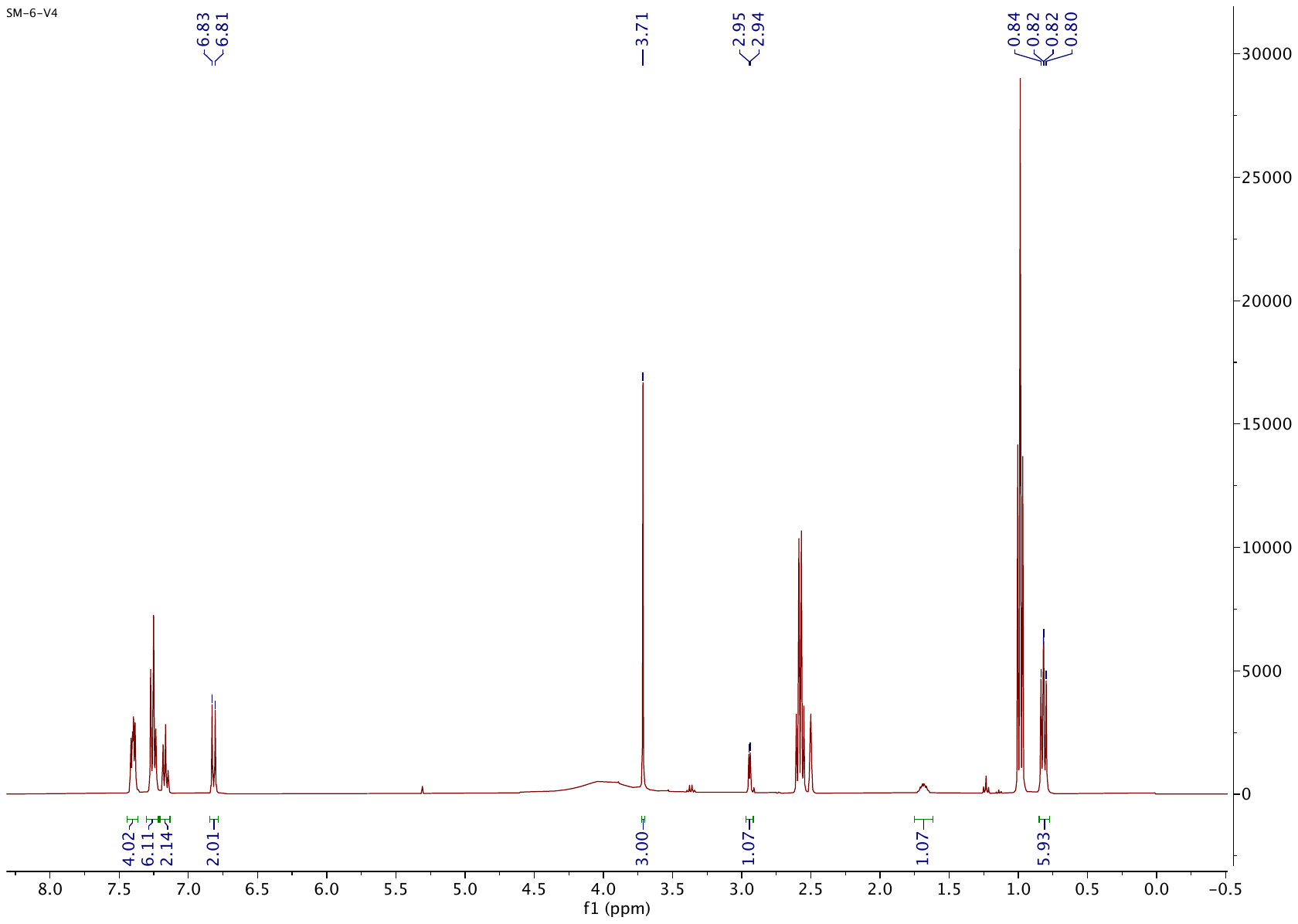

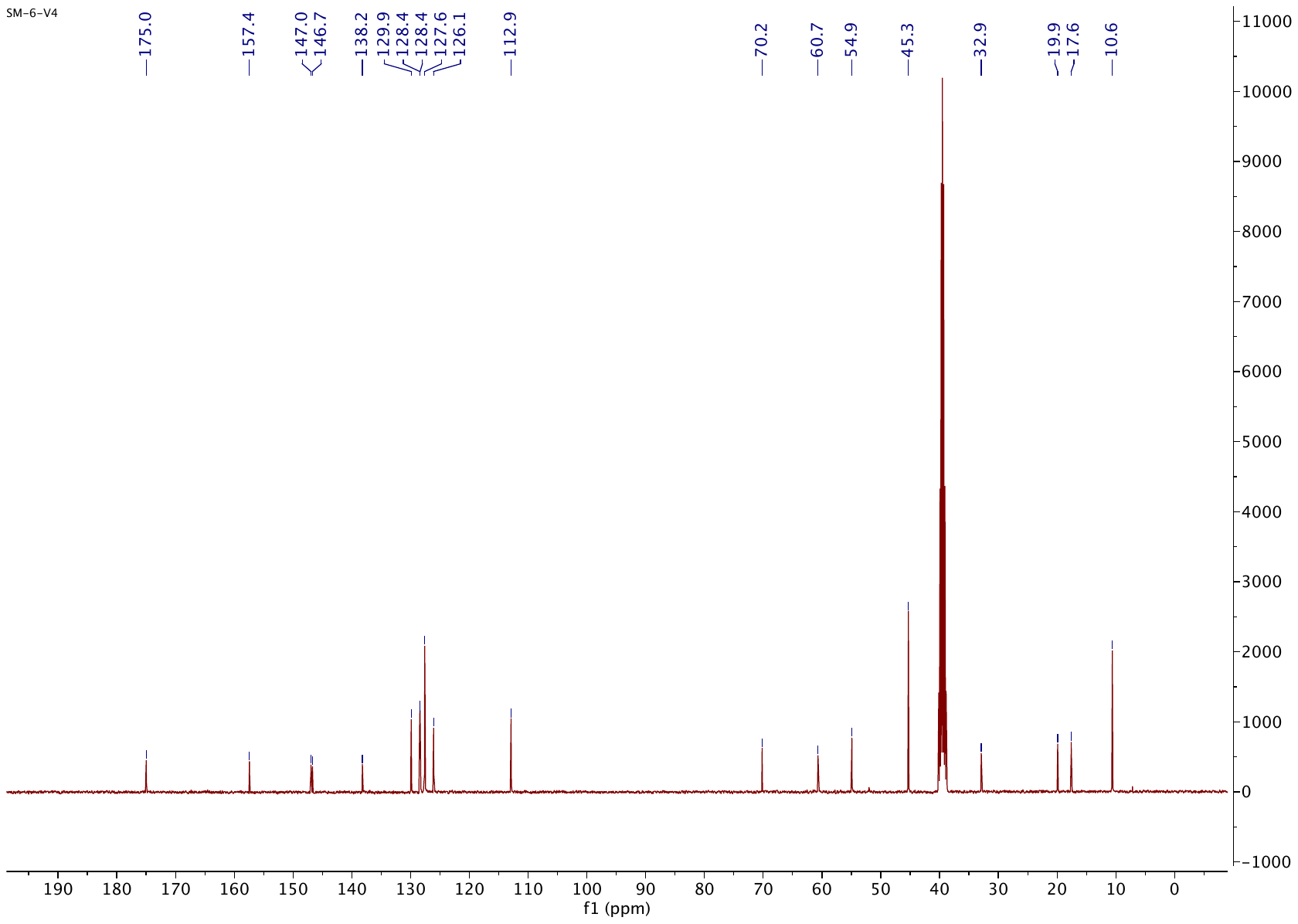

**Fig. S11.** ^1^H- and ^13^C-NMR for *N*-tritylleucine

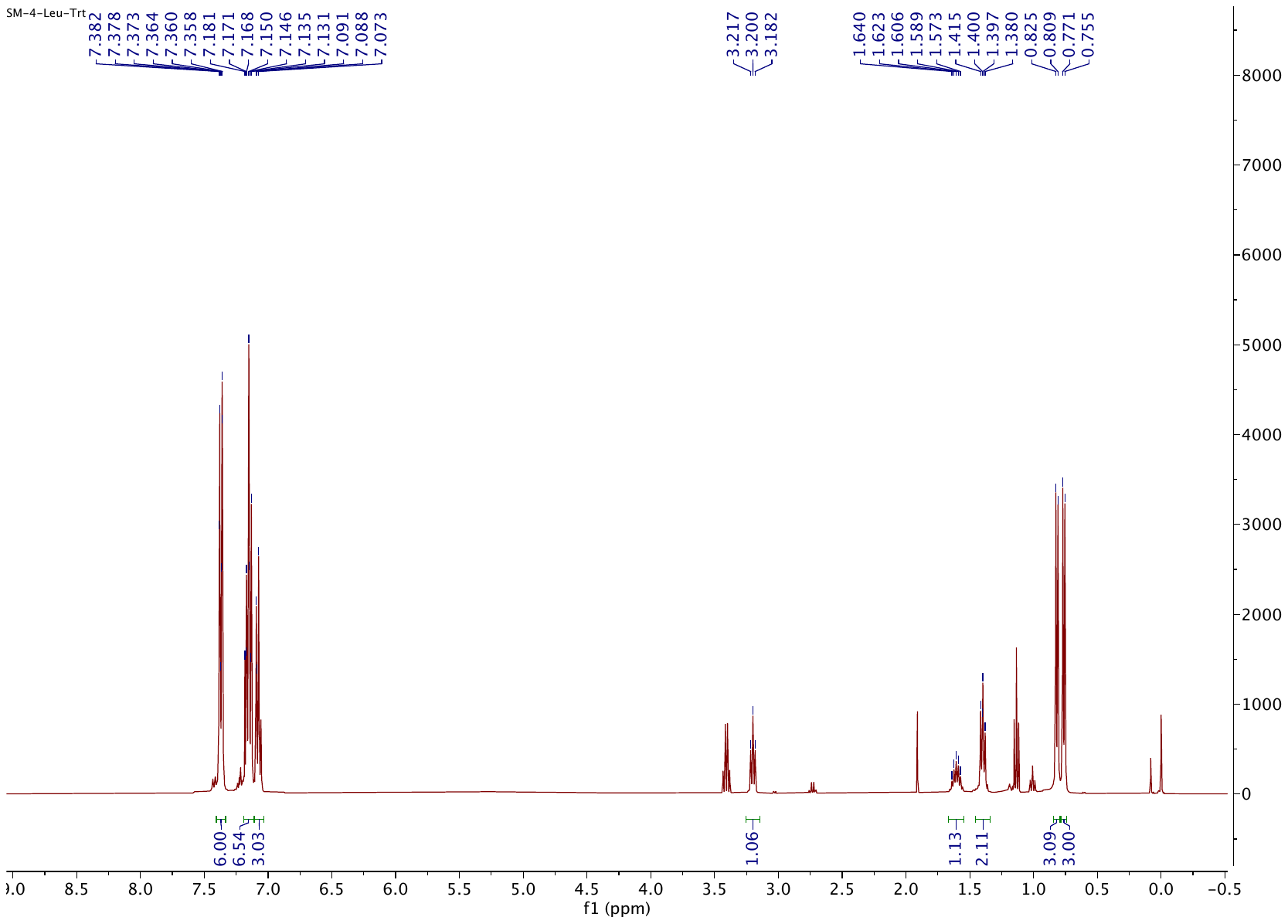

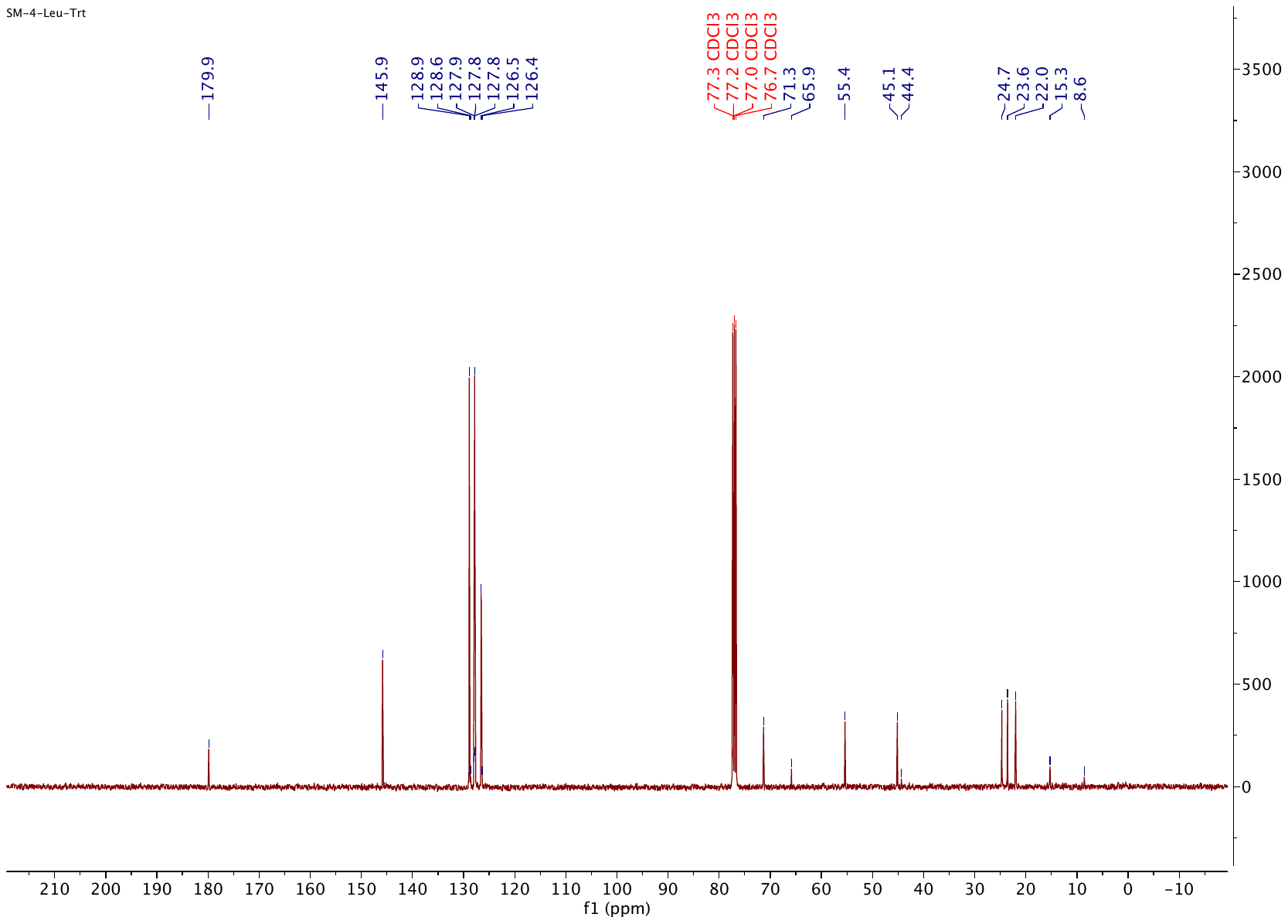

**Fig. S12.** ^1^H- and ^13^C-NMR for *N*-tritylproline

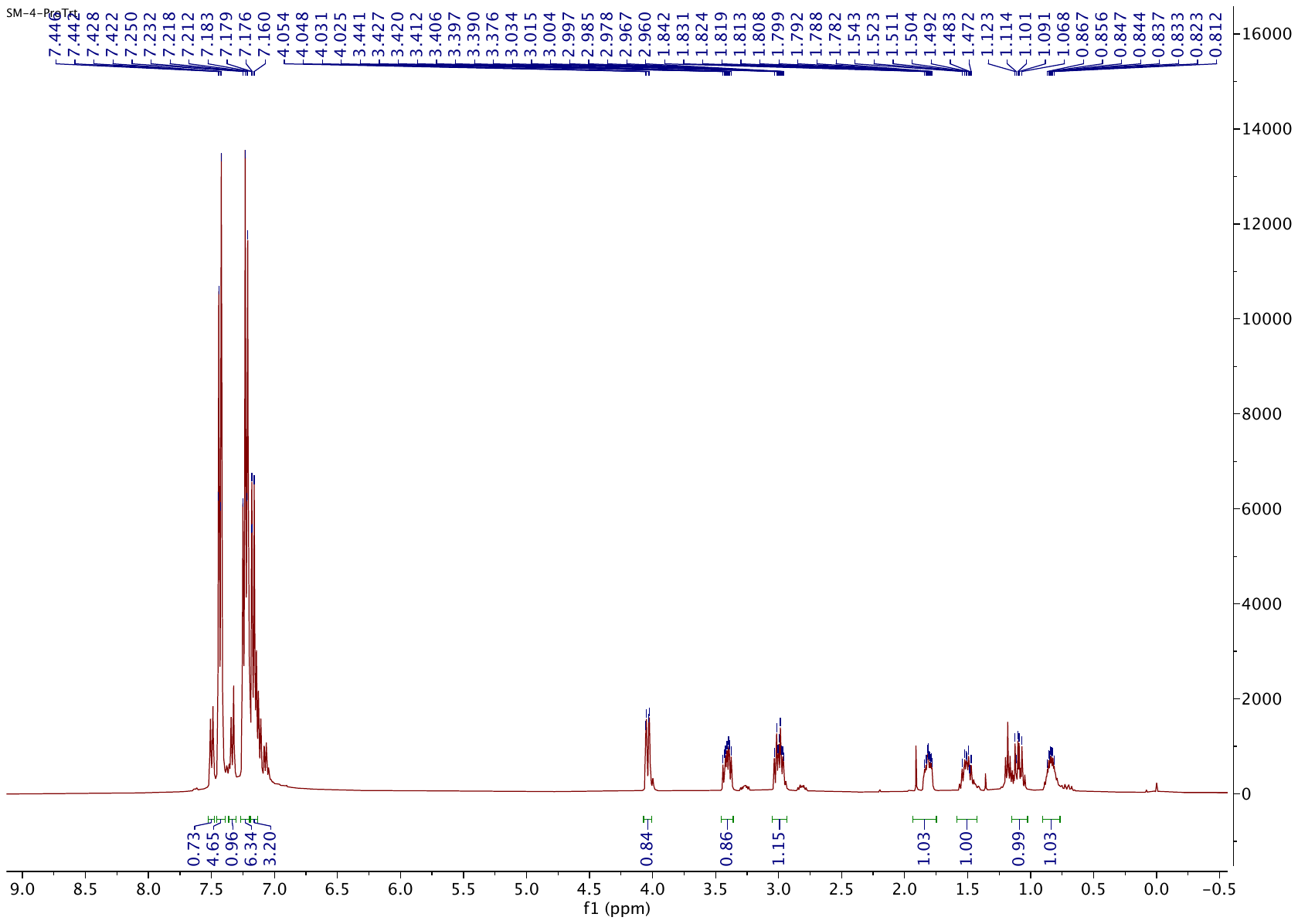

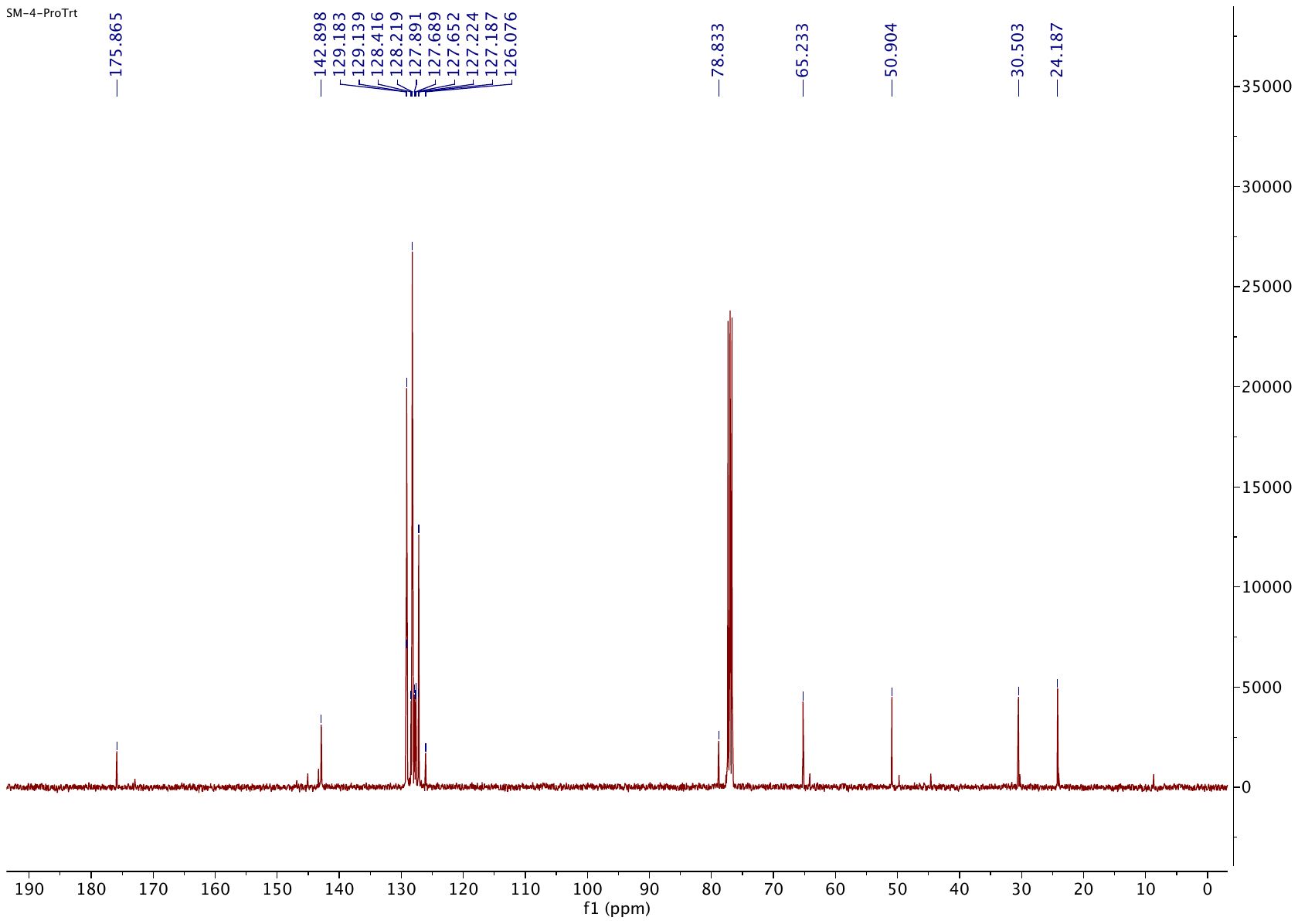

**Fig. S13. Valine transfer in a tRNA acceptor stem SM-1.**Duplex sequence:

5’UCGCUUUCCA

3’AGCGAp-L-Val;

Transfer was monitored using HPLC with 260 nm UV detection. The solution was incubated at 10°C and aliquots of 8 μL were injected into an HPLC at different time points. Peaks for the donor, the donor mixed anhydride and acceptor strands are indicated. The peak presumed to be due to the diol ester transfer product is highlighted by the dashed box. Conditions: both oligos (100 μM), NaCl (100 mM), MgCl_2_ (5 mM), HEPES (50 mM, pH 6.8).

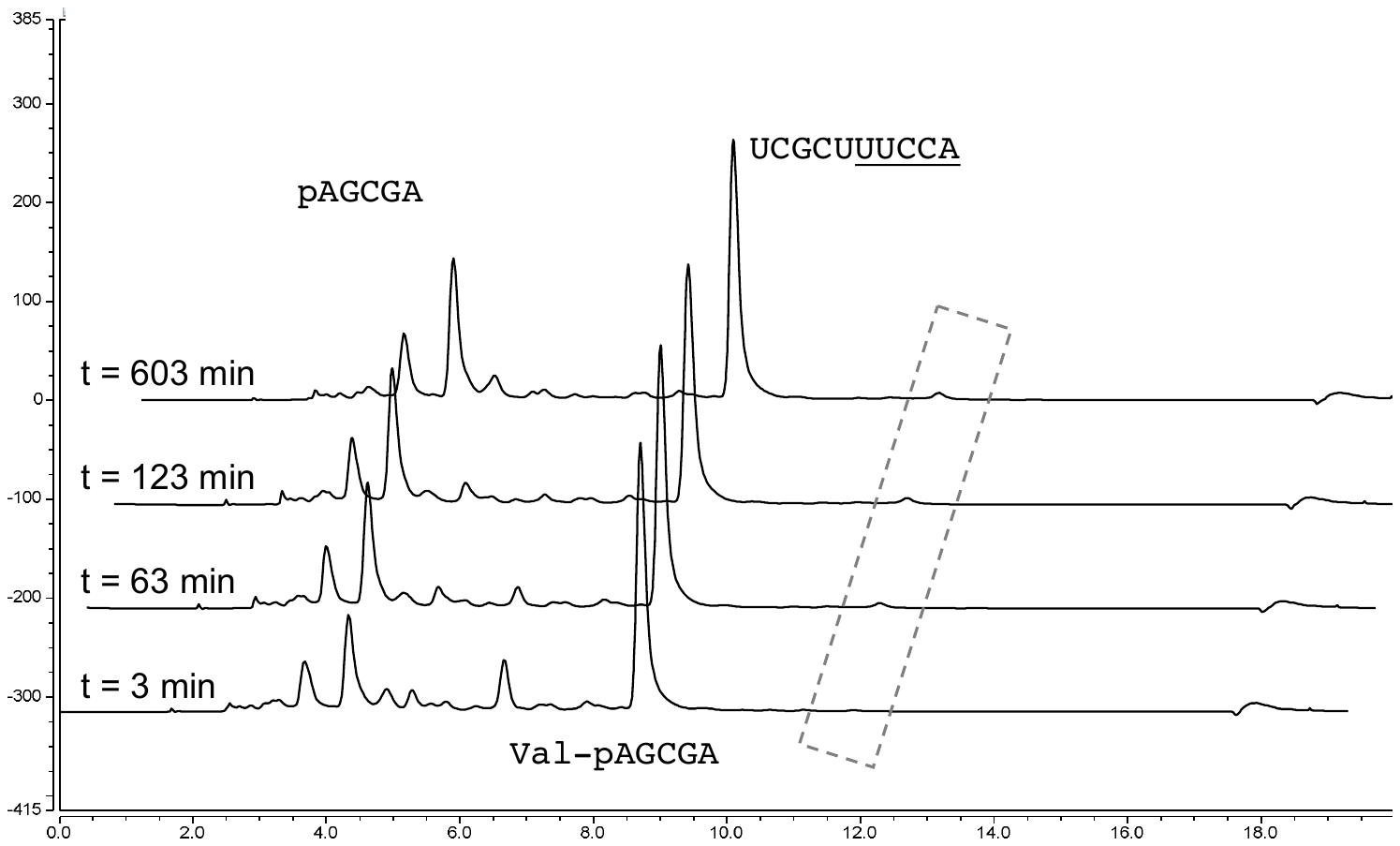

**Fig. S14. Leucine transfer in a tRNA acceptor stem SM-1.** Duplex sequence:

5’UCGCUUUCCA

3’AGCGAp-L-Leu;

Transfer was monitored using HPLC with 260 nm UV detection. The solution was incubated at 10°C and aliquots of 8 μL were injected into an HPLC at different time points. Peaks for the donor, the donor mixed anhydride and acceptor strands are indicated. The peak presumed to be due to the diol ester transfer product is highlighted by the dashed box. Conditions: both oligos (100 μM), NaCl (100 mM), MgCl_2_ (5 mM), HEPES (50 mM, pH 6.8).

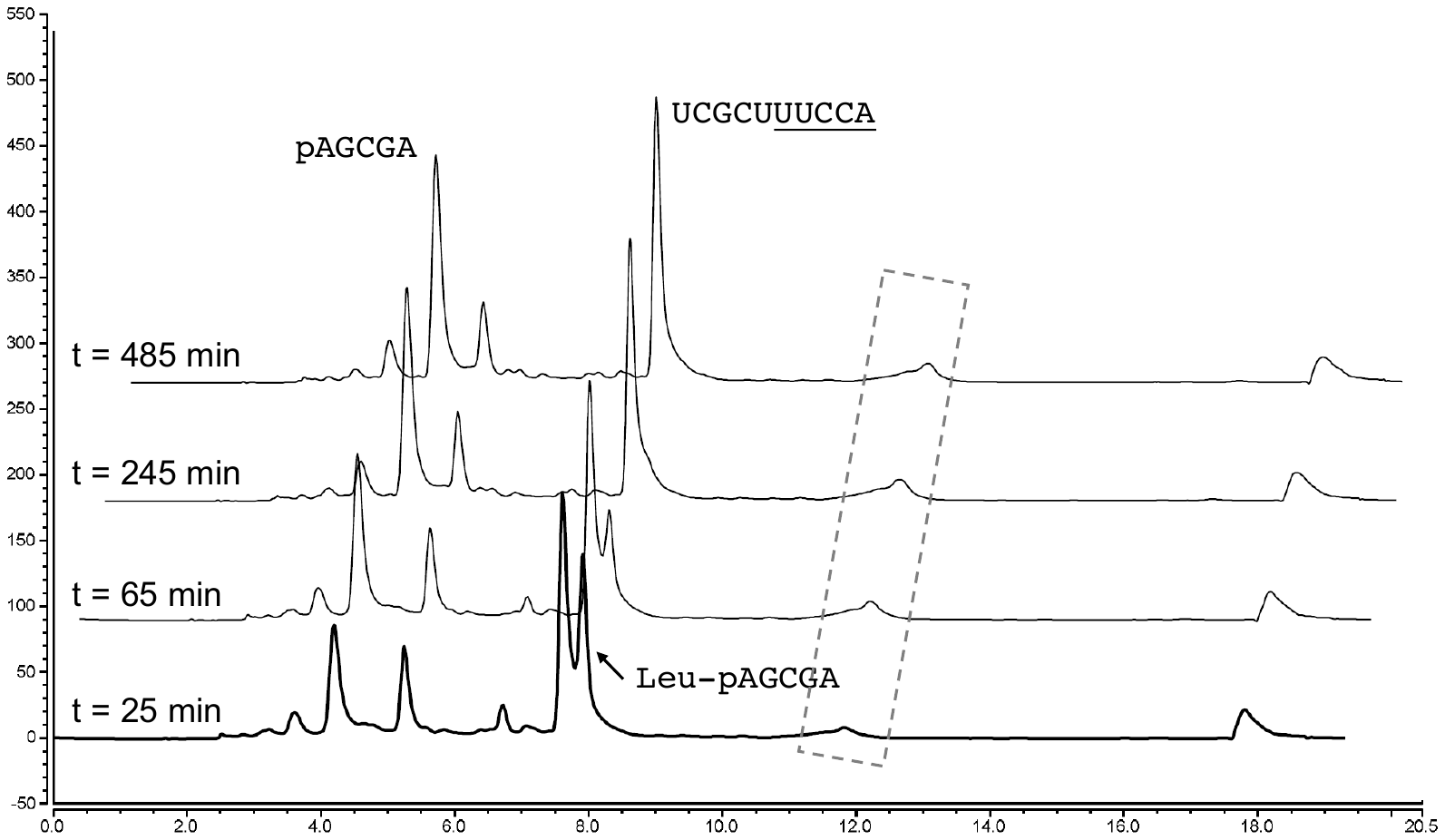

**Fig. S15. Proline transfer in a tRNA acceptor stem SM-1.** Duplex sequence:

5’UCGCUUUCCA

3’AGCGAp-L-Pro;

Transfer was monitored using HPLC with 260 nm UV detection. The solution was incubated at 10°C and aliquots of 8 μL were injected into an HPLC at different time points. Peaks for the donor, the donor mixed anhydride and acceptor strands are indicated. The peak presumed to be due to the diol ester transfer product is highlighted by the dashed box. Conditions: both oligos (100 μM), NaCl (100 mM), MgCl_2_ (5 mM), HEPES (50 mM, pH 6.8).

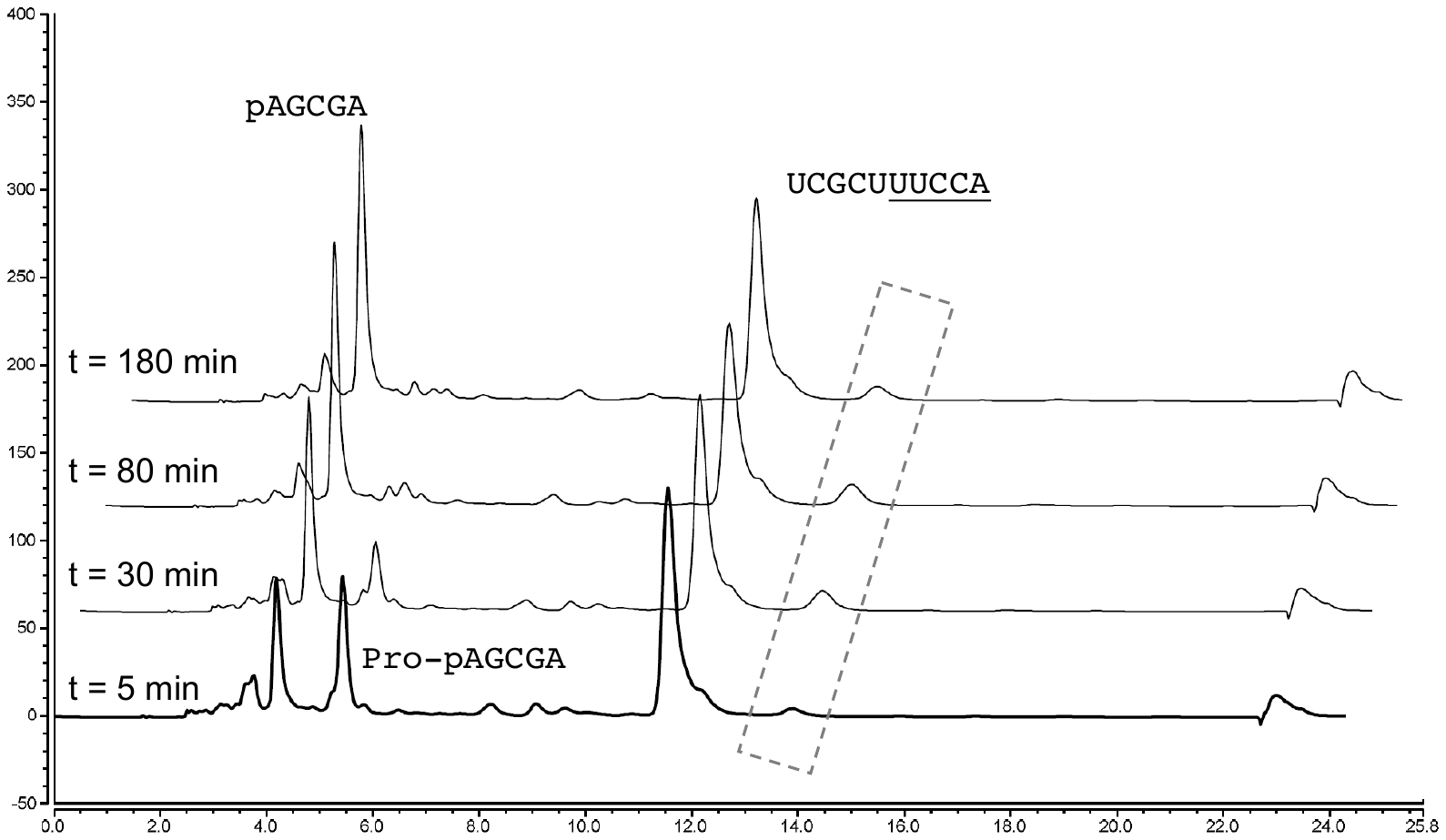

**Table S1.** Sequence of the donor and acceptor strands used in the mixed anhydride chemistry. Found, observed (raw) mass number from LCMS in negative ion mode; Expt., the experimental mass number calculated from the observed values, i.e. (Found + 1) × charges, 2 charges for 5-mer Aa donors and 3 charges for 10-mer acceptors; Calc., mass number in theory; Acc, acceptor.

| **Stem** | | **Sequence** | **mer** | **Calc.** | **Expt.** | **Found** |
| --- | --- | --- | --- | --- | --- | --- |
| **SM-1** | SM-1-Acc | UCGCU UUCCA | 10 | 3056.4 | 3056.7 | 1017.9 |
|  | Acc-1A | UCGCU AGCCA | 10 | 3118.5 | 3118.8 | 1038.6 |
|  | Acc-1C | UCGCU CGCCA | 10 | 3094.5 | 3094.5 | 1030.5 |
|  | Acc-1G | UCGCU GGCCA | 10 | 3134.5 | 3134.4 | 1043.8 |
|  | Acc-2A | UCGCU UACCA | 10 | 3079.4 | 3079.2 | 1025.4 |
|  | Acc-2C | UCGCU UCCCA | 10 | 3055.4 | 3055.5 | 1017.5 |
|  | Acc-2U | UCGCU UUCCA | 10 | 3056.4 | 3056.7 | 1017.9 |
|  | Acc-3A | UCGCU UGACA | 10 | 3119.4 | 3119.4 | 1038.8 |
|  | Acc-3G | UCGCU UGGCA | 10 | 3135.3 | 3135.3 | 1044.1 |
|  | Acc-3U | UCGCU UGUCA | 10 | 3096.4 | 3096.3 | 1031.1 |
|  | Acc-UA | UCGCU UUACA | 10 | 3080.4 | 3080.1 | 1025.7 |
|  | Acc-UG | UCGCU UUGCA | 10 | 3096.4 | 3096.3 | 1031.1 |
|  | Acc-UU | UCGCU UUUCA | 10 | 3057.4 | 3057.0 | 1018.0 |
|  | Acc-4A | UCGCU UGCAA | 10 | 3119.5 | 3119.4 | 1038.8 |
|  | Acc-4G | UCGCU UGCGA | 10 | 3135.5 | 3135.6 | 1044.2 |
|  | Acc-4U | UCGCU UGCUA | 10 | 3096.4 | 3096.9 | 1031.3 |
|  | Acc-5C | UCGCU UGCCC | 10 | 3071.4 | 3071.7 | 1022.9 |
|  | Acc-5G | UCGCU UGCCG | 10 | 3111.4 | 3111.3 | 1036.1 |
|  | Acc-5U | UCGCU UGCCU | 10 | 3072.4 | 3072.6 | 1023.2 |
|  | SM-1 | pAGCGA | 5 | 1671.3 | 1671.2 | 834.6 |
|  | SM-1-Gly | Gly-pAGCGA | 5 | 1728.3 | 1728.0 | 863.0 |
|  | SM-1-Ala | Ala-pAGCGA | 5 | 1742.3 | 1742.0 | 870.0 |
|  | SM-1-Val | Val-pAGCGA | 5 | 1770.4 | 1768.8 | 883.4 |
|  | SM-1-Leu | Leu-pAGCGA | 5 | 1784.4 | 1784.0 | 891.0 |
|  | SM-1-Pro | Pro-pAGCGA | 5 | 1768.4 | 1768.2 | 883.1 |
| **SM-2** | SM-2-Acc | UUGCU UUCCA | 10 | 3057.4 | 3057.3 | 1018.1 |
|  | SM-2 | pAGCAA | 5 | 1655.3 | 1655.0 | 826.5 |
|  | SM-2-Gly | Gly-pAGCAA | 5 | 1712.3 | 1712.0 | 855.0 |
|  | SM-2-Ala | Ala-pAGCAA | 5 | 1726.3 | 1726.0 | 862.0 |
|  | SM-2-Val | Val-pAGCAA | 5 | 1754.4 | 1754.0 | 876.0 |
|  | SM-2-Leu | Leu-pAGCAA | 5 | 1768.4 | 1768.0 | 883.0 |
|  | SM-2-Pro | Pro-pAGCAA | 5 | 1752.4 | 1752.0 | 875.0 |
| **SM-3** | SM-3-Acc | UCACU UUCCA | 10 | 3040.4 | 3040.2 | 1012.4 |
|  | SM-3 | pAGUGA | 5 | 1672.2 | 1672.0 | 835.0 |
|  | SM-3-Gly | Gly-pAGUGA | 5 | 1729.2 | 1729.0 | 863.5 |
|  | SM-3-Ala | Ala-pAGUGA | 5 | 1743.2 | 1743.0 | 870.5 |
|  | SM-3-Val | Val-pAGUGA | 5 | 1771.4 | 1771.0 | 884.5 |
|  | SM-3-Leu | Leu-pAGUGA | 5 | 1785.3 | 1785.0 | 891.5 |
|  | SM-3-Pro | Pro-pAGUGA | 5 | 1769.4 | 1769.0 | 883.5 |
| **SM-4** | SM-4-Acc | UGCCU UUCCA | 10 | 3056.4 | 3056.4 | 1017.8 |
|  | SM-4 | pAGGCA | 5 | 1671.3 | 1671.0 | 834.5 |
|  | SM-4-Gly | Gly-pAGGCA | 5 | 1728.3 | 1727.8 | 862.9 |
|  | SM-4-Ala | Ala-pAGGCA | 5 | 1742.3 | 1741.8 | 869.9 |
|  | SM-4-Val | Val-pAGGCA | 5 | 1770.4 | 1770.0 | 884.0 |
|  | SM-4-Leu | Leu-pAGGCA | 5 | 1784.4 | 1784.0 | 891.0 |
|  | SM-4-Pro | Pro-pAGGCA | 5 | 1768.4 | 1768.0 | 883.0 |
| **SM-5** | SM-5-Acc | UCGUU UUCCA | 10 | 3057.4 | 3057.3 | 1018.1 |
|  | SM-5 | pAACGA | 5 | 1655.3 | 1655.0 | 826.5 |
|  | SM-5-Gly | Gly-pAACGA | 5 | 1712.3 | 1711.8 | 854.9 |
|  | SM-5-Ala | Ala-pAACGA | 5 | 1726.3 | 1726.0 | 862.0 |
|  | SM-5-Val | Val-pAACGA | 5 | 1754.4 | 1754.0 | 876.0 |
|  | SM-5-Leu | Leu-pAACGA | 5 | 1768.4 | 1768.0 | 883.0 |
|  | SM-5-Pro | Pro-pAACGA | 5 | 1752.4 | 1751.8 | 874.9 |
| **SM-6** | SM-6-Acc | UUGCG UUCCA | 10 | 3096.4 | 3096.3 | 1031.1 |
|  | SM-6 | pCGCAA | 5 | 1631.2 | 1631.0 | 814.5 |
|  | SM-6-Gly | Gly-pCGCAA | 5 | 1688.2 | 1688.0 | 843.0 |
|  | SM-6-Ala | Ala-pCGCAA | 5 | 1702.2 | 1702.0 | 850.0 |
|  | SM-6-Val | Val-pCGCAA | 5 | 1730.4 | 1730.0 | 864.0 |
|  | SM-6-Leu | Leu-pCGCAA | 5 | 1744.3 | 1744.0 | 871.0 |
|  | SM-6-Pro | Pro-pCGCAA | 5 | 1728.4 | 1728.0 | 863.0 |

| **Stem** | | | **Sequence** | | **mer** | | **Calc.** | **Expt.** | | **Found** |
| --- | --- | --- | --- | --- | --- | --- | --- | --- | --- | --- |
| **SM-11** | SM-11-Acc | CGCAG UUCCA | | 10 | | 3118.5 | | 3118.5 | 1038.5 | |
|  | SM-11U-Acc | CGUAG UUCCA | | 10 | | 3119.5 | | 3120.0 | 1039.0 | |
|  | SM-11 | pCUGCG | | 5 | | 1624.2 | | 1624.0 | 811.0 | |
|  | SM-11-Gly | pCUGCG | | 5 | | 1681.2 | | 1681.0 | 839.5 | |
|  | SM-11-L-Ala | pCUGCG | | 5 | | 1695.2 | | 1693.4 | 845.7 | |
|  | SM-11-D-Ala | pCUGCG | | 5 | | 1695.2 | | 1695.0 | 846.5 | |
|  | SM-11-Val | pCUGCG | | 5 | | 1723.4 | | 1723.0 | 860.5 | |
|  | SM-11-Leu | pCUGCG | | 5 | | 1737.3 | | 1735.6 | 866.8 | |
|  | SM-11-Pro | pCUGCG | | 5 | | 1721.4 | | 1720.8 | 859.4 | |
| **SM-12** | SM-12-Acc | CGCUC UUCCA | | 10 | | 3055.4 | | 3055.2 | 1017.4 | |
|  | SM-12 | pGAGCG | | 5 | | 1687.3 | | 1687.0 | 842.5 | |
|  | SM-12-Gly | pGAGCG | | 5 | | 1744.3 | | 1743.8 | 870.9 | |
|  | SM-12-L-Ala | pGAGCG | | 5 | | 1758.3 | | 1758.0 | 878.0 | |
|  | SM-12-D-Ala | pGAGCG | | 5 | | 1758.3 | | 1758.0 | 878.0 | |
|  | SM-12-Val | pGAGCG | | 5 | | 1786.4 | | 1786.0 | 892.0 | |
|  | SM-12-Leu | pGAGCG | | 5 | | 1800.4 | | 1800.4 | 899.2 | |
|  | SM-12-Pro | pGAGCG | | 5 | | 1784.4 | | 1783.8 | 890.9 | |
| **SM-13** | SM-13-Acc | CGGUA UUCCA | | 10 | | 3119.5 | | 3119.4 | 1038.8 | |
|  | SM-13 | pUACCG | | 5 | | 1608.2 | | 1608.0 | 803.0 | |
|  | SM-13-Gly | pUACCG | | 5 | | 1665.2 | | 1665.2 | 831.6 | |
|  | SM-13-L-Ala | pUACCG | | 5 | | 1679.2 | | 1679.6 | 838.8 | |
|  | SM-13-D-Ala | pUACCG | | 5 | | 1679.2 | | 1679.0 | 838.5 | |
|  | SM-13-Val | pUACCG | | 5 | | 1707.4 | | 1706.8 | 852.4 | |
|  | SM-13-Leu | pUACCG | | 5 | | 1721.3 | | 1719.4 | 858.7 | |
|  | SM-13-Pro | pUACCG | | 5 | | 1705.4 | | 1705.0 | 851.5 | |
| **SM-14** | SM-14-Acc | CGGAG UUCCA | | 10 | | 3158.5 | | 3158.4 | 1051.8 | |
|  | SM-14 | pCUCCG | | 5 | | 1584.2 | | 1584.0 | 791.0 | |
|  | SM-14-Gly | pCUCCG | | 5 | | 1641.2 | | 1641.0 | 819.5 | |
|  | SM-14-L-Ala | pCUCCG | | 5 | | 1655.2 | | 1655.2 | 826.6 | |
|  | SM-14-D-Ala | pCUCCG | | 5 | | 1655.2 | | 1654.8 | 826.4 | |
|  | SM-14-Val | pCUCCG | | 5 | | 1683.4 | | 1683.0 | 840.5 | |
|  | SM-14-Leu | pCUCCG | | 5 | | 1697.4 | | 1696.8 | 847.4 | |
|  | SM-14-Pro | pCUCCG | | 5 | | 1681.3 | | 1681.0 | 839.5 | |
| **SM-15** | SM-15-Acc | CGUCU UUCCA | | 10 | | 3056.4 | | 3056.4 | 1017.8 | |
|  | SM-15 | pAGACG | | 5 | | 1671.3 | | 1671.2 | 834.6 | |
|  | SM-15-Gly | pAGACG | | 5 | | 1728.3 | | 1728.0 | 863.0 | |
|  | SM-15-L-Ala | pAGACG | | 5 | | 1742.3 | | 1742.0 | 870.0 | |
|  | SM-15-D-Ala | pAGACG | | 5 | | 1742.3 | | 1741.8 | 869.9 | |
|  | SM-15-Val | pAGACG | | 5 | | 1770.4 | | 1770.4 | 884.2 | |
|  | SM-15-Leu | pAGACG | | 5 | | 1784.4 | | 1784.0 | 891.0 | |
|  | SM-15-Pro | pAGACG | | 5 | | 1768.4 | | 1768.2 | 883.1 | |
| **SM-16** | SM-16 | pCUGC | | 4 | | 1279.2 | | 1279.0 | 638.5 | |
|  | SM-16-L-Ala | pCUGC | | 4 | | 1350.2 | | 1350.0 | 674.0 | |
|  | SM-16-Leu | pCUGC | | 4 | | 1392.4 | | 1392.0 | 695.0 | |
|  | SM-16-Pro | pCUGC | | 4 | | 1376.3 | | 1376.0 | 687.0 | |
| **SM-17** | SM-17 | pCUG | | 3 | | 974.1 | | 973.9 | 972.9 | |
|  | SM-17-L-Ala | pCUG | | 3 | | 1045.1 | | 1044.9 | 1043.9 | |
|  | SM-17-Leu | pCUG | | 3 | | 1087.3 | | 1086.9 | 1085.9 | |
|  | SM-17-Pro | pCUG | | 3 | | 1071.3 | | 1070.9 | 1069.9 | |
| **SM-18** | SM-18-Acc | GUUAG UUCCA | | 10 | | 3120.4 | | 3120.3 | 1039.1 | |
|  | SM-18 | pCUAAC | | 5 | | 1592.2 | | 1592.4 | 795.2 | |
|  | SM-18-Ala | pCUAAC | | 5 | | 1663.2 | | 1663.0 | 830.5 | |
| **SM-19** | SM-19-Acc | CAAGU UUCCA | | 10 | | 3103.5 | | 3103.2 | 1033.4 | |
|  | SM-19 | pACUUG | | 5 | | 1609.2 | | 1609.0 | 803.5 | |
|  | SM-19-Leu | pACUUG | | 5 | | 1722.4 | | 1722.0 | 860.0 | |

**Table S2.** Kinetic data of L-alanyl-transfer with varied overhangs using mixed anhydride chemistry. Aminoacyl-transfer reactions were conducted at 10^o^C in 50 mM HEPES, 100 mM NaCl, 5 mM MgCl_2_, pH = 6.8 buffer with 100 µM equal equivalent of donor strand and acceptor strand, unless otherwise noted in **Fig. 3H**. UGCCA, UGACA and UGUCA shows similar transfer yield and kinetics. This is due to the overhang palindromic structure where nicked duplex transfer occurred, along with the self-folding nicked loop transfer.

|  | **Donor** | **Acceptor** | | | **Observed yield** | **Corrected yield** | **Mixed anhyd.** | **Diol ester** | **Peak time** | **k transfer** | **k hydrolysis** |
| --- | --- | --- | --- | --- | --- | --- | --- | --- | --- | --- | --- |
|  |  | **Stem** | **Overhang** | |  |  | **half-life** /h | **half-life** /h | min | /min^-1^ | /min^-1^ |
| **SM-1** | L-Ala-pAGCGA | UCGCU | | UGCCA | 30% | 55% | 0.2 | 4.1 | 43 | 0.0780 | 0.00279 |
|  |  |  | | AGCCA | 0% | 0% | 0.8 | -- | -- | -- | -- |
|  |  |  | | CGCCA | 6% | 9% | 0.8 | 20 | 89 | 0.0508 | 0.00058 |
|  |  |  | | GGCCA | 0% | 0% | -- | -- | -- | -- | -- |
|  |  |  | | UACCA | 16% | 30% | 0.4 | 12 | 80 | 0.0503 | 0.00095 |
|  |  |  | | UCCCA | 0% | 0% | 0.2 | -- | -- | -- | -- |
|  |  |  | | UUCCA | 34% | 57% | 0.5 | 10 | 70 | 0.0569 | 0.00116 |
|  |  |  | | UGACA | 23% | 56% | 0.2 | 5.0 | 32 | 0.1278 | 0.00229 |
|  |  |  | | UGGCA | 0% | 0% | 0.5 | -- | -- | -- | -- |
|  |  |  | | UGUCA | 24% | 59% | 0.1 | 1.8 | 21 | 0.1585 | 0.00648 |
|  |  |  | | UUACA | 28% | 39% | 0.3 | 6.1 | 52 | 0.0717 | 0.00190 |
|  |  |  | | UUGCA | 0% | 0% | 0.2 | -- | -- | -- | -- |
|  |  |  | | UUUCA | 35% | 45% | 0.2 | 7.5 | 42 | 0.1008 | 0.00153 |
|  |  |  | | UGCAA | 0% | 0% | 0.2 | -- | -- | -- | -- |
|  |  |  | | UGCGA | 0% | 0% | 0.8 | -- | -- | -- | -- |
|  |  |  | | UGCUA | 6% | 10% | 0.4 | -- | -- | -- | -- |
|  |  |  | | UGCCC | 0% | 0% | 0.8 | -- | -- | -- | -- |
|  |  |  | | UGCCG | 14% | 25% | 0.3 | 6.8 | 61 | 0.0599 | 0.00169 |
|  |  |  | | UGCCU | 0% | 0% | 0.5 | -- | -- | -- | -- |
|  | L-Ala-pAGCGA | -- | | -- | -- | -- | 0.9 | -- | -- | -- | -- |

**Table S3.** Yield data of L/D-amino acid-transfer with varied overhangs using phosphoramidate-ester chemistry. Esterification reactions were conducted at room temperature for 18 hours in 50 mM HEPES, 200 mM NaCl, 50 mM MgCl_2_, 10 mM imidazole, 50 mM EDC hydrochloride pH = 7.0 buffer with 100 µM equal equivalent of donor strand and acceptor strand and 120 µM cytidine as an internal standard. All results from single experiments with exception of those performed with UUCCA overhang which were performed in triplicate and previously reported.

|  | **Donor** | **Acceptor** | | | **Corrected yield** |
| --- | --- | --- | --- | --- | --- |
|  |  | **Stem** | **Overhang** | |  |
| **SM-1** | L-Leu-pAGCGA | UCGCU | | UGCCA | 4.0% |
|  |  |  | | UACCA | 0.7% |
|  |  |  | | UCCCA | 0.9% |
|  |  |  | | UUCCA | 1.9% |
|  |  |  | | UGACA | 6.1% |
|  |  |  | | UGGCA | 8.1% |
|  |  |  | | UGUCA | 6.8% |
|  |  |  | | UGCAA | 0.6% |
|  |  |  | | UGCUA | 1.4% |
|  | D-Leu-pAGCGA | UCGCU | | UUCCA | 0.2% |
|  | L-Ala-pAGCGA | UCGCU | | UUCCA | 1.6% |
|  | D-Ala-pAGCGA | UCGCU | | UUCCA | 0.2% |
|  | L-Val-pAGCGA | UCGCU | | UUCCA | 1.0% |
|  | D-Val-pAGCGA | UCGCU | | UUCCA | 0.2% |
|  | L-Ser-pAGCGA | UCGCU | | UUCCA | 7.2% |
|  | D-Ser-pAGCGA | UCGCU | | UUCCA | 1.6% |
|  | Gly-pAGCGA | UCGCU | | UUCCA | 13% |
|  | L-Pro-pAGCGA | UCGCU | | UUCCA | 0.4% |
|  | L-Arg-pAGCGA | UCGCU | | UUCCA | 0.8% |
|  | pAGCGA | UCGCU | | UUCCA | 0% |

**Table S4.** Synthesized oligo(deoxy)ribonucleotides for Next Generation Sequencing experiments. Set a is used for mixed anhydride chemistry; set b is used for phosphoramidate-ester chemistry.

|  | **Sequence (5'-3')** | **Length** |
| --- | --- | --- |
| Lib-L5 a | GAAGACGGCAUACGAGAU UCGCU NNNNN | 28 |
| Lib-L5 b | [Cy5]GAUAAUACGACUCACAGCAGCGUCCAUCCACA UCGCU NNNNN | 42 |
| Lib-S3 a | CAGAAGACGGCAUACGA GAUUCNNN UUCCA | 30 |
| Lib-S3 b | [Cy5]GAUAAUACGACUCACAGCAGCGUCCAUCC GAUUCNNN UUCCA | 42 |
| Lib-S3-D | pNNNGAAUC | 8 |
| Blocker a | ATCTCGTATGCCGTCTT[SpcC3] | 17 |
| Ligator a* | App[N]_8-12_AGATCGGAAGAGCGTCGTGTAGGGAAAGAGTGT[SpcC3] | 41 – 45 |
| RT primer a/  Fwd primer a | AATGATACGGCGACCACCGAGATCTACACXXXXXXACACTCTTTCCCTACACGACG | 56 |
| Rev primer a | CAAGCAGAAGACGGCATACGAGAT | 24 |
| Blocker b | GGATGGACGCTGCTGTGAGTCGTATTATC[SpcC3] | 29 |
| Ligator b | pUCUACCUCAUUCUUCACUGGAGACUUGACGAAGCUG[Cy3] | 36 |
| RT primer b | TACGCCATTCGAGATCCTCATGCAGCTTCGTCAAGTCTCC | 40 |
| Fwd primer b | CAAGCAGAAGACGGCATACGAGATXXXXXXXXGTCTCGTGGGCTCGGAGATGTGTATAAGAGACAGTACGCCATTCGAGATCCTCATGCA | 90 |
| Rev primer b | AATGATACGGCGACCACCGAGATCTACACXXXXXXXXTCGTCGGCAGCGTCAGATGTGTATAAGAGACAGGATAATACGACTCACAGCAGCGTC | 94 |

N, randomized nucleotides; RT, reverse transcription; X, index sequence; *, this DNA oligo is adenylated at its 5**'**-end through the technique which is being described above using ImpA. Six different oligos with variably long stretches of randomized nucleotides on their 5**'**-end were used. By this, frameshift between the samples can be achieved, increasing complexity during sequencing which ultimately increases the quality of the generated data. The randomized nucleotides also allow for the identification of PCR duplicates.

**Table S5.** Rank of quality-filtered read numbers of CCA-ending overhangs from the **Lib-L5** sequencing results (mixed anhydride chemistry). All 4^5^ = 1024 overhangs were observed.

|  | **Blank** | **L-Ala** | **Gly** | **Pro** | **Leu** | **Val** |
| --- | --- | --- | --- | --- | --- | --- |
| **UUCCA** | **709** | **98** | **29** | **27** | **63** | **18** |
| UA**CCA** | 761 | 840 | 848 | 815 | 887 | 902 |
| AU**CCA** | 794 | 894 | 927 | 898 | 921 | 1008 |
| AG**CCA** | 864 | 853 | 1007 | 963 | 977 | 953 |
| AA**CCA** | 891 | 994 | 985 | 989 | 933 | 983 |
| GG**CCA** | 894 | 822 | 953 | 939 | 952 | 969 |
| GU**CCA** | 895 | 1005 | 967 | 1015 | 891 | 1019 |
| UG**CCA** | 897 | 816 | 1021 | 968 | 941 | 985 |
| CA**CCA** | 936 | 1023 | 915 | 873 | 974 | 992 |
| AC**CCA** | 965 | 737 | 863 | 825 | 928 | 910 |
| GA**CCA** | 976 | 922 | 911 | 895 | 962 | 888 |
| CU**CCA** | 985 | 755 | 583 | 741 | 741 | 810 |
| UC**CCA** | 993 | 667 | 482 | 653 | 837 | 967 |
| CG**CCA** | 998 | 1018 | 1020 | 1018 | 1018 | 1000 |
| CC**CCA** | 1022 | 992 | 806 | 897 | 865 | 931 |
| GC**CCA** | 1024 | 1017 | 1003 | 1002 | 958 | 1013 |

**Table S6.** Rank of quality-filtered read numbers of CCA-ending overhangs from the **Lib-L5** sequencing results (phosphoramidate-ester chemistry). All 4^5^ = 1024 overhangs were observed.

|  | **Blank** | **L-Ala** | **Gly** | **Pro** | **Leu** | **Val** |
| --- | --- | --- | --- | --- | --- | --- |
| **UUCCA** | **707** | **374** | **452** | **461** | **466** | **688** |
| UA**CCA** | 761 | 324 | 627 | 383 | 188 | 830 |
| AU**CCA** | 800 | 830 | 756 | 665 | 585 | 908 |
| AG**CCA** | 868 | 575 | 682 | 345 | 544 | 676 |
| UG**CCA** | 891 | 269 | 346 | 321 | 139 | 627 |
| GG**CCA** | 895 | 354 | 606 | 182 | 392 | 857 |
| GU**CCA** | 896 | 693 | 887 | 526 | 812 | 913 |
| AA**CCA** | 897 | 951 | 990 | 945 | 765 | 970 |
| CA**CCA** | 941 | 768 | 854 | 405 | 403 | 995 |
| AC**CCA** | 970 | 960 | 992 | 710 | 836 | 1015 |
| GA**CCA** | 978 | 827 | 949 | 620 | 673 | 952 |
| CU**CCA** | 985 | 250 | 214 | 59 | 229 | 635 |
| UC**CCA** | 989 | 383 | 350 | 206 | 439 | 869 |
| CG**CCA** | 998 | 321 | 240 | 116 | 266 | 832 |
| CC**CCA** | 1022 | 948 | 885 | 667 | 818 | 1020 |
| GC**CCA** | 1024 | 1009 | 1023 | 878 | 1010 | 1023 |

**Table S7.** Kinetic data of mixed anhydride aminoacyl-transfer with **SM-1**, relating to **Fig. 2A**.

|  | **Donor** | **Acceptor** | | | **Observed yield** | **Corrected yield** | **Mixed anhyd.** | **Diol ester** | **Peak time** | **k transfer** | **k hydrolysis** |
| --- | --- | --- | --- | --- | --- | --- | --- | --- | --- | --- | --- |
|  |  | **Stem** | **Overhang** | |  |  | **half-life** /h | **half-life** /h | min | /min^-1^ | /min^-1^ |
| **SM-1** | L-Ala-pAGCGA | UCGCU | | UUCCA | 34% | 57% | 0.47 | 10 | 70 | 0.0569 | 0.00116 |
|  | L-Ala-pAGCGA | -- | | -- | -- | -- | 0.90 | -- | -- | -- | -- |
|  | Gly-pAGCGA | UCGCU | | UUCCA | 2% | 3% | 0.72 | -- | -- | -- | -- |
|  | Gly-pAGCGA | -- | | -- | -- | -- | 0.93 | -- | -- | -- | -- |
|  | L-Val-pAGCGA | UCGCU | | UUCCA | 4% | 10% | 0.77 | >24 | >200 | 0.0310 | <0.00050 |
|  | L-Val-pAGCGA | -- | | -- | -- | -- | 0.95 | -- | -- | -- | -- |
|  | L-Leu-pAGCGA | UCGCU | | UUCCA | 22% | 35% | 0.72 | 16 | 157 | 0.0226 | 0.00073 |
|  | L-Leu-pAGCGA | -- | | -- | -- | -- | 0.70 | -- | -- | -- | -- |
|  | L-Pro-pAGCGA | UCGCU | | UUCCA | 14% | 27% | 0.44 | 2.8 | 55 | 0.0492 | 0.00416 |
|  | L-Pro-pAGCGA | -- | | -- | -- | -- | 0.96 | -- | -- | -- | -- |

**Table S12.** Kinetic data of mixed anhydride aminoacyl-transfer with **SM-2**, relating to **Fig. 2B**.

|  | **Donor** | **Acceptor** | | | **Observed yield** | **Corrected yield** | **Mixed anhyd.** | **Diol ester** | **Peak time** | **k transfer** | **k hydrolysis** |
| --- | --- | --- | --- | --- | --- | --- | --- | --- | --- | --- | --- |
|  |  | **Stem** | **Overhang** | |  |  | **half-life** /h | **half-life** /h | min | /min^-1^ | /min^-1^ |
| **SM-2** | L-Ala-pAGCAA | UUGCU | | UUCCA | 15% | 54% | 0.59 | 11 | 65 | 0.0630 | 0.00109 |
|  | L-Ala-pAGCAA | -- | | -- | -- | -- | 1.27 | -- | -- | -- | -- |
|  | Gly-pAGCAA | UUGCU | | UUCCA | 0% | 0% | 0.73 | -- | -- | -- | -- |
|  | Gly-pAGCAA | -- | | -- | -- | -- | 1.12 | -- | -- | -- | -- |
|  | L-Val-pAGCAA | UUGCU | | UUCCA | 3% | 11% | 0.98 | > 24 | > 200 | 0.0137 | < 0.00050 |
|  | L-Val-pAGCAA | -- | | -- | -- | -- | 1.65 | -- | -- | -- | -- |
|  | L-Leu-pAGCAA | UUGCU | | UUCCA | 21% | 36% | 0.54 | 12 | 136 | 0.0251 | 0.00095 |
|  | L-Leu-pAGCAA | -- | | -- | -- | -- | 0.76 | -- | -- | -- | -- |
|  | L-Pro-pAGCAA | UUGCU | | UUCCA | 21% | 29% | 0.30 | 3.9 | 78 | 0.0343 | 0.00296 |
|  | L-Pro-pAGCAA | -- | | -- | -- | -- | 0.96 | -- | -- | -- | -- |

**Table S10.** Kinetic data of mixed anhydride aminoacyl-transfer with **SM-3**, relating to **Fig. 2C**.

|  | **Donor** | **Acceptor** | | | **Observed yield** | **Corrected yield** | **Mixed anhyd.** | **Diol ester** | **Peak time** | **k transfer** | **k hydrolysis** |
| --- | --- | --- | --- | --- | --- | --- | --- | --- | --- | --- | --- |
|  |  | **Stem** | **Overhang** | |  |  | **half-life** /h | **half-life** /h | min | /min^-1^ | /min^-1^ |
| **SM-3** | L-Ala-pAGUGA | UCACU | | UUCCA | 17% | 30% | 0.49 | 4.2 | 117 | 0.0195 | 0.00273 |
|  | L-Ala-pAGUGA | -- | | -- | -- | -- | 1.20 | -- | -- | -- | -- |
|  | Gly-pAGUGA | UCACU | | UUCCA | 7% | 10% | 1.22 | > 24 | > 200 | 0.0236 | < 0.00050 |
|  | Gly-pAGUGA | -- | | -- | -- | -- | 1.63 | -- | -- | -- | -- |
|  | L-Val-pAGUGA | UCACU | | UUCCA | 7% | 21% | 0.59 | > 24 | > 200 | 0.0219 | < 0.00050 |
|  | L-Val-pAGUGA | -- | | -- | -- | -- | 1.02 | -- | -- | -- | -- |
|  | L-Leu-pAGUGA | UCACU | | UUCCA | 20% | 40% | 0.37 | 11 | 110 | 0.0320 | 0.00105 |
|  | L-Leu-pAGUGA | -- | | -- | -- | -- | 0.96 | -- | -- | -- | -- |
|  | L-Pro-pAGUGA | UCACU | | UUCCA | 21% | 26% | 0.45 | 3.3 | 69 | 0.0382 | 0.00348 |
|  | L-Pro-pAGUGA | -- | | -- | -- | -- | 1.01 | -- | -- | -- | -- |

**Table S11.** Kinetic data of mixed anhydride aminoacyl-transfer with **SM-4**, relating to **Fig. 2D**.

|  | **Donor** | **Acceptor** | | | **Observed yield** | **Corrected yield** | **Mixed anhyd.** | **Diol ester** | **Peak time** | **k transfer** | **k hydrolysis** |
| --- | --- | --- | --- | --- | --- | --- | --- | --- | --- | --- | --- |
|  |  | **Stem** | **Overhang** | |  |  | **half-life** /h | **half-life** /h | min | /min^-1^ | /min^-1^ |
| **SM-4** | L-Ala-pAGGCA | UGCCU | | UUCCA | 18% | 30% | 0.41 | 6.1 | 68 | 0.0501 | 0.00189 |
|  | L-Ala-pAGGCA | -- | | -- | -- | -- | 0.67 | -- | -- | -- | -- |
|  | Gly-pAGGCA | UGCCU | | UUCCA | 0% | 0% | 0.88 | -- | -- | -- | -- |
|  | Gly-pAGGCA | -- | | -- | -- | -- | 1.61 | -- | -- | -- | -- |
|  | L-Val-pAGGCA | UGCCU | | UUCCA | 3% | 10% | 0.94 | > 24 | > 200 | 0.0121 | < 0.00050 |
|  | L-Val-pAGGCA | -- | | -- | -- | -- | 1.06 | -- | -- | -- | -- |
|  | L-Leu-pAGGCA | UGCCU | | UUCCA | 21% | 48% | 0.37 | 10 | 97 | 0.0369 | 0.00116 |
|  | L-Leu-pAGGCA | -- | | -- | -- | -- | 0.79 | -- | -- | -- | -- |
|  | L-Pro-pAGGCA | UGCCU | | UUCCA | 21% | 48% | 0.35 | 3.1 | 44 | 0.0714 | 0.00376 |
|  | L-Pro-pAGGCA | -- | | -- | -- | -- | 1.17 | -- | -- | -- | -- |

**Table S9.** Kinetic data of mixed anhydride aminoacyl-transfer with **SM-5**, relating to **Fig. 2E**.

|  | **Donor** | **Acceptor** | | | **Observed yield** | **Corrected yield** | **Mixed anhyd.** | **Diol ester** | **Peak time** | **k transfer** | **k hydrolysis** |
| --- | --- | --- | --- | --- | --- | --- | --- | --- | --- | --- | --- |
|  |  | **Stem** | **Overhang** | |  |  | **half-life** /h | **half-life** /h | min | /min^-1^ | /min^-1^ |
| **SM-5** | L-Ala-pAACGA | UCGUU | | UUCCA | 15% | 36% | 0.54 | 6.4 | 79 | 0.0411 | 0.00182 |
|  | L-Ala-pAACGA | -- | | -- | -- | -- | 0.81 | -- | -- | -- | -- |
|  | Gly-pAACGA | UCGUU | | UUCCA | 0% | 0% | 0.48 | -- | -- | -- | -- |
|  | Gly-pAACGA | -- | | -- | -- | -- | 0.58 | -- | -- | -- | -- |
|  | L-Val-pAACGA | UCGUU | | UUCCA | 5% | 16% | 0.49 | > 24 | > 200 | 0.0199 | < 0.00050 |
|  | L-Val-pAACGA | -- | | -- | -- | -- | 0.83 | -- | -- | -- | -- |
|  | L-Leu-pAACGA | UCGUU | | UUCCA | 15% | 33% | 0.43 | 8.0 | 120 | 0.0252 | 0.00145 |
|  | L-Leu-pAACGA | -- | | -- | -- | -- | 0.63 | -- | -- | -- | -- |
|  | L-Pro-pAACGA | UCGUU | | UUCCA | 20% | 31% | 0.45 | 3.5 | 71 | 0.0370 | 0.00334 |
|  | L-Pro-pAACGA | -- | | -- | -- | -- | 0.95 | -- | -- | -- | -- |

**Table S8.** Kinetic data of mixed anhydride aminoacyl-transfer with **SM-6**, relating to **Fig. 2F**.

|  | **Donor** | **Acceptor** | | | **Observed yield** | **Corrected yield** | **Mixed anhyd.** | **Diol ester** | **Peak time** | **k transfer** | **k hydrolysis** |
| --- | --- | --- | --- | --- | --- | --- | --- | --- | --- | --- | --- |
|  |  | **Stem** | **Overhang** | |  |  | **half-life** /h | **half-life** /h | min | /min^-1^ | /min^-1^ |
| **SM-6** | L-Ala-pCGCAA | UUGCG | | UUCCA | 34% | 52% | 1.16 | 13 | 156 | 0.0212 | 0.00089 |
|  | L-Ala-pCGCAA | -- | | -- | -- | -- | 1.06 | -- | -- | -- | -- |
|  | Gly-pCGCAA | UUGCG | | UUCCA | 0% | 0% | 0.71 | -- | -- | -- | -- |
|  | Gly-pCGCAA | -- | | -- | -- | -- | 1.37 | -- | -- | -- | -- |
|  | L-Val-pCGCAA | UUGCG | | UUCCA | 8% | 29% | 0.53 | > 24 | > 200 | 0.0159 | < 0.00050 |
|  | L-Val-pCGCAA | -- | | -- | -- | -- | 1.18 | -- | -- | -- | -- |
|  | L-Leu-pCGCAA | UUGCG | | UUCCA | 43% | 60% | 0.57 | 10 | 128 | 0.0257 | 0.00111 |
|  | L-Leu-pCGCAA | -- | | -- | -- | -- | 0.94 | -- | -- | -- | -- |
|  | L-Pro-pCGCAA | UUGCG | | UUCCA | 23% | 36% | 0.67 | 2.2 | 48 | 0.0526 | 0.00531 |
|  | L-Pro-pCGCAA | -- | | -- | -- | -- | 1.47 | -- | -- | -- | -- |

**Table S13.** Quality-filtered read numbers from the sequencing results of **Lib-S3** with the mixed anhydride chemistry.

|  | **Blank** | **L-Ala** | **D-Ala** | **Gly** | **Pro** | **Leu** | **Val** |
| --- | --- | --- | --- | --- | --- | --- | --- |
| AAA | 52 | 115 | 71 | 17 | 75 | 28 | 39 |
| AAC | 30 | 71 | 76 | 13 | 59 | 23 | 43 |
| AAG | 55 | 48 | 122 | 17 | 107 | 58 | 77 |
| AAU | 70 | 104 | 182 | 23 | 64 | 18 | 44 |
| ACA | 42 | 79 | 76 | 21 | 92 | 40 | 34 |
| ACC | 23 | 14 | 33 | 8 | 38 | 8 | 18 |
| ACG | 35 | 82 | 79 | 25 | 154 | 48 | 58 |
| ACU | 88 | 43 | 142 | 12 | 80 | 18 | 39 |
| AGA | 72 | 98 | 64 | 11 | 69 | 28 | 28 |
| AGC | 19 | 28 | 76 | 9 | 76 | 37 | 43 |
| AGG | 87 | 97 | 148 | 15 | 139 | 86 | 70 |
| AGU | 91 | 109 | 213 | 11 | 78 | 52 | 55 |
| AUC | 196 | 85 | 39 | 121 | 63 | 16 | 26 |
| AUG | 58 | 188 | 145 | 16 | 119 | 67 | 78 |
| AUU | 77 | 80 | 130 | 13 | 65 | 28 | 34 |
| CAA | 34 | 124 | 67 | 22 | 100 | 35 | 50 |
| CAC | 12 | 10 | 44 | 17 | 51 | 17 | 24 |
| CAG | 36 | 178 | 83 | 13 | 91 | 40 | 37 |
| CAU | 57 | 54 | 154 | 11 | 71 | 33 | 44 |
| CCA | 28 | 17 | 50 | 5 | 81 | 19 | 36 |
| CCC | 10 | 3 | 26 | 13 | 24 | 5 | 18 |
| CCG | 12 | 12 | 49 | 17 | 91 | 26 | 46 |
| CCU | 73 | 19 | 65 | 9 | 51 | 18 | 39 |
| CGA | 36 | 18 | 31 | 3 | 41 | 30 | 18 |
| CGC | 25 | 6 | 33 | 4 | 207 | 22 | 41 |
| CGG | 18 | 43 | 42 | 12 | 94 | 43 | 50 |
| CGU | 16 | 9 | 38 | 7 | 77 | 40 | 36 |
| CUA | 40 | 58 | 64 | 28 | 151 | 65 | 58 |
| CUC | 234 | 77 | 30 | 1349 | 161 | 16 | 26 |
| CUG | 54 | 70 | 127 | 108 | 163 | 44 | 65 |
| CUU | 65 | 42 | 128 | 15 | 143 | 31 | 52 |
| GAA | 69 | 77 | 58 | 43 | 60 | 40 | 57 |
| GAC | 28 | 18 | 41 | 21 | 102 | 33 | 43 |
| GAG | 121 | 68 | 133 | 99 | 177 | 116 | 108 |
| GAU | 110 | 84 | 64 | 17 | 51 | 19 | 35 |
| GCA | 62 | 14 | 85 | 29 | 163 | 64 | 67 |
| GCC | 28 | 4 | 34 | 18 | 70 | 26 | 49 |
| GCG | 81 | 30 | 134 | 137 | 273 | 95 | 111 |
| GCU | 96 | 29 | 139 | 34 | 298 | 28 | 132 |
| GGA | 80 | 26 | 46 | 32 | 82 | 37 | 42 |
| GGC | 30 | 9 | 61 | 9 | 140 | 60 | 52 |
| GGG | 133 | 69 | 134 | 35 | 262 | 95 | 109 |
| GGU | 90 | 83 | 91 | 8 | 72 | 43 | 52 |
| GUA | 116 | 6 | 30 | 9 | 178 | 78 | 45 |
| GUC | 23 | 1 | 12 | 17 | 85 | 28 | 43 |
| GUG | 176 | 40 | 70 | 29 | 221 | 93 | 79 |
| GUU | 40 | 23 | 36 | 10 | 87 | 41 | 61 |
| UAA | 69 | 141 | 75 | 15 | 111 | 37 | 64 |
| UAA | 83 | 170 | 89 | 33 | 137 | 45 | 93 |
| UAC | 48 | 41 | 107 | 19 | 98 | 28 | 53 |
| UAG | 64 | 234 | 188 | 30 | 180 | 68 | 114 |
| UAU | 79 | 98 | 113 | 24 | 97 | 37 | 56 |
| UCA | 52 | 40 | 110 | 22 | 147 | 48 | 69 |
| UCC | 56 | 7 | 55 | 4 | 50 | 10 | 23 |
| UCG | 86 | 10 | 57 | 11 | 133 | 37 | 67 |
| UCU | 179 | 48 | 175 | 95 | 375 | 32 | 66 |
| UGA | 93 | 30 | 117 | 55 | 58 | 56 | 67 |
| UGC | 150 | 14 | 212 | 49 | 153 | 45 | 78 |
| UGG | 59 | 98 | 204 | 62 | 238 | 74 | 96 |
| UGU | 149 | 69 | 129 | 29 | 178 | 69 | 61 |
| UUA | 75 | 137 | 96 | 22 | 138 | 51 | 78 |
| UUC | 47 | 11 | 39 | 16 | 89 | 23 | 45 |
| UUG | 82 | 215 | 126 | 15 | 243 | 71 | 107 |
| UUU | 111 | 98 | 107 | 29 | 145 | 26 | 59 |

**Table S14.** Rankings of quality-filtered read numbers for each triplet and amino acid using **Lib-S3** with the mixed anhydride chemistry.

|  | **L-Ala** | **Gly** | **Pro** | **Val** | **Leu** |
| --- | --- | --- | --- | --- | --- |
| AAA | 9 | 35 | 51 | 48 | 47 |
| AAC | 25 | 43 | 57 | 44 | 52 |
| AAG | 32 | 33 | 29 | 13 | 15 |
| AAU | 11 | 23 | 53 | 40 | 58 |
| ACA | 22 | 28 | 37 | 55 | 30 |
| ACC | 50 | 58 | 63 | 62 | 63 |
| ACG | 19 | 21 | 16 | 26 | 20 |
| ACU | 35 | 48 | 46 | 49 | 57 |
| AGA | 13 | 49 | 49 | 57 | 46 |
| AGC | 43 | 56 | 42 | 42 | 35 |
| AGG | 15 | 40 | 25 | 14 | 5 |
| AGU | 10 | 50 | 43 | 29 | 17 |
| AUC | 18 | 3 | 52 | 59 | 61 |
| AUG | 3 | 38 | 27 | 11 | 11 |
| AUU | 21 | 44 | 54 | 56 | 45 |
| CAA | 8 | 24 | 30 | 34 | 36 |
| CAC | 55 | 36 | 61 | 60 | 59 |
| CAG | 4 | 46 | 38 | 51 | 29 |
| CAU | 31 | 51 | 50 | 41 | 38 |
| CCA | 49 | 61 | 45 | 52 | 55 |
| CCC | 63 | 45 | 64 | 64 | 64 |
| CCG | 53 | 34 | 33 | 37 | 50 |
| CCU | 46 | 54 | 60 | 50 | 56 |
| CGA | 48 | 64 | 62 | 63 | 41 |
| CGC | 60 | 63 | 8 | 47 | 53 |
| CGG | 34 | 47 | 40 | 35 | 25 |
| CGU | 57 | 60 | 44 | 53 | 28 |
| CUA | 30 | 20 | 19 | 25 | 12 |
| CUC | 24 | 1 | 14 | 58 | 60 |
| CUG | 26 | 4 | 15 | 20 | 23 |
| CUU | 36 | 39 | 21 | 33 | 40 |
| GAA | 23 | 10 | 55 | 27 | 27 |
| GAC | 47 | 27 | 32 | 43 | 37 |
| GAG | 29 | 5 | 9 | 5 | 1 |
| GAU | 17 | 31 | 59 | 54 | 54 |
| GCA | 51 | 19 | 13 | 17 | 13 |
| GCC | 62 | 30 | 48 | 36 | 49 |
| GCG | 40 | 2 | 4 | 3 | 3 |
| GCU | 42 | 12 | 2 | 1 | 44 |
| GGA | 44 | 14 | 41 | 46 | 33 |
| GGC | 58 | 57 | 24 | 32 | 14 |
| GGG | 27 | 11 | 3 | 4 | 2 |
| GGU | 20 | 59 | 47 | 31 | 24 |
| GUA | 61 | 55 | 11 | 38 | 6 |
| GUC | 64 | 32 | 39 | 45 | 43 |
| GUG | 39 | 16 | 7 | 9 | 4 |
| GUU | 45 | 53 | 34 | 23 | 26 |
| UAA | 6 | 41 | 28 | 21 | 34 |
| UAA | 5 | 13 | 23 | 8 | 22 |
| UAC | 37 | 29 | 31 | 30 | 42 |
| UAG | 1 | 15 | 12 | 2 | 10 |
| UAU | 14 | 22 | 36 | 28 | 32 |
| UCA | 38 | 25 | 18 | 15 | 19 |
| UCC | 59 | 62 | 58 | 61 | 62 |
| UCG | 56 | 52 | 26 | 18 | 31 |
| UCU | 33 | 6 | 1 | 19 | 39 |
| UGA | 41 | 8 | 56 | 16 | 16 |
| UGC | 52 | 9 | 17 | 12 | 21 |
| UGG | 12 | 7 | 5 | 7 | 7 |
| UGU | 28 | 17 | 10 | 22 | 9 |
| UUA | 7 | 26 | 22 | 10 | 18 |
| UUC | 54 | 37 | 35 | 39 | 51 |
| UUG | 2 | 42 | 6 | 6 | 8 |
| UUU | 16 | 18 | 20 | 24 | 48 |

**Table S15.** Selectivity scores (see Sequencing and data processing) for each amino acid transferred with **Lib-S3** in the mixed anhydride chemistry. Top 3 of each amino acid are highlighted in red. Trinucleotides selected for synthesis and kinetic verification are highlighted in yellow. A low score indicates high selectivity.

|  | **L-Ala** | **Gly** | **Pro** | **Leu** | **L&V** |
| --- | --- | --- | --- | --- | --- |
| AAA | 0.050 | 0.226 | 0.367 | 0.329 | 1.000 |
| AAC | 0.128 | 0.242 | 0.348 | 0.308 | 0.768 |
| AAG | 0.356 | 0.371 | 0.312 | 0.140 | 0.298 |
| AAU | 0.063 | 0.142 | 0.402 | 0.457 | 1.126 |
| ACA | 0.147 | 0.194 | 0.274 | 0.211 | 0.977 |
| ACC | 0.203 | 0.244 | 0.270 | 0.270 | 0.731 |
| ACG | 0.229 | 0.259 | 0.186 | 0.244 | 0.821 |
| ACU | 0.175 | 0.257 | 0.243 | 0.320 | 0.822 |
| AGA | 0.065 | 0.297 | 0.297 | 0.274 | 0.928 |
| AGC | 0.246 | 0.346 | 0.239 | 0.191 | 0.546 |
| AGG | 0.179 | 0.678 | 0.338 | 0.053 | 0.238 |
| AGU | 0.072 | 0.505 | 0.406 | 0.129 | 0.447 |
| AUC | 0.103 | 0.016 | 0.369 | 0.462 | 1.644 |
| AUG | 0.034 | 0.731 | 0.429 | 0.139 | 0.324 |
| AUU | 0.106 | 0.250 | 0.325 | 0.257 | 0.849 |
| CAA | 0.065 | 0.222 | 0.294 | 0.375 | 1.129 |
| CAC | 0.255 | 0.153 | 0.290 | 0.278 | 0.783 |
| CAG | 0.024 | 0.377 | 0.292 | 0.209 | 0.909 |
| CAU | 0.172 | 0.319 | 0.311 | 0.220 | 0.598 |
| CCA | 0.230 | 0.303 | 0.207 | 0.266 | 0.690 |
| CCC | 0.266 | 0.176 | 0.271 | 0.271 | 0.744 |
| CCG | 0.344 | 0.197 | 0.190 | 0.318 | 0.725 |
| CCU | 0.209 | 0.255 | 0.291 | 0.267 | 0.663 |
| CGA | 0.209 | 0.299 | 0.287 | 0.173 | 0.598 |
| CGC | 0.351 | 0.375 | 0.036 | 0.298 | 0.763 |
| CGG | 0.231 | 0.351 | 0.284 | 0.160 | 0.496 |
| CGU | 0.308 | 0.330 | 0.222 | 0.131 | 0.503 |
| CUA | 0.395 | 0.233 | 0.218 | 0.128 | 0.536 |
| CUC | 0.180 | 0.006 | 0.098 | 0.619 | 3.026 |
| CUG | 0.419 | 0.048 | 0.205 | 0.354 | 0.956 |
| CUU | 0.271 | 0.300 | 0.142 | 0.310 | 0.760 |
| GAA | 0.193 | 0.076 | 0.632 | 0.235 | 0.614 |
| GAC | 0.338 | 0.170 | 0.208 | 0.248 | 0.755 |
| GAG | 1.450 | 0.114 | 0.225 | 0.021 | 0.140 |
| GAU | 0.086 | 0.168 | 0.378 | 0.335 | 1.009 |
| GCA | 0.823 | 0.202 | 0.130 | 0.130 | 0.361 |
| GCC | 0.380 | 0.154 | 0.271 | 0.278 | 0.607 |
| GCG | 3.333 | 0.040 | 0.083 | 0.061 | 0.130 |
| GCU | 0.712 | 0.135 | 0.020 | 0.772 | 0.804 |
| GGA | 0.328 | 0.085 | 0.299 | 0.228 | 0.798 |
| GGC | 0.457 | 0.445 | 0.149 | 0.082 | 0.331 |
| GGG | 1.350 | 0.306 | 0.068 | 0.044 | 0.146 |
| GGU | 0.124 | 0.484 | 0.351 | 0.153 | 0.437 |
| GUA | 0.555 | 0.474 | 0.069 | 0.036 | 0.346 |
| GUC | 0.403 | 0.168 | 0.212 | 0.239 | 0.652 |
| GUG | 1.083 | 0.271 | 0.103 | 0.056 | 0.210 |
| GUU | 0.331 | 0.414 | 0.231 | 0.168 | 0.371 |
| UAA | 0.048 | 0.461 | 0.275 | 0.354 | 0.733 |
| UAA | 0.076 | 0.224 | 0.479 | 0.449 | 0.732 |
| UAC | 0.280 | 0.207 | 0.225 | 0.331 | 0.742 |
| UAG | 0.026 | 0.600 | 0.429 | 0.333 | 0.429 |
| UAU | 0.119 | 0.200 | 0.375 | 0.320 | 0.833 |
| UCA | 0.494 | 0.278 | 0.186 | 0.198 | 0.420 |
| UCC | 0.243 | 0.258 | 0.238 | 0.258 | 0.687 |
| UCG | 0.441 | 0.397 | 0.166 | 0.204 | 0.366 |
| UCU | 0.508 | 0.065 | 0.010 | 0.661 | 1.450 |
| UGA | 0.427 | 0.062 | 0.691 | 0.132 | 0.305 |
| UGC | 0.881 | 0.088 | 0.181 | 0.233 | 0.423 |
| UGG | 0.462 | 0.226 | 0.152 | 0.226 | 0.583 |
| UGU | 0.483 | 0.246 | 0.132 | 0.117 | 0.564 |
| UUA | 0.092 | 0.456 | 0.361 | 0.277 | 0.509 |
| UUC | 0.333 | 0.207 | 0.193 | 0.309 | 0.714 |
| UUG | 0.032 | 1.909 | 0.103 | 0.143 | 0.280 |
| UUU | 0.145 | 0.167 | 0.189 | 0.615 | 1.333 |

**Table S16.** Quality-filtered read numbers from the sequencing results of phosphoramidate-ester **Lib-S3**.

|  | **Blank** | **L-Ala** | **D-Ala** | **Gly** | **Pro** | **Leu** | **Val** |
| --- | --- | --- | --- | --- | --- | --- | --- |
| AAA | 111 | 1735 | 1962 | 828 | 1862 | 2797 | 1623 |
| AAC | 40 | 2274 | 2805 | 1078 | 2535 | 3661 | 2007 |
| AAG | 65 | 4772 | 5930 | 2188 | 5251 | 7443 | 4093 |
| AAU | 77 | 2124 | 2314 | 1060 | 2383 | 3128 | 1780 |
| ACA | 149 | 1582 | 1703 | 764 | 1714 | 2387 | 1333 |
| ACC | 90 | 1133 | 1508 | 609 | 1355 | 2086 | 1002 |
| ACG | 16 | 1842 | 2282 | 947 | 2105 | 3129 | 1532 |
| ACU | 91 | 1816 | 2270 | 939 | 2201 | 3085 | 1573 |
| AGA | 179 | 2233 | 2316 | 1019 | 2355 | 3789 | 2082 |
| AGC | 96 | 5595 | 6974 | 2820 | 6190 | 9212 | 4879 |
| AGG | 73 | 3276 | 3712 | 1537 | 3465 | 5842 | 2905 |
| AGU | 88 | 6560 | 7770 | 3198 | 6817 | 11166 | 5661 |
| AUC | 79 | 1660 | 1862 | 707 | 1775 | 2601 | 1398 |
| AUG | 110 | 1909 | 2366 | 815 | 2167 | 3128 | 1567 |
| AUU | 57 | 3174 | 3800 | 1527 | 3435 | 5026 | 2661 |
| CAA | 70 | 2026 | 2358 | 956 | 2247 | 3309 | 1786 |
| CAC | 82 | 1521 | 1612 | 759 | 1629 | 2289 | 1234 |
| CAG | 176 | 1705 | 1964 | 845 | 1961 | 2628 | 1341 |
| CAU | 54 | 4388 | 5487 | 1981 | 4926 | 7473 | 3850 |
| CCA | 58 | 1812 | 2148 | 927 | 2053 | 2833 | 1513 |
| CCC | 59 | 1173 | 1294 | 616 | 1322 | 1817 | 985 |
| CCG | 133 | 1257 | 1238 | 716 | 1426 | 1978 | 1041 |
| CCU | 18 | 1289 | 1493 | 670 | 1489 | 2005 | 1012 |
| CGA | 87 | 1385 | 1589 | 688 | 1690 | 2383 | 1203 |
| CGC | 86 | 1305 | 1496 | 669 | 1483 | 2002 | 1134 |
| CGG | 81 | 2440 | 3072 | 1224 | 2790 | 4361 | 2064 |
| CGU | 12 | 1255 | 1296 | 642 | 1349 | 2151 | 1050 |
| CUA | 35 | 2209 | 2720 | 1079 | 2446 | 4020 | 1918 |
| CUC | 64 | 1467 | 1637 | 730 | 1635 | 2491 | 1261 |
| CUG | 121 | 1220 | 1527 | 606 | 1392 | 2308 | 1039 |
| CUU | 36 | 2107 | 2892 | 1157 | 2423 | 3996 | 1772 |
| GAA | 55 | 1868 | 2389 | 1086 | 2205 | 3471 | 1654 |
| GAC | 23 | 2712 | 3128 | 1230 | 3032 | 4510 | 2307 |
| GAG | 234 | 5801 | 5363 | 2637 | 6103 | 9850 | 4809 |
| GAU | 196 | 3997 | 4314 | 1783 | 4201 | 6412 | 3423 |
| GCA | 150 | 1935 | 2361 | 964 | 2167 | 3336 | 1777 |
| GCC | 30 | 2210 | 2729 | 1079 | 2508 | 3897 | 1772 |
| GCG | 25 | 2618 | 3133 | 1316 | 3149 | 4634 | 2269 |
| GCU | 19 | 2663 | 3231 | 1251 | 3374 | 4805 | 2442 |
| GGA | 56 | 727 | 562 | 361 | 722 | 1101 | 587 |
| GGC | 28 | 4200 | 4633 | 2023 | 4615 | 7453 | 3658 |
| GGG | 10 | 2267 | 1828 | 1011 | 2297 | 3444 | 1783 |
| GGU | 23 | 3432 | 3478 | 1713 | 3872 | 6071 | 3152 |
| GUA | 48 | 4067 | 4845 | 1868 | 4245 | 6537 | 3484 |
| GUC | 28 | 3159 | 4268 | 1520 | 3584 | 5484 | 2580 |
| GUG | 12 | 5452 | 6904 | 2606 | 6030 | 9609 | 4785 |
| GUU | 30 | 4719 | 5912 | 2373 | 5259 | 8240 | 4328 |
| UAA | 75 | 1884 | 2157 | 915 | 2077 | 2893 | 1555 |
| UAA | 116 | 1782 | 2311 | 945 | 2058 | 3073 | 1474 |
| UAC | 40 | 3647 | 4606 | 1778 | 3953 | 6151 | 3171 |
| UAG | 69 | 2449 | 2798 | 1124 | 2662 | 3792 | 2063 |
| UAU | 93 | 1694 | 1924 | 814 | 1805 | 2614 | 1409 |
| UCA | 80 | 1384 | 1785 | 732 | 1723 | 2474 | 1141 |
| UCC | 36 | 1952 | 2544 | 1060 | 2233 | 3525 | 1649 |
| UCG | 72 | 1871 | 2224 | 963 | 2054 | 3084 | 1604 |
| UCU | 52 | 2056 | 2270 | 996 | 2119 | 3368 | 1813 |
| UGA | 62 | 745 | 788 | 363 | 835 | 1375 | 639 |
| UGC | 28 | 3608 | 4268 | 1745 | 4009 | 6605 | 3230 |
| UGG | 42 | 3824 | 4302 | 1763 | 3889 | 6764 | 3338 |
| UGU | 83 | 1993 | 2125 | 953 | 2076 | 2991 | 1663 |
| UUA | 69 | 2130 | 2563 | 1076 | 2416 | 3713 | 1880 |
| UUC | 34 | 3494 | 4310 | 1739 | 3599 | 5717 | 2837 |
| UUG | 52 | 2562 | 2870 | 1271 | 2645 | 4257 | 2171 |
| UUU | 23 | 2712 | 3128 | 1230 | 3032 | 4510 | 2307 |

**Table S17.** Rankings of quality-filtered read numbers for each triplet and amino acid in phosphoramidate-ester formation with **Lib-S3**.

|  | **L-Ala** | **D-Ala** | **Gly** | **Pro** | **Leu** | **Val** |
| --- | --- | --- | --- | --- | --- | --- |
| AAA | 10 | 47 | 47 | 47 | 48 | 47 |
| AAC | 45 | 26 | 26 | 30 | 26 | 31 |
| AAG | 31 | 5 | 4 | 6 | 6 | 8 |
| AAU | 24 | 32 | 37 | 32 | 31 | 39 |
| ACA | 6 | 51 | 52 | 50 | 52 | 53 |
| ACC | 15 | 62 | 57 | 61 | 60 | 58 |
| ACG | 61 | 43 | 39 | 41 | 41 | 38 |
| ACU | 14 | 44 | 40 | 43 | 37 | 41 |
| AGA | 3 | 28 | 36 | 34 | 32 | 29 |
| AGC | 12 | 3 | 2 | 2 | 2 | 4 |
| AGG | 26 | 17 | 18 | 17 | 18 | 16 |
| AGU | 16 | 1 | 1 | 1 | 1 | 1 |
| AUC | 23 | 50 | 49 | 55 | 50 | 50 |
| AUG | 11 | 39 | 33 | 48 | 38 | 40 |
| AUU | 36 | 18 | 17 | 18 | 19 | 19 |
| CAA | 28 | 35 | 35 | 39 | 34 | 37 |
| CAC | 20 | 52 | 54 | 51 | 55 | 56 |
| CAG | 4 | 48 | 46 | 46 | 47 | 48 |
| CAU | 39 | 7 | 6 | 9 | 7 | 6 |
| CCA | 35 | 45 | 44 | 44 | 46 | 46 |
| CCC | 34 | 61 | 61 | 60 | 62 | 62 |
| CCG | 7 | 58 | 62 | 54 | 58 | 61 |
| CCU | 60 | 57 | 59 | 57 | 56 | 59 |
| CGA | 17 | 54 | 55 | 56 | 53 | 54 |
| CGC | 18 | 56 | 58 | 58 | 57 | 60 |
| CGG | 21 | 25 | 23 | 24 | 23 | 23 |
| CGU | 62 | 59 | 60 | 59 | 61 | 57 |
| CUA | 49 | 30 | 29 | 28 | 28 | 25 |
| CUC | 32 | 53 | 53 | 53 | 54 | 51 |
| CUG | 8 | 60 | 56 | 62 | 59 | 55 |
| CUU | 47 | 33 | 24 | 25 | 29 | 26 |
| GAA | 38 | 42 | 32 | 27 | 36 | 33 |
| GAC | 43 | 9 | 16 | 7 | 11 | 9 |
| GAG | 57 | 20 | 22 | 23 | 22 | 22 |
| GAU | 1 | 2 | 7 | 3 | 3 | 2 |
| GCA | 2 | 11 | 11 | 11 | 10 | 13 |
| GCC | 5 | 38 | 34 | 37 | 39 | 36 |
| GCG | 51 | 29 | 28 | 29 | 27 | 27 |
| GCU | 56 | 22 | 21 | 20 | 21 | 21 |
| GGA | 59 | 21 | 20 | 22 | 20 | 20 |
| GGC | 37 | 64 | 64 | 64 | 64 | 64 |
| GGG | 53 | 8 | 9 | 8 | 8 | 7 |
| GGU | 64 | 27 | 50 | 35 | 33 | 34 |
| GUA | 58 | 16 | 19 | 16 | 15 | 15 |
| GUC | 42 | 10 | 8 | 10 | 9 | 12 |
| GUG | 54 | 19 | 14 | 19 | 17 | 18 |
| GUU | 63 | 4 | 3 | 4 | 4 | 3 |
| UAA | 52 | 6 | 5 | 5 | 5 | 5 |
| UAA | 25 | 40 | 43 | 45 | 42 | 45 |
| UAC | 9 | 46 | 38 | 42 | 44 | 43 |
| UAG | 46 | 13 | 10 | 12 | 13 | 14 |
| UAU | 29 | 24 | 27 | 26 | 24 | 28 |
| UCA | 13 | 49 | 48 | 49 | 49 | 49 |
| UCC | 22 | 55 | 51 | 52 | 51 | 52 |
| UCG | 48 | 37 | 31 | 33 | 35 | 32 |
| UCU | 27 | 41 | 42 | 38 | 45 | 42 |
| UGA | 40 | 34 | 41 | 36 | 40 | 35 |
| UGC | 33 | 63 | 63 | 63 | 63 | 63 |
| UGG | 55 | 14 | 15 | 14 | 12 | 11 |
| UGU | 44 | 12 | 13 | 13 | 14 | 10 |
| UUA | 19 | 36 | 45 | 40 | 43 | 44 |
| UUC | 30 | 31 | 30 | 31 | 30 | 30 |
| UUG | 50 | 15 | 12 | 15 | 16 | 17 |
| UUU | 41 | 23 | 25 | 21 | 25 | 24 |

**Table S18.** Kinetic data of mixed anhydride aminoacyl-transfer with **SM-11**, relating to **Fig. 3A**. The yield in bold indicates the highest yield among all the amino acid transfers. Shading line indicates a G:U wobble basepair.

|  | **Donor** | **Acceptor** | | | **Observed yield** | **Corrected yield** | **Mixed anhyd.** | **Diol ester** | **Peak time** | **k transfer** | **k hydrolysis** |
| --- | --- | --- | --- | --- | --- | --- | --- | --- | --- | --- | --- |
|  |  | **Stem** | **Overhang** | |  |  | **half-life** /h | **half-life** /h | min | /min^-1^ | /min^-1^ |
| **SM-11** | **L-Ala-pCUGCG** | CGCAG | | UUCCA | **49%** | **61%** | 0.71 | 7.5 | 63 | 0.0591 | 0.00154 |
|  | L-Ala-pCUGCG | CGUAG | | UUCCA | 28% | 34% | / | 14 | 135 | 0.0268 | 0.00081 |
|  | L-Ala-pCUGCG |  | | -- | -- | -- | 1.23 | -- | -- | -- | -- |
|  | D-Ala-pCUGCG | CGCAG | | UUCCA | 8% | 15% | 0.46 | 20 | 207 | 0.0170 | 0.00057 |
|  | D-Ala-pCUGCG |  | | -- | -- | -- | 1.52 | -- | -- | -- | -- |
|  | Gly-pCUGCG | CGCAG | | UUCCA | 0% | 0% | / | -- | -- | -- | -- |
|  | Gly-pCUGCG |  | | -- | -- | -- | / | -- | -- | -- | -- |
|  | L-Val-pCUGCG | CGCAG | | UUCCA | 14% | 24% | 1.11 | > 24 | > 200 | 0.0170 | < 0.00050 |
|  | L-Val-pCUGCG |  | | -- | -- | -- | 1.48 | -- | -- | -- | -- |
|  | L-Leu-pCUGCG | CGCAG | | UUCCA | 28% | 45% | 0.40 | 7.7 | 65 | 0.0573 | 0.00151 |
|  | L-Leu-pCUGCG |  | | -- | -- | -- | 0.90 | -- | -- | -- | -- |
|  | L-Pro-pCUGCG | CGCAG | | UUCCA | 29% | 36% | 0.31 | 2.2 | 51 | 0.0494 | 0.00523 |
|  | L-Pro-pCUGCG |  | | -- | -- | -- | 1.50 | -- | -- | -- | -- |

/, not measured.

**Table S19.** Kinetic data of mixed anhydride aminoacyl-transfer with **SM-13**, relating to **Fig. 3B**.

|  | **Donor** | **Acceptor** | | **Observed**  **yield** | **Corrected yield** | **Mixed anhyd.** | **Diol ester** | **Peak time** | **k transfer** | **k hydrolysis** |
| --- | --- | --- | --- | --- | --- | --- | --- | --- | --- | --- |
|  |  | **Stem** | **Overhang** |  |  | **half-life** /h | **half-life** /h | min | /min^-1^ | /min^-1^ |
| **SM-13** | L-Ala-pUACCG | CGGUA | UUCCA | 14% | 18% | 0.90 | 4.8 | 36 | 0.1084 | 0.00243 |
|  | L-Ala-pUACCG |  | -- | -- | -- | 1.14 | -- | -- | -- | -- |
|  | D-Ala-pUACCG | CGGUA | UUCCA | 15% | 17% | 0.39 | 6.4 | 79 | 0.0413 | 0.00179 |
|  | D-Ala-pUACCG |  | -- | -- | -- | 0.41 | -- | -- | -- | -- |
|  | Gly-pUACCG | CGGUA | UUCCA | 2% | 2% | 0.73 | -- | -- | -- | -- |
|  | Gly-pUACCG |  | -- | -- | -- | 1.27 | -- | -- | -- | -- |
|  | L-Val-pUACCG | CGGUA | UUCCA | 0% | 0% | 0.70 | -- | -- | -- | -- |
|  | L-Val-pUACCG |  | -- | -- | -- | 0.72 | -- | -- | -- | -- |
|  | **L-Leu-pUACCG** | CGGUA | UUCCA | **34%** | **50%** | 0.76 | 18 | 166 | 0.0220 | 0.00063 |
|  | L-Leu-pUACCG |  | -- | -- | -- | 0.99 | -- | -- | -- | -- |
|  | L-Pro-pUACCG | CGGUA | UUCCA | 22% | 37% | 0.52 | 4.2 | 74 | 0.0379 | 0.00277 |
|  | L-Pro-pUACCG |  | -- | -- | -- | 1.30 | -- | -- | -- | -- |

**Table S20.** Kinetic data of mixed anhydride aminoacyl-transfer with **SM-14**, relating to **Fig. 3C**.

|  | **Donor** | **Acceptor** | | | **Observed yield** | **Corrected yield** | **Mixed anhyd.** | **Diol ester** | **Peak time** | **k transfer** | **k hydrolysis** |
| --- | --- | --- | --- | --- | --- | --- | --- | --- | --- | --- | --- |
|  |  | **Stem** | **Overhang** | |  |  | **half-life** /h | **half-life** /h | min | /min^-1^ | /min^-1^ |
| **SM-14** | L-Ala-pCUCCG | CGGAG | | UUCCA | 31% | 33% | 0.43 | 7.0 | 97 | 0.0326 | 0.00165 |
|  | L-Ala-pCUCCG |  | | -- | -- | -- | 1.36 | -- | -- | -- | -- |
|  | D-Ala-pCUCCG | CGGAG | | UUCCA | 14% | 15% | 1.01 | 7.7 | 137 | 0.0207 | 0.00150 |
|  | D-Ala-pCUCCG |  | | -- | -- | -- | 1.31 | -- | -- | -- | -- |
|  | Gly-pCUCCG | CGGAG | | UUCCA | 7% | 8% | / | 17 | 54 | 0.0912 | 0.00068 |
|  | Gly-pCUCCG |  | | -- | -- | -- | 2.06 | -- | -- | -- | -- |
|  | **L-Val-pCUCCG** | CGGAG | | UUCCA | **6%** | **32%** | 0.90 | > 24 | > 200 | 0.0130 | < 0.00050 |
|  | L-Val-pCUCCG |  | | -- | -- | -- | 1.58 | -- | -- | -- | -- |
|  | **L-Leu-pCUCCG** | CGGAG | | UUCCA | **29%** | **51%** | 0.65 | 16 | 135 | 0.0278 | 0.00072 |
|  | L-Leu-pCUCCG |  | | -- | -- | -- | 1.44 | -- | -- | -- | -- |
|  | L-Pro-pCUCCG | CGGAG | | UUCCA | 21% | 30% | 0.68 | 2.6 | 84 | 0.0249 | 0.00445 |
|  | L-Pro-pCUCCG |  | | -- | -- | -- | 1.89 | -- | -- | -- | -- |

**Table S21.** Kinetic data of mixed anhydride aminoacyl-transfer with **SM-15**, relating to **Fig. 3D**.

|  | **Donor** | **Acceptor** | | | **Observed yield** | **Corrected yield** | **Mixed anhyd.** | **Diol ester** | **Peak time** | **k transfer** | **k hydrolysis** |
| --- | --- | --- | --- | --- | --- | --- | --- | --- | --- | --- | --- |
|  |  | **Stem** | **Overhang** | |  |  | **half-life** /h | **half-life** /h | min | /min^-1^ | /min^-1^ |
| **SM-15** | L-Ala-pAGACG | CGUCU | | UUCCA | 22% | 23% | 0.47 | 6.5 | 123 | 0.0223 | 0.00177 |
|  | L-Ala-pAGACG |  | | -- | -- | -- | 1.43 | -- | -- | -- | -- |
|  | D-Ala-pAGACG | CGUCU | | UUCCA | 15% | 17% | 0.74 | 3.2 | 145 | 0.0119 | 0.00356 |
|  | D-Ala-pAGACG |  | | -- | -- | -- | 1.65 | -- | -- | -- | -- |
|  | Gly-pAGACG | CGUCU | | UUCCA | 0% | 0% | 1.05 | -- | -- | -- | -- |
|  | Gly-pAGACG |  | | -- | -- | -- | 1.00 | -- | -- | -- | -- |
|  | L-Val-pAGACG | CGUCU | | UUCCA | 4% | 7% | 0.74 | > 24 | > 200 | 0.0174 | < 0.00050 |
|  | L-Val-pAGACG |  | | -- | -- | -- | 1.48 | -- | -- | -- | -- |
|  | L-Leu-pAGACG | CGUCU | | UUCCA | 14% | 25% | 0.44 | > 24 | > 200 | 0.0218 | < 0.00050 |
|  | L-Leu-pAGACG |  | | -- | -- | -- | 1.48 | -- | -- | -- | -- |
|  | **L-Pro-pAGACG** | CGUCU | | UUCCA | **24%** | **32%** | 0.42 | 3.1 | 65 | 0.0405 | 0.00375 |
|  | L-Pro-pAGACG |  | | -- | -- | -- | 0.96 | -- | -- | -- | -- |

**Table S22.** Kinetic data of mixed anhydride aminoacyl-transfer with **SM-12**, relating to **Fig. 3E**.

|  | **Donor** | **Acceptor** | | **Observed yield** | **Corrected yield** | **Mixed anhyd.** | **Diol ester** | **Peak time** | **k transfer** | **k hydrolysis** |
| --- | --- | --- | --- | --- | --- | --- | --- | --- | --- | --- |
|  |  | **Stem** | **Overhang** |  |  | **half-life** /h | **half-life** /h | min | /min^-1^ | /min^-1^ |
| **SM-12** | L-Ala-pGAGCG | CGCUC | UUCCA | 18% | 21% | 0.48 | 27 | 141 | 0.0308 | 0.00043 |
|  | L-Ala-pGAGCG |  | -- | -- | -- | 1.52 | -- | -- | -- | -- |
|  | D-Ala-pGAGCG | CGCUC | UUCCA | 11% | 13% | 0.93 | 24 | 150 | 0.0273 | 0.00048 |
|  | D-Ala-pGAGCG |  | -- | -- | -- | 1.86 | -- | -- | -- | -- |
|  | **Gly-pGAGCG** | CGCUC | UUCCA | **9%** | **12%** | 0.32 | > 24 | 74 | 0.0732 | < 0.00050 |
|  | Gly-pGAGCG |  | -- | -- | -- | 1.67 | -- | -- | -- | -- |
|  | L-Val-pGAGCG | CGCUC | UUCCA | 1% | 7% | 0.61 | > 24 | > 200 | 0.0394 | < 0.00050 |
|  | L-Val-pGAGCG |  | -- | -- | -- | 0.65 | -- | -- | -- | -- |
|  | L-Leu-pGAGCG | CGCUC | UUCCA | 10% | 23% | 0.58 | > 24 | 190 | 0.0228 | < 0.00050 |
|  | L-Leu-pGAGCG |  | -- | -- | -- | 1.67 | -- | -- | -- | -- |
|  | L-Pro-pGAGCG | CGCUC | UUCCA | 12% | 16% | 0.63 | 6.1 | 98 | 0.0302 | 0.00189 |
|  | L-Pro-pGAGCG |  | -- | -- | -- | 2.06 | -- | -- | -- | -- |

**Table S23.** Kinetic data of mixed anhydride aminoacyl-transfer with **SM-16** and **SM-17**, relating to **Fig. 3F** and **3G**.

|  | **Donor** | **Acceptor** | | | **Observed yield** | **Corrected yield** | **Mixed anhyd.** | **Diol ester** | **Peak time** | **k transfer** | **k hydrolysis** |
| --- | --- | --- | --- | --- | --- | --- | --- | --- | --- | --- | --- |
|  |  | **Stem** | **Overhang** | |  |  | **half-life** /h | **half-life** /h | min | /min^-1^ | /min^-1^ |
| **SM-16** | **L-Ala-pCUGC** | CGCAG | | UUCCA | **48%** | **62%** | / | 6.6 | 63 | 0.0573 | 0.00175 |
|  | L-Ala-pCUGC |  | | -- | -- | -- | 0.43 | -- | -- | -- | -- |
|  | L-Pro-pCUGC | CGCAG | | UUCCA | 26% | 35% | / | 1.8 | 50 | 0.0456 | 0.00631 |
|  | L-Pro-pCUGC |  | | -- | -- | -- | 1.12 | -- | -- | -- | -- |
|  | L-Leu-pCUGC | CGCAG | | UUCCA | 27% | 46% | 0.41 | 14 | 97 | 0.0408 | 0.00083 |
|  | L-Leu-pCUGC |  | | -- | -- | -- | 0.56 | -- | -- | -- | -- |

| **SM-17 Donor** | **Acceptor** | | | **Temp.** | **Observed yield** | **Corrected yield** | **Mixed anhyd.** | **Diol ester** | **Peak time** | **k transfer** | **k hydrolysis** | **Donor eq.** |
| --- | --- | --- | --- | --- | --- | --- | --- | --- | --- | --- | --- | --- |
|  | **Stem** | **Overhang** | | ℃ |  |  | **half-life** /h | **half-life** /h | min | /min^-1^ | /min^-1^ |  |
| L-Ala-pCUG | CGCAG | | UUCCA | 5 | 37% | 44% | / | 13 | 88 | 0.0458 | 0.00089 | 1 eq. |
| **L-Ala-pCUG** | CGCAG | | UUCCA | **10** | **38%** | **46%** | / | 8.3 | 115 | 0.0272 | 0.00140 | 1 eq. |
| L-Ala-pCUG | CGCAG | | UUCCA | 16 | 23% | 27% | / | 6.1 | 71 | 0.0470 | 0.00190 | 1 eq. |
| L-Ala-pCUG | CGCAG | | UUCCA | 16 | 43% | 52% | / | 4.8 | 74 | 0.0404 | 0.00239 | 3 eq. |
| L-Ala-pCUG |  | | -- | 10 | -- | -- | 0.63 | -- | -- | -- | -- |  |
| L-Pro-pCUG | CGCAG | | UUCCA | 10 | 17% | 20% | / | 3.1 | 88 | 0.0255 | 0.00373 | 1 eq. |
| L-Pro-pCUG | CGCAG | | UUCCA | 16 | 13% | 14% | / | 2.7 | 75 | 0.0304 | 0.00423 | 1 eq. |
| L-Pro-pCUG |  | | -- | 10 | -- | -- | 1.20 | -- | -- | -- | -- |  |
| L-Leu-pCUG | CGCAG | | UUCCA | 10 | 19% | 23% | 0.69 | 26 | 172 | 0.0234 | 0.00045 | 1 eq. |
| L-Leu-pCUG | CGCAG | | UUCCA | 16 | 11% | 12% | 0.44 | 17 | 103 | 0.0398 | 0.00069 | 1 eq. |
| L-Leu-pCUG |  | | -- | 10 | -- | -- | 0.66 | -- | -- | -- | -- |  |

**Table S24.** Kinetic data of mixed anhydride aminoacyl-transfer with **SM-18** (triplet sequence for best L-Ala transfer yield), and **SM-19**.

|  | **Donor** | **Acceptor** | | | **Observed yield** | **Corrected yield** | **Mixed anhyd.** | **Diol ester** | **Peak time** | **k transfer** | **k hydrolysis** |
| --- | --- | --- | --- | --- | --- | --- | --- | --- | --- | --- | --- |
|  |  | **Stem** | **Overhang** | |  |  | **half-life** /h | **half-life** /h | min | /min^-1^ | /min^-1^ |
| **SM-18** | L-Ala-pCUAAC | GUUAC | | UUCCA | 40% | 51% | 0.37 | 4.0 | 42 | 0.0829 | 0.00288 |
|  | L-Ala-pCUAAC | -- | | -- | -- | -- | 0.63 | -- | -- | -- | -- |
| **SM-19** | L-Leu-pACUUG | CAAGU | | UUCCA | 15% | 25% | 1.05 | 8.0 | > 200 | 0.0105 | 0.0014 |
|  | L-Leu-pACUUG | -- | | -- | -- | -- | 1.19 | -- | -- | -- | -- |

**Table S25**. Overview of duplexes for which simulations were conducted in **Fig. 4**, **Fig. S8** and **S9**. Energy landscapes and characteristic structures for these systems are publicly available from the zenodo repository.

|  | **Donor** | **Acceptor** | | **Corrected** | **k transfer** |
| --- | --- | --- | --- | --- | --- |
|  |  | **Stem** | **Overhang** | **yield** | /min^-1^ |
| **SM-18** | L-Ala-pCUAAC | 5’-GUUAG | UUCCA-3’ | 51% | 0.0829 |
| **SM-4** | L-Ala-pAGGCA | 5’-UGCCU | UUCCA-3’ | 30% | 0.0501 |
| **SM-6** | L-Leu-pCGCAA | 5’-UUGCG | UUCCA-3’ | 60% | 0.0257 |
| **SM-19** | L-Leu-pACUUG | 5’-CAAGU | UUCCA-3’ | 25% | 0.0105 |
| **SM-11** | L-Ala-pCUGCG | 5’-CGCAG | UUCCA-3’ | 61% | 0.0591 |
| **SM-11** | D-Ala-pCUGCG | 5’-CGCAG | UUCCA-3’ | 15% | 0.0170 |
| **SM-11** | L-Leu-pCUGCG | 5’-CGCAG | UUCCA-3’ | 45% | 0.0573 |
| **SM-11** | L-Val-PCUGCG | 5’-CGCAG | UUCCA-3’ | 24% | 0.0170 |
| **SM-13** | L-Ala-pUACCG | 5’-CGGUA | UUCCA-3’ | 18% | 0.1084 |
| **SM-13** | D-Ala-pUACCG | 5’-CGGUA | UUCCA-3’ | 17% | 0.0413 |
| **SM-13** | L-Leu-pUACCG | 5’-CGGUA | UUCCA-3’ | 50% | 0.0220 |
| **SM-13** | L-Val-pUACCG | 5’-CGGUA | UUCCA-3’ | 0% | -- |
